# Elucidation of the molecular interactions that enable stable assembly and structural diversity in multicomponent immune receptors

**DOI:** 10.1101/2020.12.22.424036

**Authors:** Lam-Kiu Fong, Matthew J. Chalkley, Sophia K. Tan, Michael Grabe, William F. DeGrado

## Abstract

Multi-component immune receptors are essential complexes in which distinct ligand-recognition and signaling subunits are held together by interactions between acidic and basic residues of their transmembrane helices. A 2-to-1 acidic to basic motif in the transmembrane domains of the subunits is necessary and sufficient to assemble these receptor complexes. Here, we study a prototype for these receptors, a DAP12-NKG2C 2:1 heterotrimeric complex, in which the two DAP12 subunits each contribute a single transmembrane Asp residue, and the NKG2C subunit contributes a Lys to form the complex. DAP12 can also associate with 20 other subunits using a similar motif. Here we use molecular dynamics simulations to understand the basis for the high affinity and diversity of interactions in this group of receptors. Simulations of the transmembrane helices with differing protonation states of the Asp-Asp-Lys triad identified a structurally stable interaction in which a singly protonated Asp-Asp pair forms a hydrogen-bonded carboxyl-carboxylate clamp that clasps onto a charged Lys sidechain. This polar motif was also supported by density functional theory and a Protein Data Bank-wide search. In contrast, the helices are dynamic at sites distal to the stable carboxyl-carboxylate clamp motif. Such a locally stable but globally dynamic structure is well-suited to accommodate the sequence and structural variations in the transmembrane helices of multi-component receptors, which mix and match subunits to create combinatorial functional diversity from a limited number of subunits. It also supports a signaling mechanism based on multi-subunit clustering rather than propagation of rigid conformational changes through the membrane.

**SIGNIFICANCE:** Receptors that separate ligand recognition and intracellular signaling into separate protein subunits are ubiquitous in immunity. These subunits mix and match to create combinatorial functional diversity. The transmembrane domains of these receptors assemble through the interaction between two acidic and one basic residue on different helices. Using computational methods to study the DAP12-NKG2C receptor complex we identified a polar motif in which a singly protonated Asp-Asp pair forms a carboxyl-carboxylate clamp that clasps a charged Lys sidechain in the membrane. This local interaction allows dynamic variations in other regions of the helices that tolerate sequence diversity of the interacting subunits in this class of receptors, which signal through multi-subunit clustering rather than propagation of rigid conformational changes through the membrane.

## INTRODUCTION

Immune cells often employ multi-component protein receptors that separate specific extracellular ligand recognition and intracellular downstream signaling functions into distinct subunits (1–3). Assembly of these single-pass protein subunits through helix-helix interactions of their transmembrane domains is required for receptor function. The ligand-binding subunit exhibits an elaborate extracellular domain that is adapted for ligand specificity, while the signaling subunit typically contains a pair of evolutionarily conserved intracellular signaling elements, immunoreceptor tyrosine-based activation motifs (ITAMs), that couple to downstream phosphorylation and signaling pathways (4). The details of receptor subunit assembly have not been fully worked out, obscuring the mechanism of signal transduction across the membrane.

The interaction between a pair of acidic aspartate residues on the signaling dimer and a basic residue (lysine or arginine) on the ligand-binding coreceptor are required for complex assembly within the otherwise nonpolar membrane (5–7). This 2-to-1 acidic to basic polar association motif is ubiquitous in multi-component receptor families including Fc receptors (8), T-cell receptor-CD3 complexes (9–11) and the more than 20 receptors across most immune cell types that associate with the signaling subunit DNAX-activation protein 12 (DAP12), a disulfide-linked homodimer with a minimal extracellular region (2, 12–14). DAP12-coreceptor complexes are noteworthy because malfunctions of DAP12 or its coreceptors have been implicated in an array of diseases (15–19). Notably, point mutations of DAP12-associated coreceptor TREM2 have recently been linked to Alzheimer’s disease (20, 21). Additionally, DAP12 must be able to assemble with coreceptors that are as diverse in sequence as they are in ligand-recognition capabilities (22). A molecular level understanding of what enables this promiscuous, yet specific association with so many ligand-binding coreceptors is limited. Therefore, DAP12-coreceptor association serves as an ideal model system for studying polar residue-induced assembly of 2-to-1 multi-component receptors.

The sequence diversity of DAP12-associated receptors suggests that interaction stability of the polar triad association motif (Asp-Asp-Lys) is the primary driver of proper subunit assembly and therefore dictates signaling function in the receptor complex. This motif has the potential to exist in four protonation states: a fully ionized triad with an overall −1 charge (designated D^−^D^−^K^+^), a neutral triad in which the lysine and either of the two aspartates are ionized (D^−^D^0^K^+^), a second neutral state in which each residue is non-ionized (D^0^D^0^K^0^), and the positively-charged state due to ionization of the lysine (D^0^D^0^K^+^). The anionic D^−^D^−^K^+^ state has been widely accepted to drive complex assembly, but evidence for this protonation state is based solely on short coarse-grained simulations in which resolution is limited or relatively short (<60 ns) atomistic molecular dynamics simulation where the time scale prevented observation of conformational differences between protonation states (23–28). Given that the D^−^D^−^K^+^ state would result in an energetically unfavorable net charge in the center of the nonpolar bilayer, we felt that a re-examination of the ionization using multiple computational methods was warranted. We were particularly interested in comparing the fully charged state with the D^−^D^0^K^+^ and D^0^D^0^K^0^ states, which do not require burial of a charge near the center of the bilayer. The D^−^D^0^K^+^ appeared to be particularly favorable, because aspartates on transmembrane peptides have elevated p*K*_a_s that assist helix association, and such an elevation in p*K*_a_ should allow facile protonation of one of the two carboxylate sidechains (29–35). Moreover, there is an extensive literature on the role of carboxyl-carboxylate pairs (in which one of the two carboxylate groups is protonated and the other ionized) in stabilizing protein structures (36–39). Consequently, we hypothesized that the Asp-Asp-Lys interaction in these receptor complexes is stabilized by protonation that results in an overall neutral charge of the triad. We further wanted to understand how the presence and formation of this polar residue network dictates the packing interface of the receptor helices and provide insights into how this conserved signaling subunit is able to stably engage with so many diverse coreceptors.

Here, we used microsecond-long fully atomistic molecular dynamics (MD) simulations to probe the conformational dynamics as a function of polar residue protonation state of the only known structure of DAP12 with a coreceptor, DAP12-NKG2C (PDB id: 2L35), both at the polar residue level and at the helix-helix level. The results suggest that effective assembly of multi-subunit receptors is conformationally static at the local polar residue level, but conformationally dynamic at the level of the tertiary structure, thereby allowing a single conserved signaling module, DAP12, to be simultaneously specific and promiscuous for lysine containing coreceptors. This favorable polar arrangement is only possible with protonation of one of the aspartic acids on the DAP12 signaling subunit. Protonation ensures charge compensation and generates a carboxyl-carboxylate pair that conformationally stabilizes hydrogen bonding with the lysine residue on the coreceptor.

## RESULTS

### A carboxyl-carboxylate pair conformationally stabilizes the 2-to-1 polar residue motif

To interrogate the geometric and conformational stability of the (Asp-Asp-Lys) polar residue network, we simulated the DAP12-NKG2C complex, starting from the solution NMR structure (7), in three different protonation states of the polar residue triad (see Materials and Methods). In complex 1 (D^−^D^−^K^+^), all three residues were ionized leading to a 2-to-1 negative to positive charge ratio. In complex 2 (D^0^D^−^K^+^) and complex 3 (D^−^D^0^K^+^), the aspartate on one of the DAP12 monomers was protonated such that the charge ratio was 1-to-1. Finally, in complex 4 (D^0^D^0^K^0^) all polar groups were in their neutral protonation state. Three independent μs-long simulations were performed for each complex. To ensure structural stability in simulation, Cα root-mean squared deviations (RMSD) from the starting NMR structure were calculated for every complex. RMSD values plateaued within the first 50 ns of simulation and then remained relatively stable throughout the remainder of the microsecond simulation time, indicating equilibration in this time regime (**Figure S1**). Given that the simulations began with NMR structures, it is hence likely that we are evaluating relevant conformational ensembles on this time scale. After excluding the first 100 ns of each simulation, we performed principal component analysis (PCA) and cluster analysis of the polar group sidechains to understand the conformational variation as a function of protonation state (**Figure 1**, **S2, S3**). Structures for analysis were taken every 200 ps for a total of 15,000 structures per complex. Geometrically similar structures were grouped with a conservative RMSD cutoff of 0.5 Å (see Materials and Methods).

**Figure 1.**
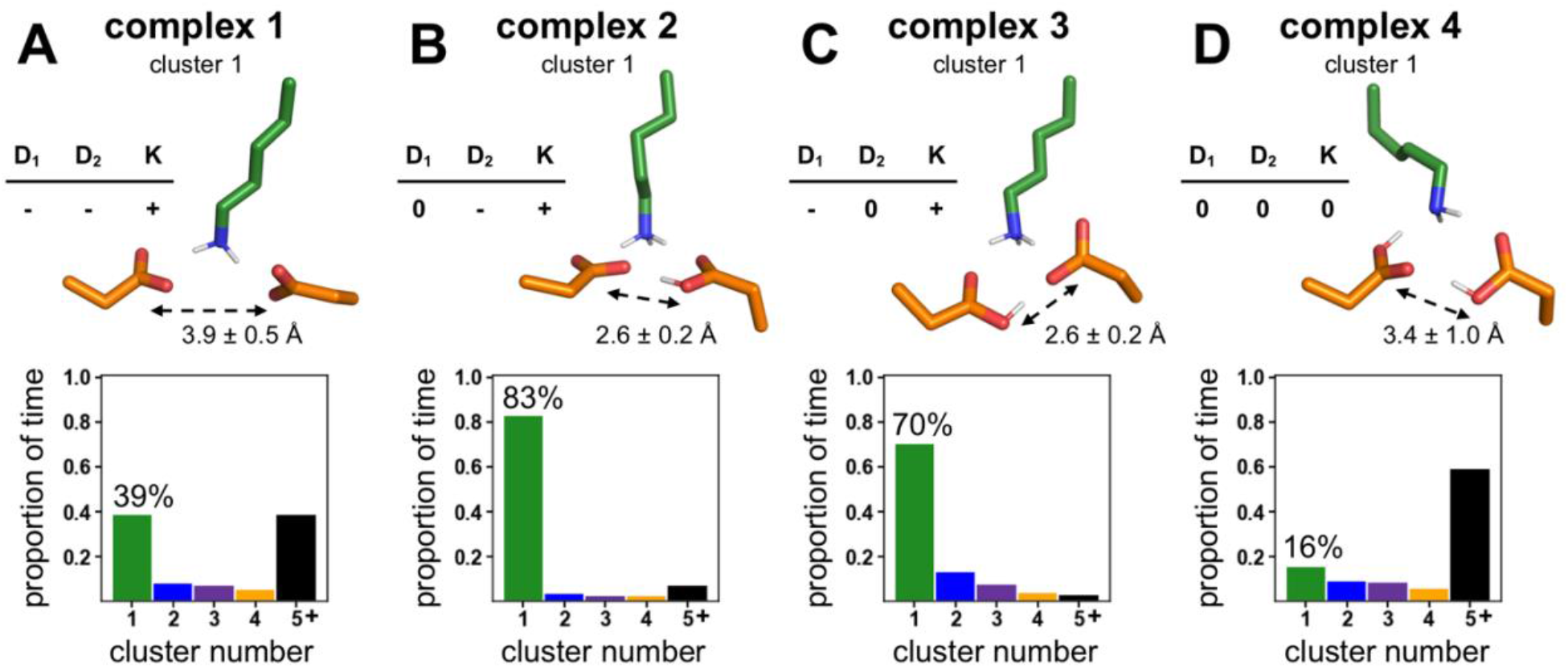
Polar group conformational arrangement as a function of protonation state. **A**. Complex 1 (D^−^D^−^K^+^) **B**. Complex 2 (D^0^D^−^K^+^) **C**. Complex 3 (D^−^D^0^K^+^) and **D**. Complex 4 (D^0^D^0^K^0^). For all complexes, structural centroid of polar residue (Asp-Asp-Lys) cluster 1 (top) and quantification of proportion of time spent in each of the clusters highlighting the percent of time spent in cluster 1 (bottom).

Analysis of complex 1 (D^−^D^−^K^+^) indicated that the geometry of the polar groups populated multiple conformational states in the course of simulation time (**Figure 1A**). The top conformer for complex 1 was populated 39% of the time and frequently fluctuated to other conformations (**Figure S4**). An unfavorable electrostatic term to the short-range nonbonded potential energy between aspartate residues suggested that the conformational dynamics were driven by electrostatic repulsion between aspartates (**Figure S5A**). Indeed, the minimum aspartate-aspartate O-O distance was long and highly fluxional (3.9 ± 0.5 Å), and in two out of three simulations a sodium ion bound the aspartates for over 40% of the time, consistent with uncompensated charge. Notably, a large influx of water molecules into the membrane (~10 waters within 3.5 Å of the polar residues) disrupted critical hydrogen bonding interactions between aspartate residues and lysine (**Figure S5 and S6**). In sum, electrostatic repulsion between two negatively charged aspartates conformationally led to a dynamic and potentially less stable polar residue triad arrangement.

In stark contrast to complex 1, the polar group arrangement was conformationally rigid when only one aspartate was protonated, as in the case of complex 2 (D^0^D^−^K^+^) and complex 3 (D^−^D^0^K^+^). The predominant polar residue conformers for these complexes were populated 83% and 70% of the time respectively (**Figure 1B-C**). The aspartic acid formed a short ionic hydrogen bond with the aspartate, average O-O distance of 2.6 ± 0.1 Å, that persisted for greater than 97% of simulation time (**Figure S7A-B**). The syn-anti arrangement of the carboxyl-carboxylate clamp between DAP12 monomers was stabilized by association with lysine on the NKG2C helix. In the absence of the NKG2C helix, the carboxyl-carboxylate hydrogen bond was either broken or adopted an energetically unfavorable anti-anti arrangement (**Figure S8**). Persistent charge interactions and hydrogen bonding between lysine and the carboxyl-carboxylate clamp effectively stapled the complex together for 83% of simulation time in the case of complex 2 (D^0^D^−^K^+^). Though three to five water molecules were always present in the proximity of this polar triad, the dynamic influx and orientation of these waters did not disrupt the conformational stability of the polar group and instead contributed to fulfilling otherwise unmet hydrogen bonds in lysine and the deprotonated aspartate (**Figure S9, S10**).

Finally, analysis of complex 4 (D^0^D^0^K^0^) demonstrated that assembly was the most conformationally dynamic when the polar residues in the triad were all in the neutral state. The top conformer for complex 4 was only populated 16% of the time (**Figure 1D**), a dynamic arrangement due, in part, to a weaker carboxyl-carboxyl hydrogen bond between protonated aspartic acid residues which persisted for 59% of simulation time (**Figure S11A**). Consistent with a weak hydrogen bond, a long and highly fluxional O-O distance of 3.4 ± 1.0 Å between aspartic acid residues was observed (**Figure S11B**). Taken together, these data demonstrate that conformational stability of the polar triad can only be achieved if both charge is compensated and there is a strong, conformationally rigid interaction between the acidic residues facilitated by the carboxyl-carboxylate pair found in complex 2 (D^0^D^−^K^+^) and complex 3 (D^−^D^0^K^+^).

### Polar triad arrangement imposed by carboxyl-carboxylate pair is supported by DFT calculations and database search

We next wanted to provide support for the polar group configurations of complex 2 (D^0^D^−^K^+^) and complex 3 (D^−^D^0^K^+^) predicted from MD simulations. Because classical MD forcefields do not account for polarization and may lead to incorrect geometries, especially for charged moieties, we first performed density functional theory (DFT) calculations to determine if this was an energetically reasonable arrangement of the polar groups. We were particularly interested in interrogating the quantum mechanical feasibility of the short carboxyl-carboxylate O-O distances that were responsible for the observed conformational stability. As a starting point, we used a model that contained the two aspartates, the lysine, a nearby threonine, and the two nearest waters with the positions extracted from the centroid of cluster 1 from complex 2 (see SI Methods). Holding the backbone atoms fixed and allowing the sidechains to relax resulted in a structure that was very similar to that observed in MD. Indeed, the RMSD of 0.24 Å is well within our clustering cutoff of 0.5 Å (see SI for details). This suggests that any restrictions imposed by parameterization and fixed geometries in the force field were not responsible for the polar group arrangement observed in these complexes. In particular, a short and strong carboxyl-carboxylate interaction was maintained in the relaxed structure (2.56 Å vs. 2.54 ± 0.07 Å, **Figure 2A, S12A**).

**Figure 2.**
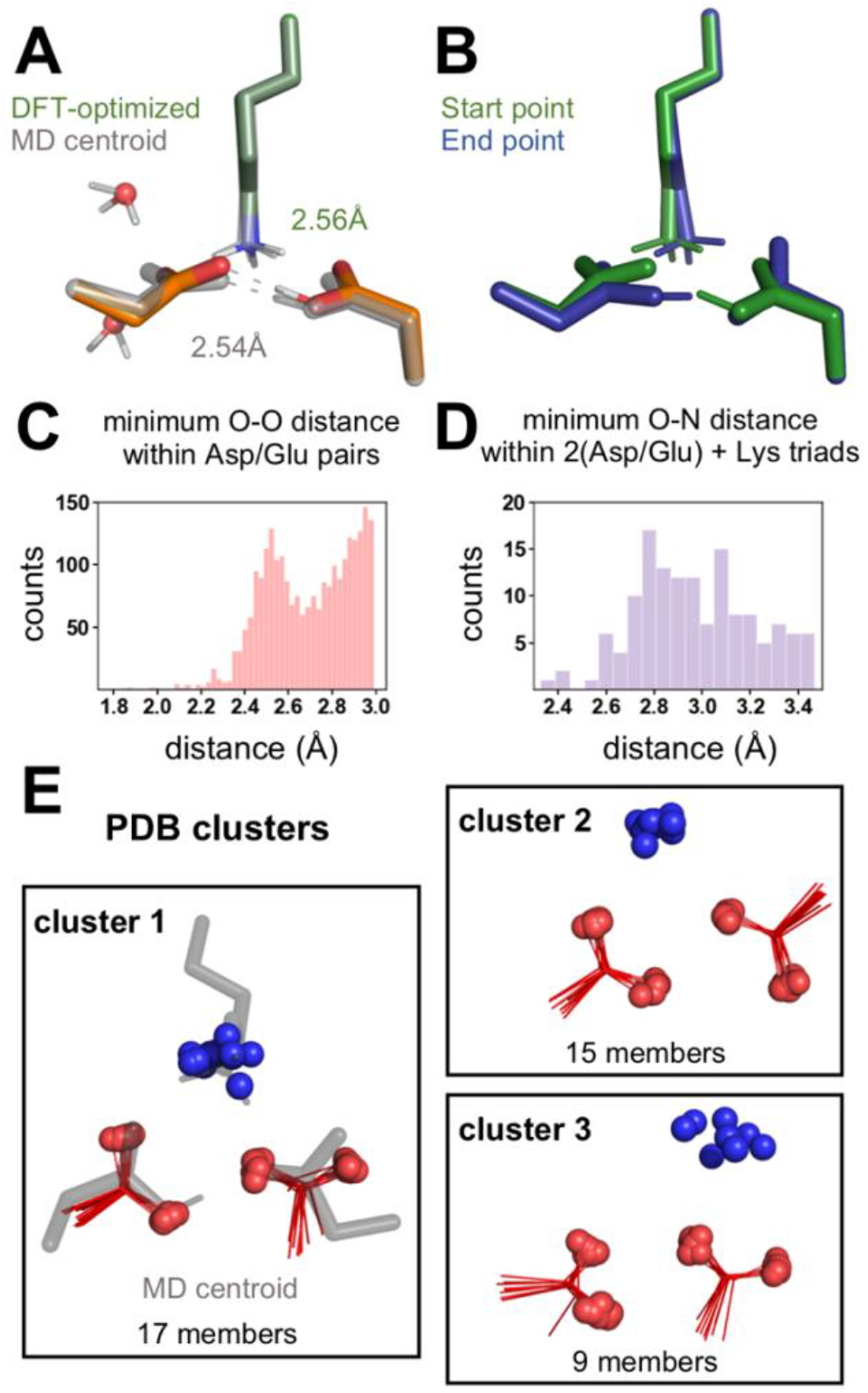
Evaluation of the (Asp-Asp-Lys) geometry imposed by monoprotonation of the acidic pair. **A**. Comparison of the DFT-optimized structure (green/orange) and centroid of cluster 1 from MD simulations of complex 2 (gray) highlighting the similarly short Asp-AspH hydrogen bond (AspH designates a protonated Asp). Overall RMSD of 0.24 Å of the clustered atoms (Cβ and Cγ of the Asp/AspH and NZ of the lysine). **B**. End points found by following the transition state mode for proton transfer between the Asp and AspH showing interconversion of complex 2 (start point, green) and complex 3 (end point, blue). Nonpolar protons, backbone atoms, and threonine are not shown in all DFT calculations. **C**. Distribution of minimum carboxylate O-O distances from the PDB. **D**. Distribution of minimum lysine-(carboxyl-carboxylate) N-O distances from the PDB. E. Geometric cluster of lysine-(carboxyl-carboxylate) triads from the PDB that are in closest agreement with complex 2 (D^0^D^−^K^+^) cluster 1 centroid from MD simulations.

Given the short nature of the carboxyl-carboxylate hydrogen bonding interaction and that carboxyl-carboxylate pairs can have geometries in which the hydrogen is shared, we wanted to evaluate the energetic penalty for proton transfer between carboxylates. In this explicitly solvated model, proton transfer is uphill by ~5 kcal/mol (**Figure S13**). However, in a model without explicit waters, we find a nearly barrierless proton transfer (Δ*G*^‡^ = +2 kcal/mol) that interconverts complex 2 (start point, **Figure 2B**) and complex 3 (end point, **Figure 2B**). These findings suggest that water reorganization may be the greatest barrier to proton transfer consistent with the nonpolar nature of the microenvironment. Nonetheless, these results strongly suggest that proton transfer is energetically feasible and, therefore, all of the conformational states of complex 2 (D^0^D^−^K^+^) and complex 3 (D^−^D^0^K^+^) are likely simultaneously available to be explored during subunit assembly (**Figure S14-S15**).

To further confirm that the geometry observed in complex 2 (D^0^D^−^K^+^) and complex 3 (D^−^D^0^K^+^) is both reasonable and frequently observed in protein structures, we analyzed lysine-(carboxyl-carboxylate) triads in the Protein Data Bank (PDB) (40) based on methods we developed in previous work (41) (see SI Methods). We first explored the occurrence frequency and geometry of proteins with glutamate or aspartate carboxylate oxygens within a distance of 3.0 Å. Strikingly, a histogram of the minimum carboxylate O-O distances is bimodal, with a local maximum near 2.5 Å (**Figure 2C**). This maximum is associated with carboxyl-carboxylate pairs based on previous surveys of the small molecule crystallographic database (42), in which the hydrogens can be assigned explicitly and is consistent with our MD and DFT results. Because of the two distinct populations in the distribution of carboxyl-carboxylate distances, we refined our dataset to only the population with the shortest O-O distances (<2.8 Å). Of those 1,583 carboxyl-carboxylate pairs, we observed 140 (9%) that form a salt bridge with the terminal sidechain atom (NZ) of a lysine (i.e., N-O distance ≤ 3.5 Å). The distribution of minimum N-O distances observed for carboxyl-carboxylate pairs is similar to that previously seen for lysine-glutamate/aspartate salt bridges (43), showing a maximum near 2.8 Å (**Figure 2D**). Importantly, this distribution is consistent with MD-derived N-O distances which vary depending on the protonation state of the aspartate, 2.8 ± 0.1 Å for the deprotonated aspartate and 3.1 ± 0.4 Å for the protonated aspartate in the case of complex 2 (D^0^D^−^K^+^). Lastly, we grouped lysine-carboxyl-carboxylate triads into geometrically similar clusters with an RMSD cutoff of 0.6 Å (see Materials and Methods). Most interestingly, the top cluster showed a very close superposition with the geometry observed for the centroid of complex 2 (D^0^D^−^K^+^) (**Figure 2E, S16**). From these data we conclude that the polar triad geometry observed in simulation is well-represented in the examples of carboxyl-carboxylate ammonium triads found in the PDB.

### Mutant data support a biological role for the carboxyl-carboxylate pair

We wanted to understand the biological importance of a conformationally static arrangement of the polar residues. We simulated mutants that have been shown experimentally to have reduced complex assembly relative to the wild type DAP12-NKG2C complex and interrogated the conformational dynamics of their polar triad. First, we simulated a conservative asparagine mutation (ND^−^K^+^) which was previously found to experimentally assemble with 27% of the efficiency of wild type (5). The top conformer of this mutant had a geometry and hydrogen bonding network that closely resembled that of wild type complex 2 (D^0^D^−^K^+^) but was observed only 26% of simulation time (**Figure 3A, S17A**). A carboxamide-carboxylate bond with an average O-N distance of 2.9 ± 0.2 Å stabilized the DAP12 monomers for interaction with the coreceptor lysine. Analogous to DFT calculations performed on the wild type, we used the centroid of cluster 1 as the input structure and allowed the sidechains to relax. This protocol again resulted in a structure in good agreement with the MD centroid (**Figure S12C**). Then, natural bond orbital (NBO) calculations were performed on this asparagine mutant and on wild type complex 2 to provide insight into the relative strength of non-covalent interactions in the polar region. Consistent with other studies, we found that the Asn-Asp hydrogen bond was approximately 20% weaker than the carboxyl-carboxylate hydrogen bond in the wild type complex (**Figure S12B**). These data are consistent with previous computational evaluation of carboxylate/carboxamide substitutions (39) and in line with the broader finding that hydrogen bonding distance and strength is strongly correlated with the p*K*_a_ match between the donor and acceptor partner (44). Consistent with a longer and weaker interaction, the Asn-Asp hydrogen bond was broken in 49% of MD simulation time (**Figure S17C-D**). These combined data suggest that a short (2.6 ± 0.1 Å) and strong carboxyl-carboxylate hydrogen bond between wild type DAP12 monomers is necessary for a conformationally stable interaction with the lysine on the NKG2C coreceptor and that even small deviations in the hydrogen bond strength are conformationally destabilizing.

**Figure 3.**
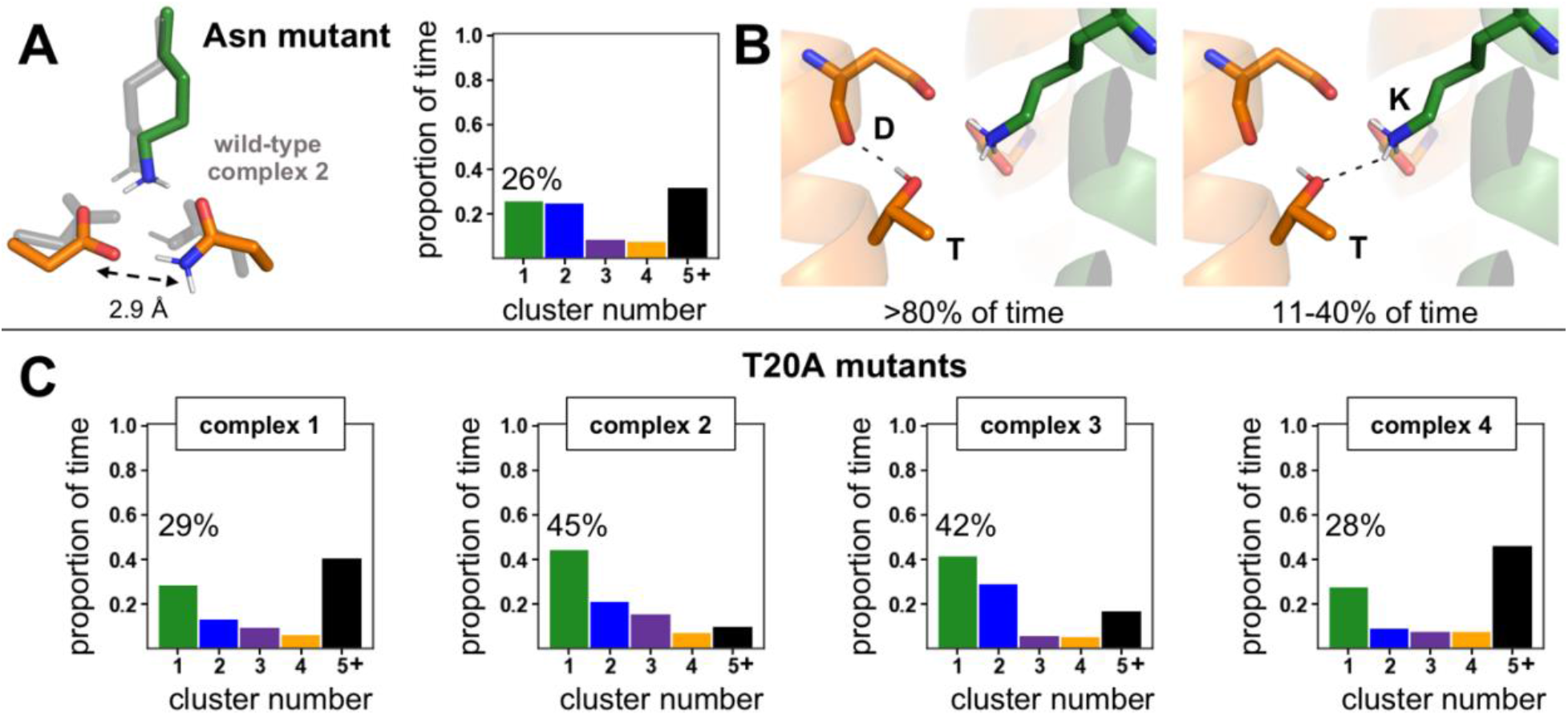
The role of mutation on polar triad conformational stability. **A**. Comparison of the top polar residue conformer of the asparagine mutant (ND^−^K^+^) to the top conformer of wild type complex 2 (D^0^D^−^K^+^) (left) and a quantification of the proportion of time this mutant arrangement is present in simulation (right) **B**. The two roles of threonine in complex assembly: threonine-aspartate sidechain to main chain hydrogen bonding present >80% of simulation time (left) and threonine-lysine hydrogen bonding present 11-40% of simulation time depending on ionization state (right) **C**. Quantification of proportion of frames spent in each of the polar residue (Asp-Asp-Lys) clusters for T20A mutants of complex 1 (D^−^D^−^K^+^), complex 2 (D^0^D^−^K^+^), complex 3 (D^−^D^0^K^+^), and complex 4 (D^0^D^0^K^0^). The percent of time spent in the top cluster is highlighted.

Previous experimental studies have also shown that an alanine mutation of a threonine residue present one helical turn (i+4 residues) away from each aspartate on DAP12 resulted in a 70% loss of assembly with NKG2C (7). To understand how this threonine to alanine (T20A) mutation on DAP12 influenced polar triad conformational stability, we compared T20A mutants in each protonation state to wild type complexes. Threonine performed two essential functions in wild type assembly: 1) it formed a sidechain to main chain hydrogen bond with the aspartate carbonyl oxygen, rigidifying that stretch of the DAP12 helix for greater than 80% of simulation time and 2) it directly engaged the lysine residue through hydrogen bonding for 11% to 40% of simulation time depending on protonation state (**Figure 3B, S18**). The T20A mutation conformationally destabilized the Asp-Asp-Lys polar triad (**Figure 3C, S19A**). Mutation in complex 2 (D^0^D^−^K^+^) and complex 3 (D^−^D^0^K^+^) resulted in a dramatic decrease in the population of the top conformer, with the T20A mutant of complex 2 (D^0^D^−^K^+^) exhibiting 45% representation compared to 83% in the wild type. This conformational destabilization was not as pronounced in complex 1 (D^−^D^−^K^+^), suggesting that the threonine residue in this protonation state would not contribute considerably to observed assembly, inconsistent with the experimental data. Consistent with this observation, the number of waters that formed hydrogen bonds with lysine increased with the T20A mutation only for complex 2 (D^0^D^−^K^+^) and complex 3 (D^−^D^0^K^+^) (**Figure S19B**). These T20A mutant data again demonstrate that local polar group conformational stability correlates to experimentally observed assembly in the DAP12-NKG2C complex. Importantly, this and the asparagine mutant data support the hypothesis that persistence of the polar groups in the specific arrangement necessary to stabilize DAP12-NKG2C complex assembly is only possible when the aspartate pair of wild type DAP12 is monoprotonated to form a carboxyl-carboxylate clamp as in complex 2 (D^0^D^−^K^+^) and complex 3 (D^−^D^0^K^+^).

### Complex assembly is globally dynamic

With substantial evidence in support of a charge-compensated polar group arrangement, we sought to determine how local polar group conformational stability influenced helix-helix conformational dynamics. We reasoned that helix rearrangement dynamics would occur over longer time scales, and in an effort to capture more conformational space, we performed two additional independent 1 μs-long simulations of complex 2 (D^0^D^−^K^+^), for a total of 5 μs. The local polar triad arrangement of the 5 μs of simulations was in agreement with the arrangement found after 3 μs (**Figure S20A**) with a cluster 1 centroid RMSD of 0.13 Å. We then clustered simulations and performed PCA at the helix level based on the Cα RMSD of the 11 core residues in each helix including the three polar residues in the triad with a 1.5 Å RMSD cutoff (see Materials and Methods). Surprisingly, we found a conformationally dynamic arrangement of helices with multiple clusters (**Figure 4A**). Visual inspection of aligned cluster centroids and RMSD analysis of helix pairs within the trimer suggested that the position of the NKG2C coreceptor helix relative to DAP12 varied considerably (**Figure 4B, S20B**). To corroborate this observation, we then used PCA to identify the greatest source of conformational variance. Principal component 1 (PC-1) accounted for 47.8% of variance and represented the translational motion of the NKG2C helix from close engagement with DAP12-1 to DAP12-2 (**Figure 4C-D**). PC-2 accounted for variance in the DAP12 homodimer crossing angle, but the biggest differentiating factor between clusters was the NKG2C coreceptor position. In sum, a local, fixed polar group arrangement did not dictate the helix arrangement. Instead, the conformational arrangement of the polar groups in this complex was stably maintained even as the NKG2C position relative to the DAP12 homodimer varied such that the helical interface remained dynamic.

**Figure 4.**
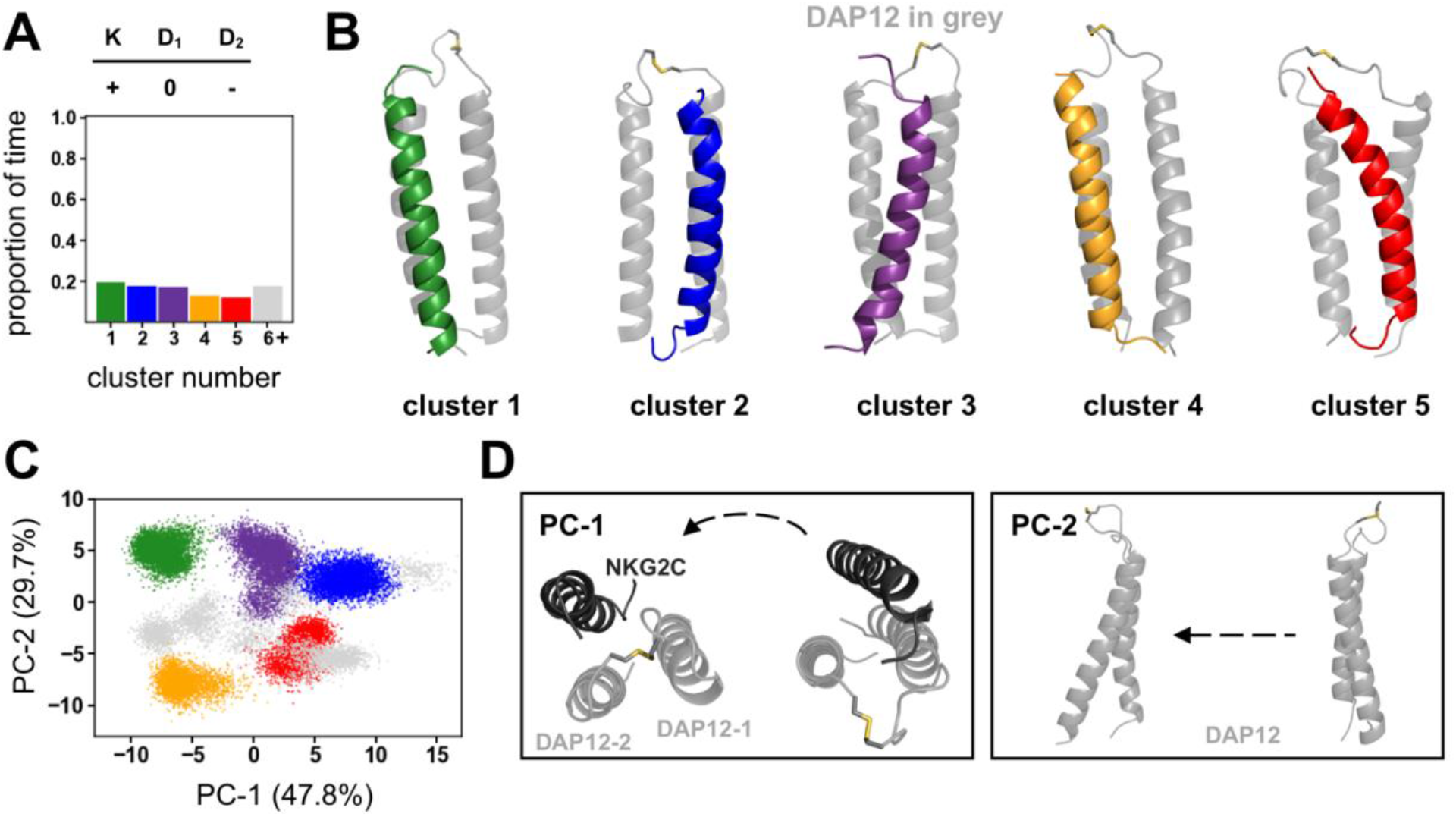
Global helix dynamics of complex 2 (D^0^D^−^K^+^) **A**. Quantification of proportion of time DAP12-NKG2C complex spends in geometrically different 33 residue Cα clusters. **B**. Structural centroids of the top five clusters. **C**. Principal component analysis of global helix residues. Projections of structural coordinates along PC-1 and PC-2 (points are colored according to the helical clustering analysis) **D**. Visual representation of the global helical motions along PC-1 and PC-2.

## DISCUSSION

Multi-component receptor assembly that proceeds through a 2-to-1 acidic to basic polar residue triad is conformationally static at the local polar residue level only if the acidic residue pair is monoprotonated. Atomistic MD simulations of the DAP12-NKG2C complex have demonstrated that when both aspartates are deprotonated multiple conformational states of the polar residues are populated due to electrostatic repulsion of the aspartates. Importantly, increased conformational dynamics of the polar residues correlate with reduced experimental complex assembly in asparagine and threonine mutants. There is no appreciable difference in polar residue dynamics between the wild type doubly deprotonated state and these mutants, allowing us to establish this protonation state as unproductive for assembly. Protonation of one of the aspartates eliminates electrostatic repulsion and generates a stabilizing carboxyl-carboxylate clamp with a nearly invariant 2.6 Å hydrogen bonding interaction that more than doubles the conformational stability of the polar triad. The carboxyl-carboxylate clamp engages with the coreceptor lysine in a conformationally persistent arrangement that was found to be energetically favorable through DFT and well-precedented in soluble protein structures. The short carboxyl-carboxylate hydrogen bond between DAP12 monomers that staples this triad together is of particular interest because it carries the potential for added conformational stability. Given that the proton in the carboxyl-carboxylate pair has a low barrier for transfer between the two carboxylate groups, complex 2 (D^0^D^−^K^+^) and complex 3 (D^−^D^0^K^+^) can each be considered a limiting case, imposed by the classical force field, of what is likely a shared interaction. Therefore, both of the tautomeric states of complex 2 (D^0^D^−^K^+^) and complex 3 (D^−^D^0^K^+^) are likely available to be explored during subunit assembly. Conceivably, all residues in the triad could be in the neutral state if lysine (intrinsic p*K*_a_ = 10.5) were deprotonated, but some coreceptors rely on arginine (intrinsic p*K*_a_ = 12) for association which is more likely to remain ionized. Therefore, we reasoned that lysine should also be charged, and the data from complex 4 (D^0^D^0^K^0^) corroborate our hypothesis that an ionized basic residue is necessary for receptor assembly.

A monoprotonated acidic pair was suggested previously to be unlikely because of a 3-fold decrease in complex assembly efficiency in the conservative mutation of one aspartate to asparagine (5). However, this change in stability corresponds to less than a single kcal/mol, and a carboxamide-carboxylate pair is not a perfect mimic of the hydrogen bonding and electrostatic network of a carboxyl-carboxylate pair in the partially protonated wild type system. Moreover, aspartate to asparagine mutations have been previously shown to decrease modestly (2 to 3 °C) the thermostability of carboxyl-carboxylate pairs (45), a difference that could account for the similarly modest effect of an Asp-to-Asn mutation on assembly of the DAP12-NKG2C complex. In line with literature precedent, we find that although asparagine shares many characteristics with aspartic acid, it is not a perfect mimic. The 2.6 Å carboxyl-carboxylate hydrogen bond conformationally stabilizes the DAP12 dimer interface, while a longer 2.9 Å carboxamide-carboxylate bond makes this point of attachment weaker and more fluxional in the asparagine mutant. This slight destabilization of the DAP12 homodimer interface, turns out to be conformationally destabilizing for assembly with the lysine residue on the coreceptor. These subtle changes observed due to conservative mutations highlight the means by which the DAP12 signaling subunit ensures specific assembly with a coreceptor as long as the coreceptor contains a properly positioned lysine residue.

One notable feature of the 2-to-1 polar residue motif is its residence in the nonpolar membrane. Because MD force fields represent the structures of both water-soluble and membrane-soluble proteins well, the geometry of the local atomic interactions in the polar triad observed in simulation would be expected to hold across a wide range of environments. Indeed, the DFT analysis and the PDB search for this motif corroborate the conclusion that this geometry is locally optimized and largely invariant with environment. Conversely, the energetics of polar triad association, though locally favorable, will vary depending on the folding environment. Energetics of folding are based on the unfolded or reference state of a system. In the case of multicomponent receptor assembly, the “unfolded state” is composed of non-interacting, membrane-inserted helices with largely desolvated polar groups due to the nonpolar environment. Previous studies with model peptides have shown that aspartic acid in the membrane interior has an elevated p*K*_a_, which allows protonation near neutral pH (29–35). Folding would bring the interacting groups together without incurring the energetically costly dehydration of the polar sidechains or protonation of one of the two carboxyl groups involved in the carboxyl-carboxylate clamp. By contrast, the formation of a (carboxyl-carboxylate)-lysine triad in water-soluble proteins would require a significant dehydration penalty as the chain folds from the hydrated random coil to its native structure (46). We suspect that this energetic advantage in the membrane is one of the reasons why this 2-to-1 motif is so prevalent across multi-component receptor families. We further expect that (carboxyl-carboxylate)-lysine/arginine triads will be found across a variety of other membrane proteins as more structures become available. The precise geometries will likely vary, based on the surrounding sequence, but the database search performed here should be generally helpful for identifying favorable geometries.

Interestingly, the local rigidity in the polar group arrangement observed to be necessary for specific assembly did not carry forward to helix-helix arrangement and packing which was conformationally dynamic. We propose that it is this locally static and globally dynamic arrangement that enables DAP12 to assemble specifically with so many sequence diverse ligand-binding coreceptors. Because the global complex arrangement is inherently dynamic without disrupting the polar residue arrangement, it is better able to accommodate the sequence diversity. Additionally, the lack of a single structural conformation explains, in part, why detailed biophysical structural characterization has been so challenging in these systems. Though a great degree of sequence tolerance is built into the association of DAP12 with a coreceptor and specificity of assembly is centered on the polar residue triad (5–7), the remaining transmembrane residues do play a role in specificity which is why DAP12 cannot assemble with a coreceptor like NKG2D (47). The T20A mutant data clearly demonstrate the importance of distal residues on proper complex assembly, but additional studies are necessary to illuminate the nature of specificity-generating interactions in the rest of the transmembrane sequence.

The work presented here has clarified details of protein subunit assembly structure and dynamics which define effectiveness in multi-component receptors. These details help inform the mechanism of signal transduction across the membrane in this class of receptors which is still unknown. The 2-to-1 acidic to basic polar residue association motif within the membrane is found in receptors where signaling is thought to be mediated by extracellular clustering of the ligand-binding subunits (1, 14, 48–50). This clustering hypothesis proposes that increased local concentrations of ITAM motifs dictate the efficiency of receptor phosphorylation and subsequent downstream signaling (51–53). This suggests that the goal of the transmembrane association motif is to staple the ligand-binding and signaling subunits in a stable conformation such that when extracellular receptor clustering in response to a ligand occurs, the signaling subunits also cluster. For receptors that are hypothesized to signal through extracellular clustering, stable assembly with the signaling coreceptor, in this case DAP12, is necessary for clustering of signaling motifs. The carboxyl-carboxylate clamp on DAP12 is strong enough to staple to lysine on a coreceptor and allows the two subunits to move in the membrane as one receptor. Additionally, the dynamic helix packing found for the DAP12-NKG2C receptor complex further supports this signaling-by-clustering hypothesis. The global dynamics observed appear to contradict a ligand-induced conformational change mechanism. Instead, dynamic helix rearrangement enables the ligand-binding subunit to freely adapt its position and orientation while still maintaining a stable point of attachment to the signaling subunit through the polar motif. This global conformational flexibility would be important for cluster-mediated signaling in order to allow the conformational changes necessary for clustering with other receptors in response to a ligand.

## CONCLUSION

Overall, the findings reported here provide an atomic level picture that help explain the simultaneous specificity and promiscuity of DAP12 for lysine containing receptors with very little sequence homology. DAP12 assembly with a coreceptor is mediated by a monoprotonated aspartate pair, not a doubly deprotonated aspartate pair as was previously proposed. This carboxyl-carboxylate pair forms a short and strong hydrogen bond that mediates a conformationally static interaction with the coreceptor lysine. Specificity of interaction with the lysine results from both the conformational restriction and directionality imparted on the aspartate-aspartic acid pair by such a tight hydrogen bond as well as hydrogen bonding of lysine to a distal threonine. This local fixed arrangement does not extend to the helix level, which is found to be highly flexible. Thus, the DAP12 signaling module is able to promiscuously interact with many sequence diverse coreceptors as long as they retain the lysine residue that imparts stability. These findings elucidate details of receptor assembly that proceeds by means of a 2 -to-1 acidic to basic polar residue motif and are widely applicable across receptor families because this motif is so pervasive. Further, this work provides evidence in support of cluster-mediated signaling in this class of receptors, which needs to be stable enough to ensure subunit assembly while also dynamically adaptable to respond to a ligand.

## MATERIALS AND METHODS

All MD simulations used the DAP12-NKG2C NMR structure (PDB id: 2L35) (7). The structure was embedded in a POPC lipid membrane and solvated in 150 mM NaCl using CHARMM-GUI (54). All production runs were performed in the isothermal-isobaric (NPT) ensemble using the AMBER ff14SB protein force field and the lipid17 lipid force field with the Amber GPU engine (55–57). Changes to the ionization state of the polar residues and mutations were made in the AMBER LEaP program (58). Three independent 1.1 μs simulations were performed per ionization or mutation state starting from the same initial position but different starting velocities. For analysis, trajectories were combined for a total of 3 μs per ionization state (after discarding the first 100ns of each simulation). Two additional simulations were performed for complex 2 for analysis of global helix dynamics.

For DFT calculations, the input positions were taken from the centroid for the top cluster from the complex of interest. The inputs were completed by appropriately terminating the N- and C-termini of each residue. Then, optimization, frequency, and transition state calculations were done with the ORCA software package (version 4.2) (59). Natural bond orbital (NBO) calculations were performed with the Gaussian09 package (60, 61).

For the PDB search, we used a non-redundant dataset of 13,026 protein crystal structures curated by PISCES (62) on September 14th, 2020 with ≤ 30% sequence identity, R factor of ≤ 0.3, and resolution of ≤ 2.5 Å. All structures were re-refined by PDB-REDO (63). We searched for lysine-(carboxyl-carboxylate) triads, and identical triads within oligomers belonging to the same structure were eliminated.

All clustering analysis was performed using a hierarchical agglomerative average-linkage approach (64) with a 0.5 and 1.5 Å cutoff for the local polar group and global helix analysis of MD simulations respectively. A 0.6 Å cutoff was used for clustering of PDB structures.

Further details on all MD, DFT, database search and analysis methods can be found in the SI Methods.

## Supporting information

SI PDB Cluster Codes

SI DFT Coordinates

## ACKNOWLEDGEMENTS

We thank Kevin J. Metcalf, Paola Bisignano and Andrew Natale for useful discussions. This work was supported by the National Institutes of Health.

## REFERENCES

1. M. E. Call, K. W. Wucherpfennig, Common themes in the assembly and architecture of activating immune receptors. Nat. Rev. Immunol. 7, 841–850 (2007).

2. L. L. Lanier, DAP10- and DAP12-associated receptors in innate immunity. Immunol. Rev. 227, 150–160 (2009).

3. L. L. Lanier, Turning on natural killer cells. J. Exp. Med. 191, 1259–1262 (2000).

4. M. B. Humphrey, L. L. Lanier, M. C. Nakamura, Role of ITAM-containing adapter proteins and their receptors in the immune system and bone. Immunol. Rev. 208, 50–65 (2005).

5. J. Feng, D. Garrity, M. E. Call, H. Moffett, K. W. Wucherpfennig, Convergence on a distinctive assembly mechanism by unrelated families of activating immune receptors. Immunity 22, 427–438 (2005).

6. J. Feng, M. E. Call, K. W. Wucherpfennig, The assembly of diverse immune receptors is focused on a polar membrane-embedded interaction site. PLoS Biol 4, e142 (2006).

7. M. E. Call, K. W. Wucherpfennig, J. J. Chou, The structural basis for intramembrane assembly of an activating immunoreceptor complex. Nat. Immunol. 11, 1023–1029 (2010).

8. A. Blazquez-Moreno et al., Transmembrane features governing Fc receptor CD16A assembly with CD16A signaling adaptor molecules. Proc. Natl. Acad. Sci. U.S.A. 114, E5645–E5654 (2017).

9. N. Manolios, J. S. Bonifacino, R. D. Klausner, Transmembrane helical interactions and the assembly of the T cell receptor complex. Science 249, 274–277 (1990).

10. M. E. Call, J. Pyrdol, M. Wiedmann, K. W. Wucherpfennig, The organizing principle in the formation of the T cell receptor-CD3 complex. Cell 111, 967–979 (2002).

11. M. E. Call et al., The structure of the zetazeta transmembrane dimer reveals features essential for its assembly with the T cell receptor. Cell 127, 355–368 (2006).

12. L. L. Lanier, B. C. Corliss, J. Wu, C. Leong, J. H. Phillips, Immunoreceptor DAP12 bearing a tyrosine-based activation motif is involved in activating NK cells. Nature 391, 703–707 (1998).

13. L. L. Lanier, B. Corliss, J. Wu, J. H. Phillips, Association of DAP12 with activating CD94/NKG2C NK cell receptors. Immunity 8, 693–701 (1998).

14. I. R. Turnbull, M. Colonna, Activating and inhibitory functions of DAP12. Nat. Rev. Immunol. 7, 155–161 (2007).

15. J. Paloneva et al., Loss-of-function mutations in TYROBP (DAP12) result in a presenile dementia with bone cysts. Nat. Genet. 25, 357–361 (2000).

16. J. Paloneva et al., Mutations in two genes encoding different subunits of a receptor signaling complex result in an identical disease phenotype. Am. J. Hum. Genet. 71, 656–662 (2002).

17. J. Paloneva et al., DAP12/TREM2 deficiency results in impaired osteoclast differentiation and osteoporotic features. J. Exp. Med. 198, 669–675 (2003).

18. S. T. Chen et al., CLEC5A is critical for dengue-virus-induced lethal disease. Nature 453, 672–676 (2008).

19. R. Thomas et al., NKG2C deletion is a risk factor of HIV infection. AIDS Res. Hum. Retroviruses 28, 844–851 (2012).

20. J. D. Ulrich, T. K. Ulland, M. Colonna, D. M. Holtzman, Elucidating the role of TREM2 in Alzheimer’s disease. Neuron 94, 237–248 (2017).

21. F. L. Yeh, D. V. Hansen, M. Sheng, TREM2, microglia, and neurodegenerative diseases. Trends. Mol. Med. 23, 512–533 (2017).

22. L. L. Lanier, A. B. Bakker, The ITAM-bearing transmembrane adaptor DAP12 in lymphoid and myeloid cell function. Immunol. Today 21, 611–614 (2000).

23. X. Cheng, W. Im, NMR observable-based structure refinement of DAP12-NKG2C activating immunoreceptor complex in explicit membranes. Biophys. J. 102, L27–29 (2012).

24. S. Sharma, A. H. Juffer, An atomistic model for assembly of transmembrane domain of T cell receptor complex. J. Am. Chem. Soc. 135, 2188–2197 (2013).

25. H. Sun, H. Chu, T. Fu, H. Shen, G. Li, Theoretical elucidation of the origin for assembly of the DAP12 dimer with only one NKG2C in the lipid membrane. J. Phys. Chem. B. 117, 4789–4797 (2013).

26. P. Wei, B. K. Zheng, P. R. Guo, T. Kawakami, S. Z. Luo, The association of polar residues in the DAP12 homodimer: TOXCAT and molecular dynamics simulation studies. Biophys. J. 104, 1435–1444 (2013).

27. P. Wei et al., Molecular dynamic simulation of the self-assembly of DAP12-NKG2C activating immunoreceptor complex. PLoS One 9, e105560 (2014).

28. N. Dube, J. K. Marzinek, R. C. Glen, P. J. Bond, The structural basis for membrane assembly of immunoreceptor signalling complexes. J. Mol. Model. 25, 277 (2019).

29. H. Gratkowski, J. D. Lear, W. F. DeGrado, Polar side chains drive the association of model transmembrane peptides. Proc. Natl. Acad. Sci. U.S.A. 98, 880–885 (2001).

30. F. X. Zhou, H. J. Merianos, A. T. Brunger, D. M. Engelman, Polar residues drive association of polyleucine transmembrane helices. Proc. Natl. Acad. Sci. U.S.A. 98, 2250–2255 (2001).

31. G. A. Caputo, E. London, Cumulative effects of amino acid substitutions and hydrophobic mismatch upon the transmembrane stability and conformation of hydrophobic α-helices. Biochemistry 42, 3275–3285 (2003).

32. G. A. Caputo, E. London, Position and ionization state of Asp in the core of membrane-inserted α helices controlboth the equilibrium between transmembrane and nontransmembrane helix topography and transmembrane helix positioning. Biochemistry 43, 8794–8806 (2004).

33. A. Senes, D. E. Engel, W. F. DeGrado, Folding of helical membrane proteins: the role of polar, GxxxG-like and proline motifs. Curr. Opin. Struct. Biol. 14, 465–479 (2004).

34. D. T. Moore, B. W. Berger, W. F. DeGrado, Protein-protein interactions in the membrane: sequence, structural, and biological motifs. Structure 16, 991–1001 (2008).

35. K. Shahidullah, S. S. Krishnakumar, E. London, The effect of hydrophilic substitutions and anionic lipids upon the transverse positioning of the transmembrane helix of the ErbB2 (neu) protein incorporated into model membrane vesicles. J. Mol. Biol. 396, 209–220 (2010).

36. A. Langkilde et al., Short strong hydrogen bonds in proteins: a case study of rhamnogalacturonan acetylesterase. Acta. Crystallogr. D. Biol. Crystallogr. D64, 851–863 (2008).

37. L. Sawyer, M. N. James, Carboxyl-carboxylate interactions in proteins. Nature 295, 79–80 (1982).

38. M. M. Flocco, S. L. Mowbray, Strange bedfellows: interactions between acidic side-chains in proteins. J. Mol. Biol. 254, 96–105 (1995).

39. G. Wohlfahrt, Analysis of pH-dependent elements in proteins: geometry and properties of pairs of hydrogen-bonded carboxylic acid side-chains. Proteins 58, 396–406 (2005).

40. H. M. Berman et al., The protein data bank. Nucleic Acids Res. 28, 235–242 (2000).

41. S. K. Tan et al., Modulating integrin alphaIIbbeta3 activity through mutagenesis of allosterically regulated intersubunit contacts. Biochemistry 58, 3251–3259 (2019).

42. L. D’Ascenzo, P. Auffinger, A comprehensive classification and nomenclature of carboxyl-carboxyl(ate) supramolecular motifs and related catemers: implications for biomolecular systems. Acta. Crystallogr. B Struct. Sci. Cryst. Eng. Mater. 71, 164–175 (2015).

43. J. E. Donald, D. W. Kulp, W. F. DeGrado, Salt bridges: geometrically specific, designable interactions. Proteins 79, 898–915 (2011).

44. L. C. Remer, J. H. Jensen, Toward a general theory of hydrogen bonding: the short, strong hydrogen bond [HOH···OH]. J. Phys. Chem. A 104, 9266–9275 (2000).

45. J. Lin, E. Pozharski, M. A. Wilson, Short carboxylic acid-carboxylate hydrogen bonds can have fully localized protons. Biochemistry 56, 391–402 (2017).

46. W. F. DeGrado, H. Gratkowski, J. D. Lear, How do helix-helix interactions help determine the folds of membrane proteins? Perspectives from the study of homo-oligomeric helical bundles. Protein Sci. 12, 647–665 (2003).

47. J. Wu, H. Cherwinski, T. Spies, J. H. Phillips, L. L. Lanier, DAP10 and DAP12 form distinct, but functionally cooperative, receptor complexes in natural killer cells. J. Exp. Med. 192, 1059–1068 (2000).

48. R. N. Germain, T-cell signaling: the importance of receptor clustering. Curr. Biol. 7, R640–644 (1997).

49. A. M. Duchemin, L. K. Ernst, C. L. Anderson, Clustering of the high affinity Fc receptor for immunoglobulin G (Fc gamma RI) results in phosphorylation of its associated gamma-chain. J. Biol. Chem. 269, 12111–12117 (1994).

50. Y. Ma et al., Clustering of the zeta-chain can initiate T cell receptor signaling. Int. J. Mol. Sci. 21 (2020).

51. D. M. Underhill, H. S. Goodridge, The many faces of ITAMs. Trends Immunol. 28, 66–73 (2007).

52. P. E. Love, E. W. Shores, ITAM multiplicity and thymocyte selection: how low can you go? Immunity 12, 591–597 (2000).

53. J. R. James, Tuning ITAM multiplicity on T cell receptors can control potency and selectivity to ligand density. Sci. Signal 11 (2018).

54. S. Jo, T. Kim, W. Im, Automated builder and database of protein/membrane complexes for molecular dynamics simulations. PloS one 2, e880–e880 (2007).

55. J. A. Maier et al., ff14SB: Improving the accuracy of protein side chain and backbone parameters from ff99SB. J. Chem. Theory Comput. 11, 3696–3713 (2015).

56. C. J. Dickson et al., Lipid14: The Amber lipid force field. J. Chem. Theory Comput. 10, 865–879 (2014).

57. R. Salomon-Ferrer, A. W. Gotz, D. Poole, S. Le Grand, R. C. Walker, Routine microsecond molecular dynamics simulations with AMBER on GPUs. 2. explicit solvent particle mesh Ewald. J. Chem. Theory Comput. 9, 3878–3888 (2013).

58. D.A. Case et al. AMBER 2016. University of California, San Francisco (2016).

59. F. Neese, Software update: the ORCA program system, version 4.0. WIREs Computational Molecular Science 8, e1327 (2018).

60. E. D. Glendening, A. E. Reed, J.E. Carpenter, F. Weinhold NBO Version 3.1., Gaussian Inc., Pittsburgh (2003).

61. M. J. Frisch et al. Gaussian 09, Revision E.01 (2015).

62. G. Wang, R. L. Dunbrack, Jr., PISCES: a protein sequence culling server. Bioinformatics 19, 1589–1591 (2003).

63. R. P. Joosten, F. Long, G. N. Murshudov, A. Perrakis, The PDB_REDO server for macromolecular structure model optimization. IUCrJ 1, 213–220 (2014).

64. J. Shao, S. W. Tanner, N. Thompson, T. E. Cheatham, Clustering molecular dynamics trajectories: 1. characterizing the performance of different clustering algorithms. J. Chem. Theory Comput. 3, 2312–2334 (2007).

