## Supplementary Information for "Elucidation of the molecular interactions that enable stable assembly and structural diversity in multicomponent immune receptors"

Table of Contents:

I. Molecular Dynamics Simulations Methods (Table S1)

II. Density Functional Theory Methods

III. PDB Database Search Methods

IV. Polar Residue Protonation (Figures S1-S11)

V. Discussion of DFT Calculations (Figures S12-S15, Table S2)

VI. PDB Clusters (Figure S16, Table S3)

VII. Effect of Mutations on Polar Residues (Figures S17-S19)

VIII. Helix Dynamics (Figure S20)

IX. References

### I. MOLECULAR DYNAMICS SIMULATIONS METHODS

*System Set Up*

The coordinates for the DAP12-NKG2C complex were obtained from the average NMR structure (PDB id: 2L35) (1).In the NMR structure, one of the DAP12 monomers is covalently linked to the NKG2C coreceptor by a single glycine residue. While this linker is useful for ensuring covalent stability of the DAP12-NKG2C helix trimer in experiment, we were concerned that the linker could bias complex stability for the purposes of testing our protonation state hypothesis. To ensure that we were not biasing complex stability, we removed the glycine residue and allowed the DAP12 and NKG2C monomers to move independently. The N-terminus of NKG2C was then acetylated, and the C-termini of all three helices were amidated. Using the CHARMM-GUI webserver (2), the DAP12-NKG2C complex was embedded in a palmitoyl-oleoyl phosphatidylcholine (POPC) bilayer and hydrated (minimum water height of 20Å). $Na^+$ ions were added to neutralize and the NaCl concentration was brought to 150mM. The system was converted to the AMBER format with a combination of in-house scripts and the AMBER LEaP program in AmberTools 16 (3). Changes to the ionization state and mutations were also made in LEaP. Details of the simulations performed are listed in **Table S1**.

**Table S1**: Description of atomistic simulations performed of DAP12-NKG2C complex

| System | Protonation/Mutation State | Total Atoms | Simulation time (μs) x replicas |
|---|---|---|---|
| **complex 1** | Asp-Asp-Lys | 31,477 | 1 x 3 |
| **complex 2** | AspH-Asp-Lys | 31,476 | 1 x 5 |
| **complex 3** | Asp-AspH-Lys | 31,476 | 1 x 3 |
| **complex 4** | AspH-AspH-Lys | 31,476 | 1 x 3 |
| **Nmutant** | Asn-Asp-Lys | 31,477 | 1 x 3 |
| **Tmutant 1** | Asp-Asp-Lys, Thr20Ala | 31,469 | 1 x 3 |
| **Tmutant 2** | AspH-Asp-Lys, Thr20Ala | 31,468 | 1 x 3 |
| **Tmutant 3** | Asp-AspH-Lys, Thr20Ala | 31,468 | 1 x 3 |
| **Tmutant 4** | AspH-AspH-Lys, Thr20Ala | 31,468 | 1 x 3 |

*Atomistic Simulation Procedure*

Atomistic molecular dynamics (MD) simulations were performed with the AMBER ff14SB protein force field and the lipid17 lipid force field parameter sets (4, 5). Water molecules were described using the TIP3P model. Simulations were carried out on GPUs with the PMEMD MD engine (6). To remove high energy contacts, minimization consisted of 5000 steps, 2,500 steps of steepest descent algorithm with a switch to 2,500 steps of the conjugated gradient algorithm. For NVT equilibration, the systems were gradually heated from 0 to 310 K over 300 ps. Temperature was maintained with the Langevin thermostat with a friction coefficient of $\gamma = 1$ $ps^{-1}$. The proteins and lipids were restrained with a harmonic potential and force constant of 5 kcal/(mol·Å$^2$). Once a temperature of 310 K was reached, the system was switched to NPT and pressure was maintained at 1 atm using a semi-isotropic pressure tensor and the Monte Carlo barostat. The 5 kcal/(mol·Å$^2$) restraint on the proteins and lipids was gradually removed over 1 ns. During the production period, the system was simulated for a total of 1.1 μs. Simulations were carried out under periodic boundary conditions. The SHAKE algorithm was used to constrain bonds involving hydrogen atoms enabling a 2 fs timestep (7). The particle-mesh Ewald method was used to compute long-range electrostatics, and non-bonded interactions were cutoff at 12 Å with force-based switching at 10 Å. Three independent simulations were performed per ionization or mutation state starting from the same initial position but different starting velocities. Two additional simulations were performed for complex 2 for analysis of global helix dynamics. Atomic coordinates were saved every 200 ps for subsequent trajectory analysis.

*Clustering Analysis*

Clustering analysis of MD simulations is used to group similar conformations of a biomolecule together in order to understand how that molecule samples conformational space during simulation time (8). Because sampling of conformational space should occur in accordance with a Boltzmann distribution such that higher energy states are sampled less than lower energy states, clustering can be used to discern stability of a particular conformation.

All structural clustering analysis was performed using a hierarchical agglomerative (i.e. bottom-up) average-linkage approach. In this approach, each structure (simulation frame) begins as its own cluster. At each step, the two closest clusters are merged based on a distance metric, coordinate root-mean-square deviation (RMSD) in this case. With average-linkage, the distance from one cluster to another is defined by determining the RMSD between every structure in cluster one and every structure in cluster two and averaging the distances. Structures that are more similar have a smaller distance between clusters. Clustering occurs until the distance between all clusters exceeds a cutoff value.

For local polar group conformations, clustering was performed on 15,000 structures per system. The RMSD of six atoms {Asp1 (Cβ, Cγ), Asp2 (Cβ, Cγ), and Lys (Cε, NZ)} was used with a 0.5 Å cutoff. The carboxyl/carboxylate oxygens in the Asp residues were excluded to ensure that flipped carboxylates/carboxylic acids in the same geometry were counted in the same cluster. This method also enabled clustering of the Asn-Asp-Lys mutant using the same strategy. For global helical conformations, clustering was performed on 25,000 structures. The Cα RMSD of 33 residues (Asp-Asp-Lys and the 10 surrounding each of these residues) was used with a 1.5 Å cutoff.

*Principal Component Analysis*

Principal component analysis (PCA) of MD simulations can be used to identify major modes of motion and explain variance in the coordinates of a molecule as a trajectory proceeds (9). We used PCA to complement the clustering analysis, because it can give insight into what concerted motions in the protein structure lead to the conformations identified from clustering.

In practice, a coordinate covariance matrix was calculated from the positional coordinates of specified atoms in the trajectory. To remove global rotations and translations of the protein structure, each frame was RMS-fit to the overall average structural coordinates. The coordinates from the trajectories of each simulation were then separately projected along each eigenvector to obtain a time series of principal component projection values for each principal component (PC). The motions along PC-1 and PC-2 provide a visual representation of what atoms move in concert to yield the observed conformations in different clusters, while a plot of PC-2 vs. PC-1 provides a quantitative 2D representation of the majority of the variance (> 75%) in the coordinate data.

For local polar group PCA, the covariance matrix was calculated with the coordinates of the same six atoms used for clustering {Asp1 (Cβ, Cγ), Asp2 (Cβ, Cγ), and Lys (Cε, NZ)} and a combined trajectory from the simulations of all four protonation states (complex 1, complex 2, complex 3, complex 4). For global helix PCA, the covariance matrix was calculated with the coordinates of the same 33 atoms used for clustering (Cα of 33 core residues including the polar triad) and using a combined trajectory from all 5 simulations of complex 2 ($D^{0}D^{-}K^{+}$).

*Hydrogen Bonding Analysis*

A hydrogen bond was considered if the distance between the acceptor and donor heavy atoms (Ex: O-O distance in the case of an Asp-Asp hydrogen bond) was less than 3.0 Å and the angle between the acceptor heavy atom, the donor proton, and the donor heavy atom was greater than 135°. For time series data, hydrogen bonds were given a binary classification, either a hydrogen bond was present, or it was not.

All analysis of MD simulations was performed using the CPPTRAJ analysis package on AmberTools 16 (10).

### II. DENSITY FUNCTIONAL THEORY METHODS

For all density functional theory (DFT) calculations the starting points for the calculation were prepared as follows. The input positions of the four residues ($D_{16}$, $T_{20}$, $D_{49}$, and $K_{85}$) and the two closest waters were taken from the MD simulations using the centroid of cluster 1 for the complex of interest. The C-terminal end of each residue was extended with the NH from the next residue (i+1) to which a proton was added to terminate with an $NH_2$ group. The N-terminal end of the residue was extended to include the C=O from the previous residue (i−1) to which a proton was added to terminate with a CH(O) group.

Optimization and frequency calculations were done with the ORCA software package (version 4.2) (11). Calculations were performed with the BP86 (12) basis set and the def2-tzvp (13, 14) functional on C and H and the def2-tzvpp (13, 14) functional on N and O. The DFT-D3 dispersion correction (15) was applied and optimizations were done with the conductor-like polarizable continuum model (CPCM) (16) with an ε of 2.4 to match with estimates of dielectric constants in the interior of a lipid bilayer (17). In optimization calculations, the heavy atoms of the backbone and of the waters (C, N, O) were constrained. All other atoms were allowed to relax. The resulting geometries were evaluated with frequency calculations, which typically identified several small imaginary stretching frequencies ($> -200\ cm^{-1}$). Given the small nature of these imaginary modes and their diffuse nature when visualized, we believe that they are a result of constraining the backbone atoms, which is necessary to retain the alpha-helical structure enforced by the protein secondary structure. As we are not evaluating the absolute energies of these structures, these small imaginary frequencies are not of concern.

### III. PDB DATABASE SEARCH METHODS

For the database search of the Protein Data Bank (PDB) (18), we used a non-redundant dataset of 13,026 protein crystal structures curated by PISCES (19) on September 14th, 2020 with ≤ 30% sequence identity, R factor of ≤ 0.3, and resolution of ≤ 2.5 Å. Ideally, we would have restricted the search to membrane proteins only, but there were insufficient examples at high resolution, necessitating the use of a database that includes primarily water-soluble structures. All structures were re-refined by PDB-REDO (20). Using a method we developed previously (21), we searched for lysine-(carboxyl-carboxylate) triads and eliminated identical triads within oligomers belonging to the same structure. Next, we distinguished different lysine-(carboxyl-carboxylate) interaction geometries by performing hierarchical agglomerative clustering using the average-linkage approach and a 0.6 Å RMSD cutoff (22) on 7 atoms of each lysine-(carboxyl-carboxylate) triad. Specifically, the Lys NZ and terminal Asp or Glu sidechain atoms (Cγ-OD1-OD2 or Cδ-OE1-OE2) were used, taking care to classify Asp and Glu side chain oxygens based on their spatial positions instead of their crystallography-assigned atom names.

### IV. POLAR RESIDUE PROTONATION

**Figure S1.** To ensure structural stability in simulation Cα root-mean squared deviation (RMSD) from the starting NMR structure was calculated for the 74 core residues in each of the three helices of the DAP12-NKG2C complex. RMSD is calculated for each of the protonation states and mutation states of the receptor complex.

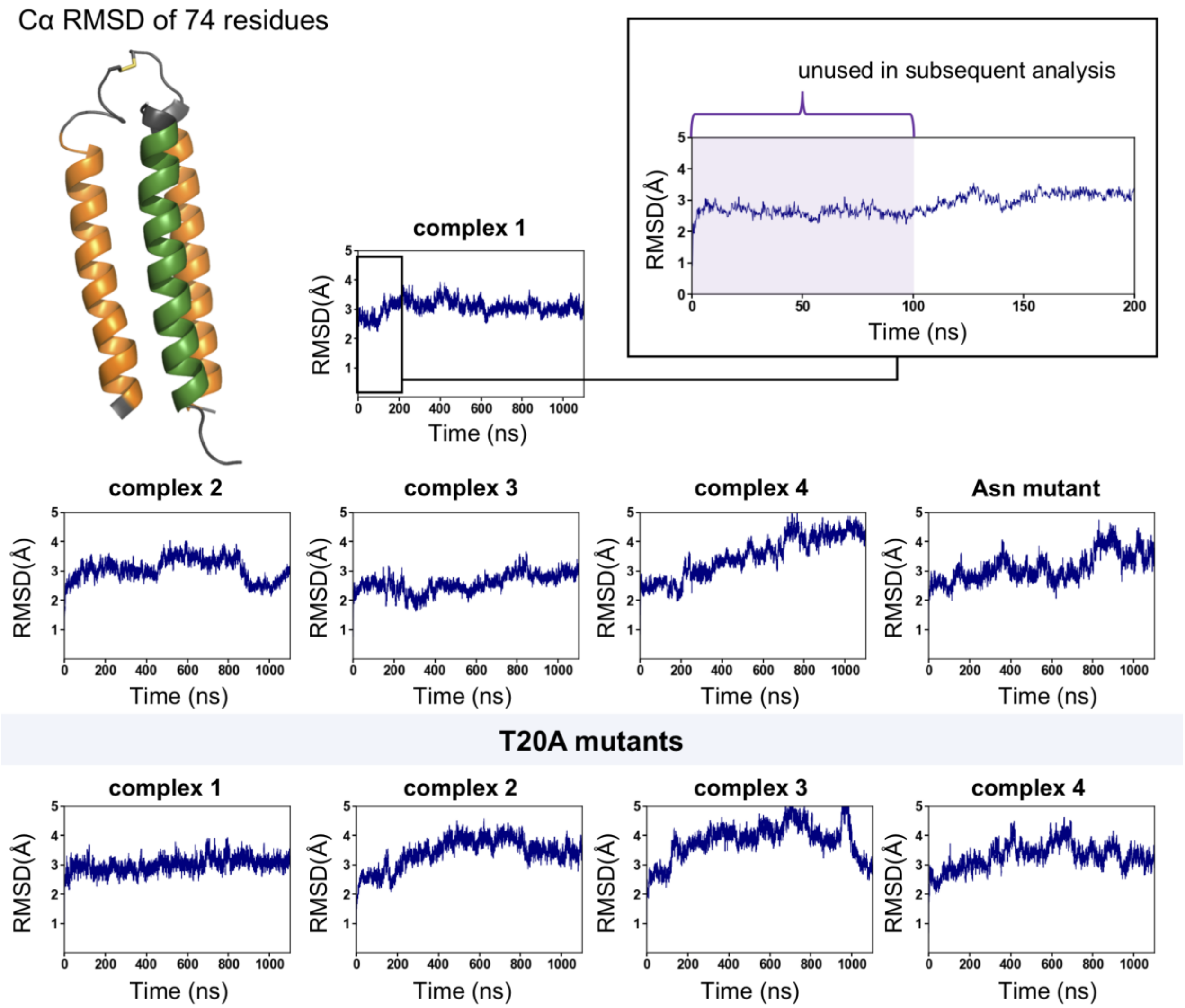

**Figure S2.** Quantification of polar residue dynamics as a function protonation state: complex 1 ($D^-D^-K^+$), complex 2 ($D^0D^-K^+$), complex 3 ($D^-D^0K^+$) and complex 4 ($D^0D^0K^0$). **A.** Principal component analysis of local polar residues. PC-2 vs. PC-1 plots with complex 2 as a point of comparison. **B.** Visual representation of the motions along PC-1 and PC-2. PC-1 accounts for the motion pushing lysine away from Asp1. This motion also brings the two Asp residues closer together. PC-2 accounts for the motion of lysine from Asp1 to Asp2. As lysine moves to Asp2, Asp1 is pushed away. PCA plots confirm that complex 2 ($D^0D^-K^+$) and complex 3 ($D^-D^0K^+$) are conformationally similar, but there is significantly more variance in the conformations explored by complex 1 ($D^-D^-K^+$) and complex 4 ($D^0D^0K^0$).

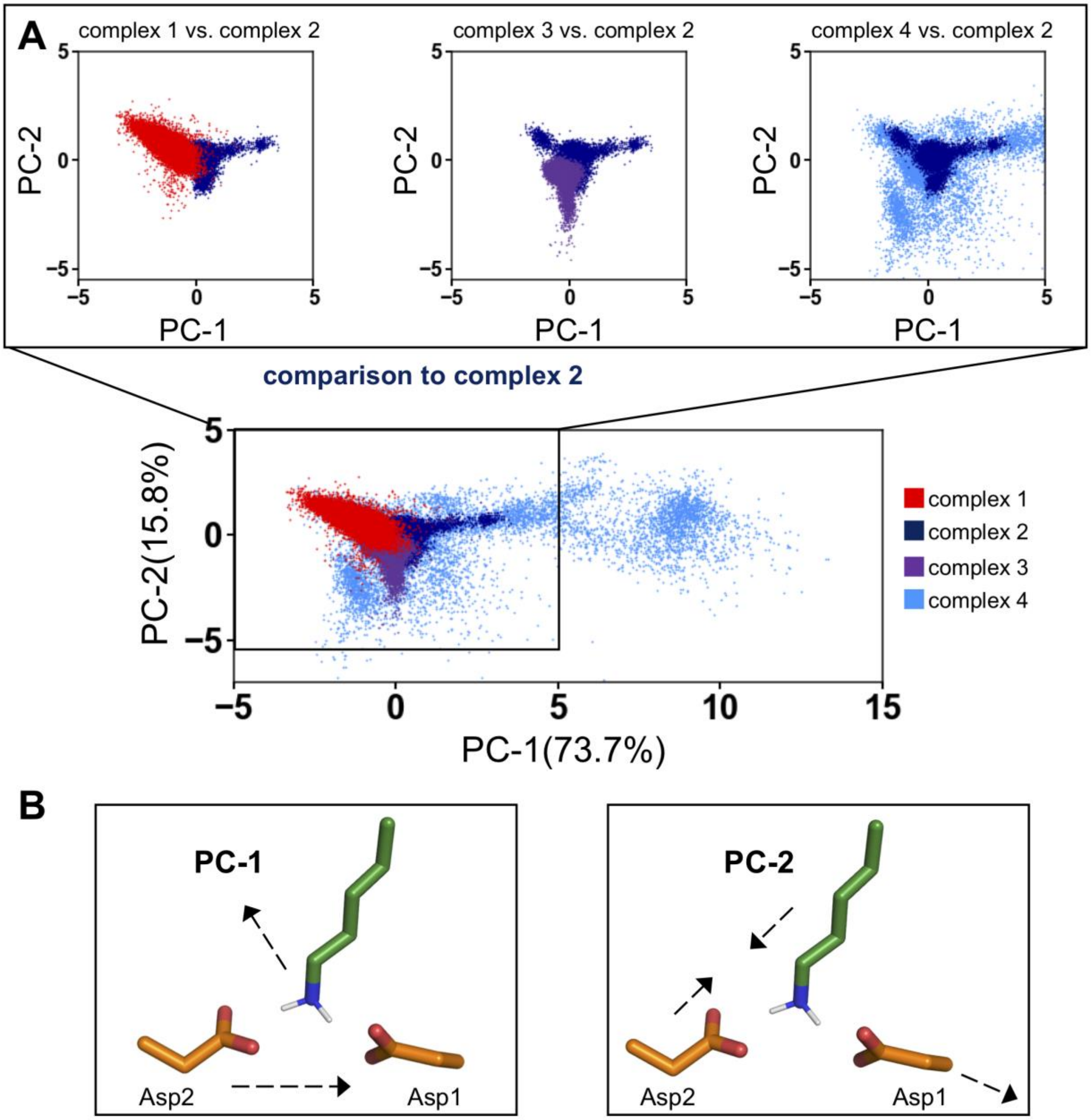

**Figure S3.** Comparison of cluster 1 centroids in different protonation states: complex 1 ($D^-D^-K^+$), complex 2 ($D^{\mathbf{0}}D^-K^+$), complex 3 ($D^-D^{\mathbf{0}}K^+$), and complex 4 ($D^{\mathbf{0}}D^{\mathbf{0}}K^{\mathbf{0}}$). Complex 2 is used as a reference.

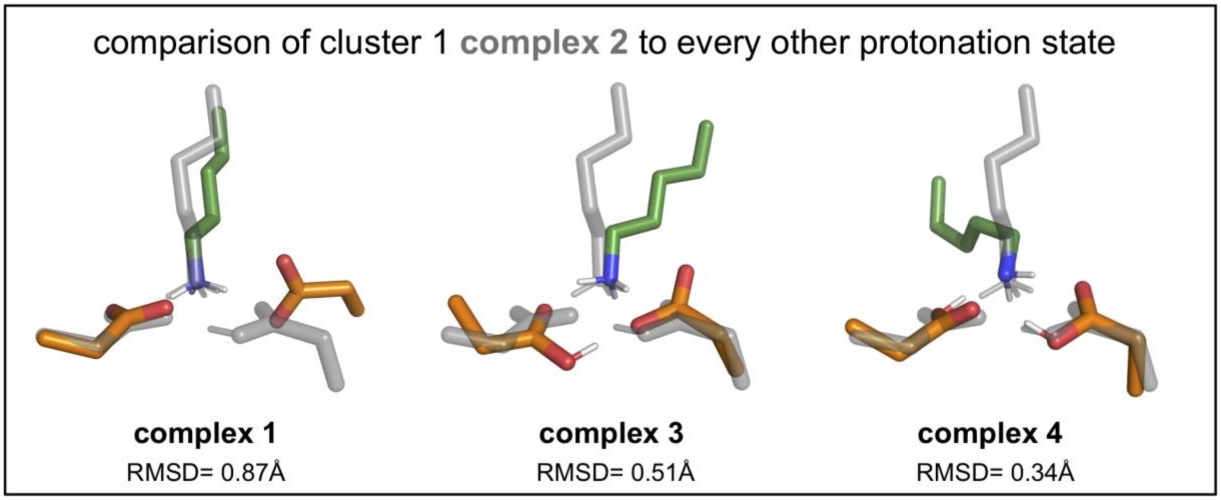


**Figure S4.** Complex 1 ($D^-D^-K^+$) conformational states. **A.** Polar residue cluster number versus time (left) and quantification of proportion of frames spent in each cluster (right). **B.** Structural centroids of the top four polar residue clusters.

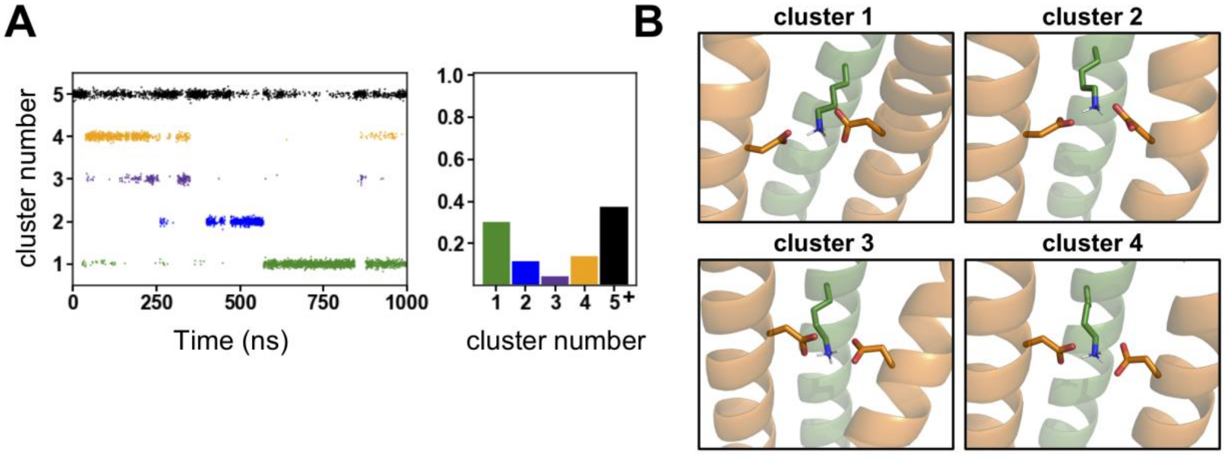

**Figure S5.** Contributions to polar group conformational instability for complex 1 (D⁻D⁻K⁺). **A.** Short-range nonbonded potential energy between aspartate residues for each protonation state of the wild-type complex. Electrostatic contributions are in red and van de Waals contributions are in gray. **B.** Minimum distance between aspartate oxygens as a function of simulation time. **C.** Quantification of the presence of a sodium ion in association with the polar residues. The proportion of time the ion spends associated with the polar residues is given as a function of simulation number. **D.** Quantification of polar group hydration. Number of waters within 3.4 Å of the Asp-Asp-Lys motif as a function of time are plotted for one simulation. **E.** Number of water molecules interacting with each of the polar residues. Plot shows the proportion of time a given number of waters are hydrogen bonded to each of the three polar residues (Lys, Asp1, Asp2).

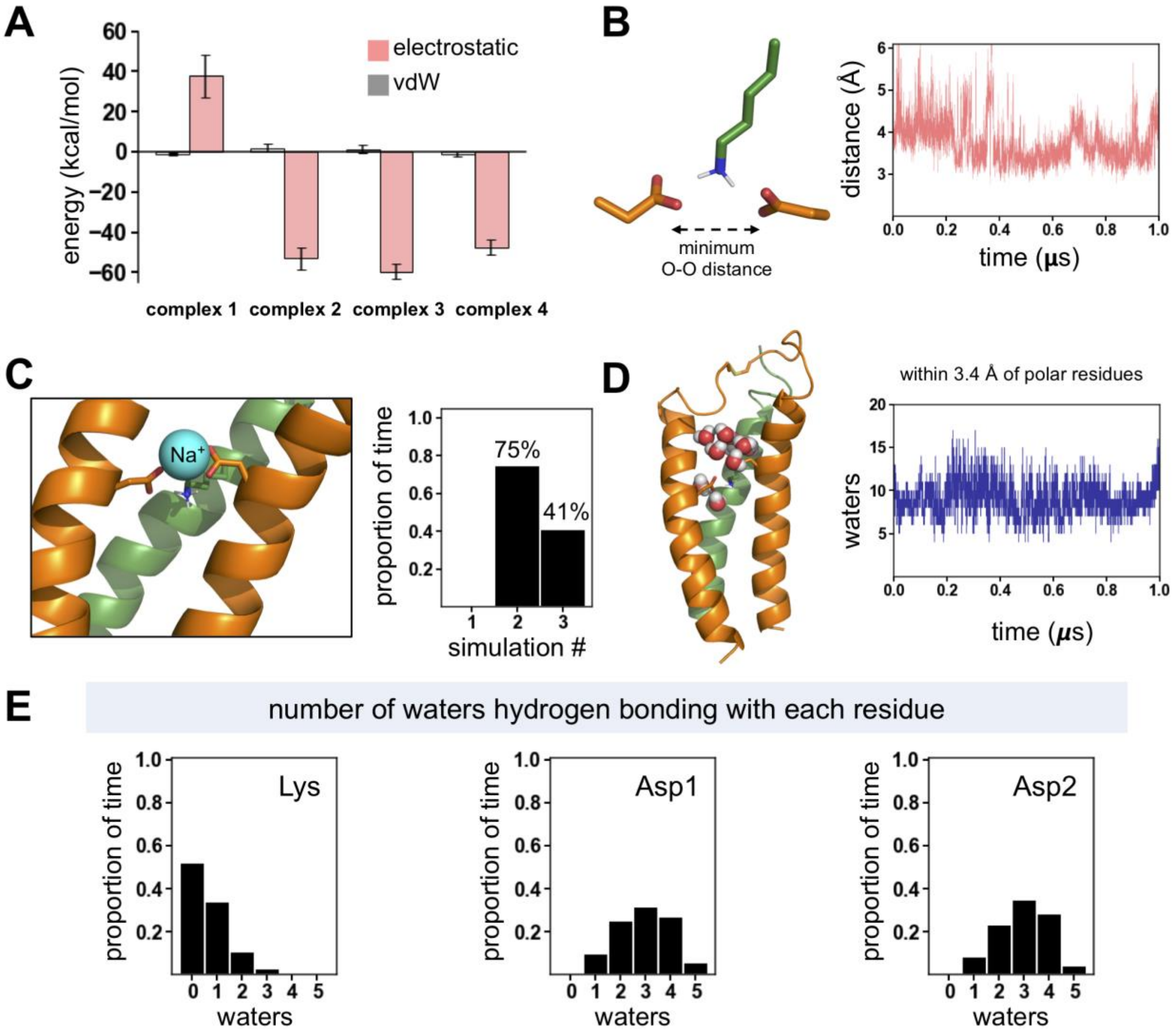

**Figure S6.** Evaluation of Asp-Lys association in complex1 ($D^{-}D^{-}K^{+}$) **A.** Minimum distance between aspartate oxygen and lysine nitrogen as a function of simulation time for Asp1 (left) and Asp2 (right). **B.** Quantification of lysine-aspartate hydrogen bond persistence given as the proportion of time spent hydrogen bonded in each of three simulations.

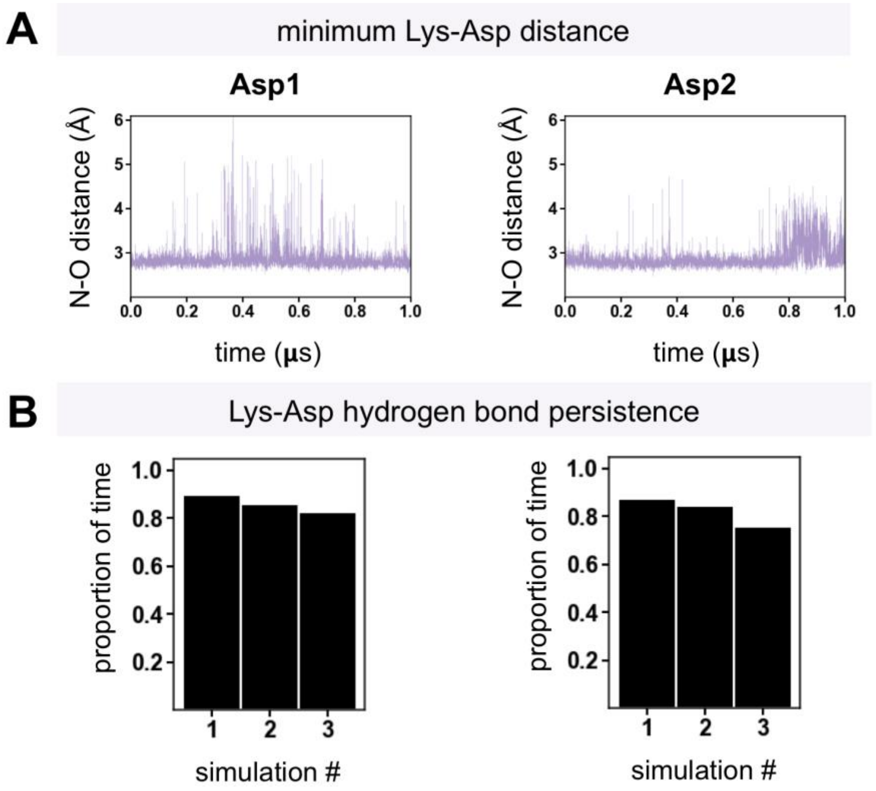

**Figure S7.** Analysis of carboxyl-carboxylate pair in **A.** complex 2 ($D^0D^-K^+$) and **B.** complex 3 ($D^-D^0K^+$). For each complex, minimum distance between aspartate oxygens as a function of simulation time (left). Quantification of aspartate-aspartic acid hydrogen bond persistence given as the proportion of time spent hydrogen bonded in each of three simulations (middle). Quantification of the position of the aspartate carboxylate relative to the syn-carboxyl it is hydrogen bonded to, measured based on the angle formed between the center of the hydrogen bond, and the two carboxylate oxygens. Angles above 130° designate an anti arrangement of the carboxylate relative to the syn-carboxyl (right).

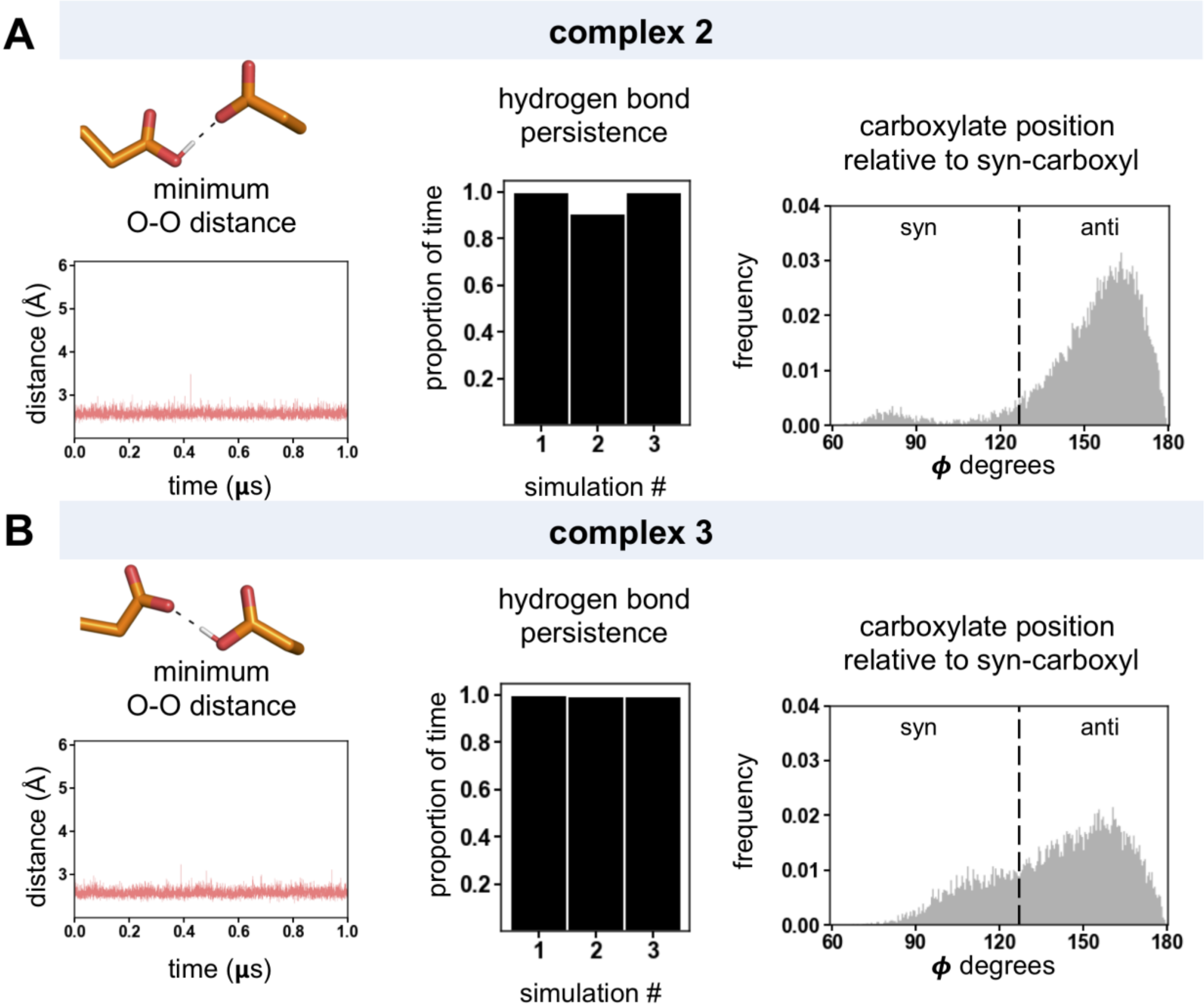

**Figure S8.** Simulation of the DAP12 homodimer alone in the protonation state of complex 2 ($D^0D^-$) **A.** Quantification of the proportion of time spent in the syn or anti arrangement of the aspartic acid carboxyl based on the dihedral angle formed between the OD1-Cγ-OD2-HD2 atoms. Angles around 180° designate an anti-arrangement of the carboxyl **B.** Quantification of aspartate-aspartic acid hydrogen bond persistence given as the proportion of time spent hydrogen bonded in each of three simulations.

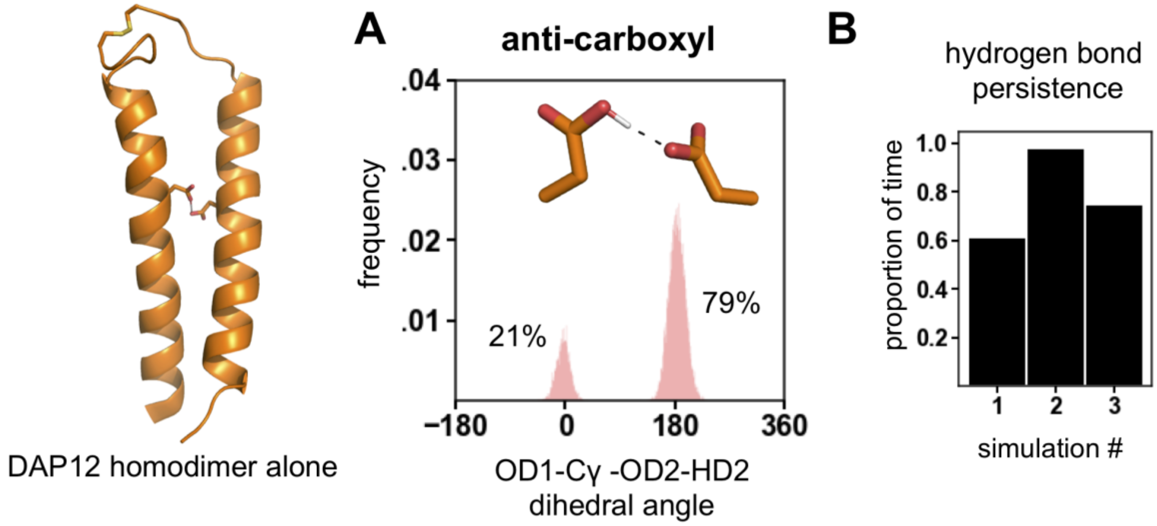

**Figure S9.** Quantification of polar group hydration. Number of waters within 3.4 Å of the Asp-Asp-Lys motif as a function of time are plotted for one simulation **A.** complex 2 ($D^0D^-K^+$) and **B.** complex 3 ($D^-D^0K^+$) **C.** Number of water molecules interacting with each of the polar residues. Plot shows the proportion of time a given number of waters are hydrogen bonded to each of the three polar residues (Lys, Asp1, Asp2) in complex 2 ($D^0D^-K^+$) (top) and complex 3 ($D^-D^0K^+$) (bottom).

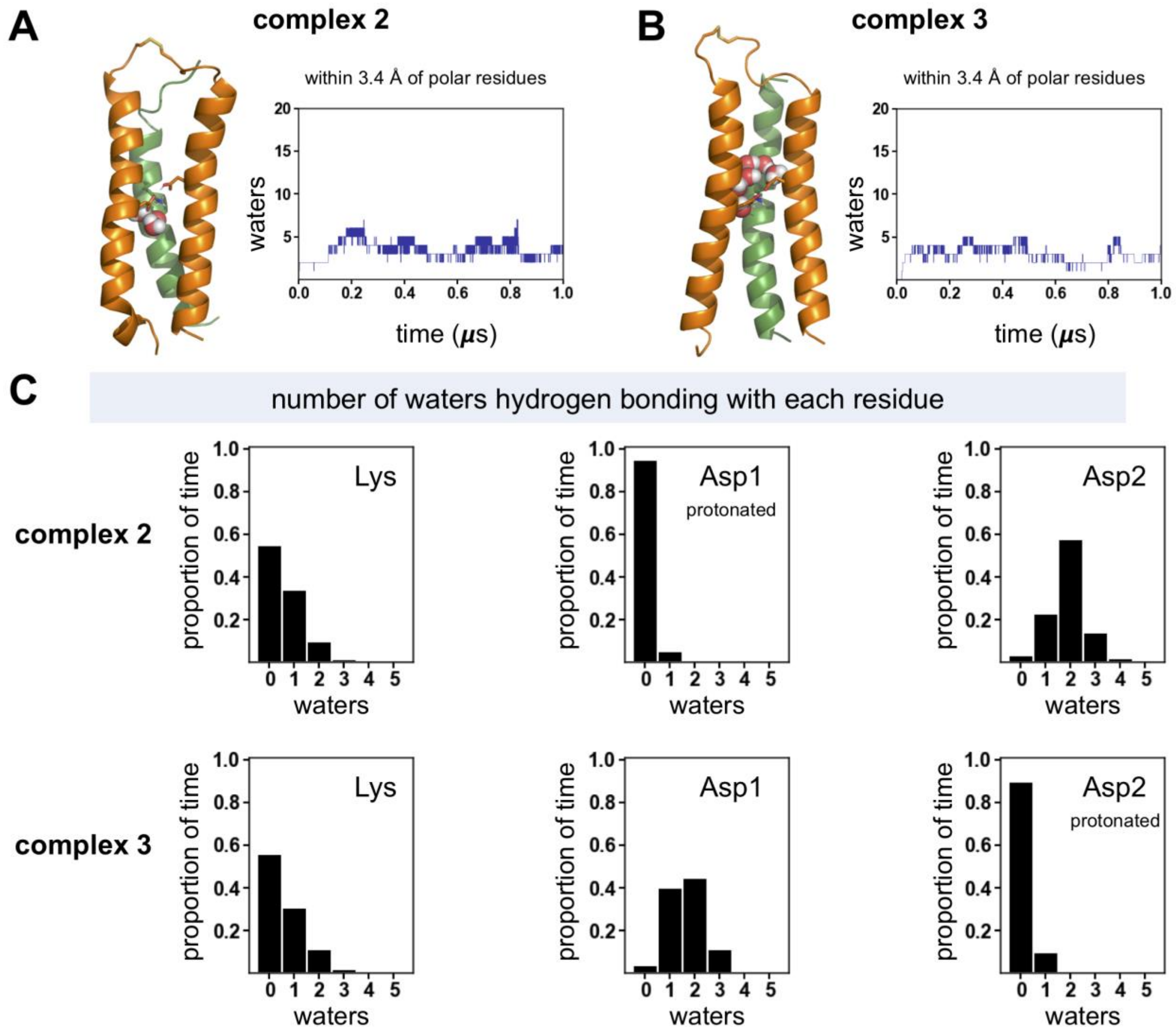

**Figure S10.** Evaluation of Asp-Lys association in **A.** complex 2 ($D^0D^-K^+$) **B.** complex 3 ($D^-D^0K^+$). For both complexes the panel shows the minimum distance between aspartate or aspartic acid oxygen and lysine nitrogen as a function of simulation time and quantification of lysine-aspartate hydrogen bond persistence given as the proportion of time spent hydrogen bonded in each of three simulations for Asp1 (left) and Asp2 (right).

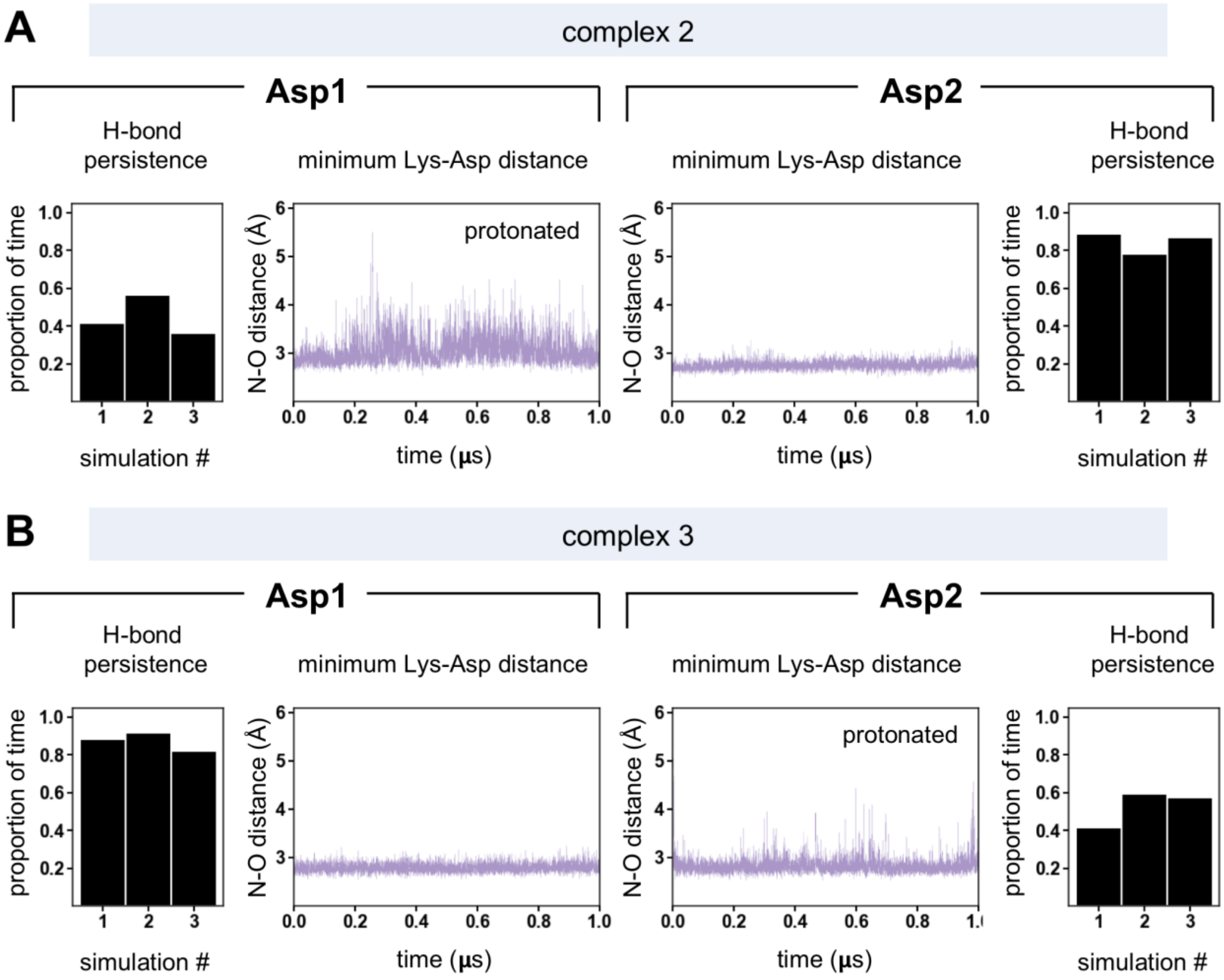

**Figure S11.** Analysis of carboxyl-carboxyl pair and its association with lysine in complex 4 ($D^0D^0K^0$) **A.** Hydrogen bond persistence given as the proportion of time spent hydrogen bonded in each of three simulations for Asp1-Asp2 (left), Asp1-Lys (middle) and Asp2-Lys (right) **B.** Minimum distance between aspartic acid oxygens as function of simulation time.

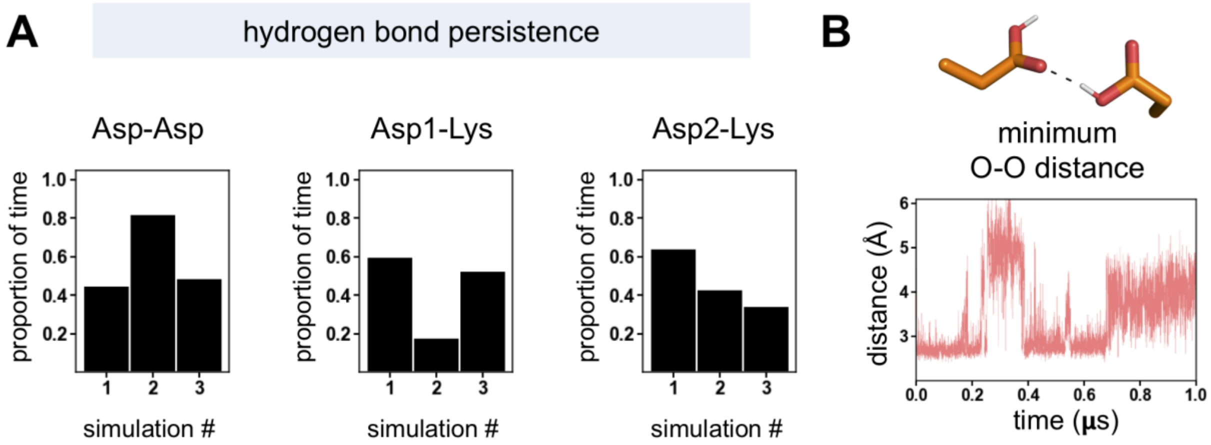

### V. DISCUSSION OF DFT CALCULATIONS

To estimate the strength of the hydrogen bonding interactions natural bond orbital (NBO) (23) calculations were performed with the Gaussian09 package (24). The donor-acceptor interaction energies for any pair of donor and acceptor heavy atoms were summed to evaluate the overall strength of the hydrogen bond (**Figure S12**) (25). These values were confirmed to be reasonable by their similarity to the interaction energy of the complex. The interaction energy was defined as the difference in the electronic energy between the entire complex and that of the individual elements considered (Lys, Asp + 2 $H_2O$, AspH/Asn + Thr; **Table S2**). These elements were chosen so as to primarily consider the energy gained from hydrogen bonding of the polar triad, rather than with waters and the backbone.

**Figure S12.** Comparison of polar group interaction energies between wild type complex 2 ($D^0D^-K^+$) and the Asn mutant ($N^0D^-K^+$). **A.** Stabilizing NBO interaction energies identified in the relaxed structure of cluster 1 for complex 2. **B**. Stabilizing NBO interaction energies identified in the relaxed structure of cluster 1 for the Asn mutant. In both **A** and **B**, dashed lines indicate the donor acceptor pair for the NBO interaction with the values being the interaction magnitude in kcal/mol. In both cases, interactions between the Asp and the waters were ignored because these were considered as one unit for the calculation of interaction energies (**Table S2**). **C**. Comparison of the cluster 1 centroid for the asparagine mutant from the MD simulations (gray) to the DFT optimized structure (colored). These reveal a highly similar geometry with the RMSD for the clustered atoms of 0.11 Å and for all of the relaxed heavy atoms of 0.15 Å.

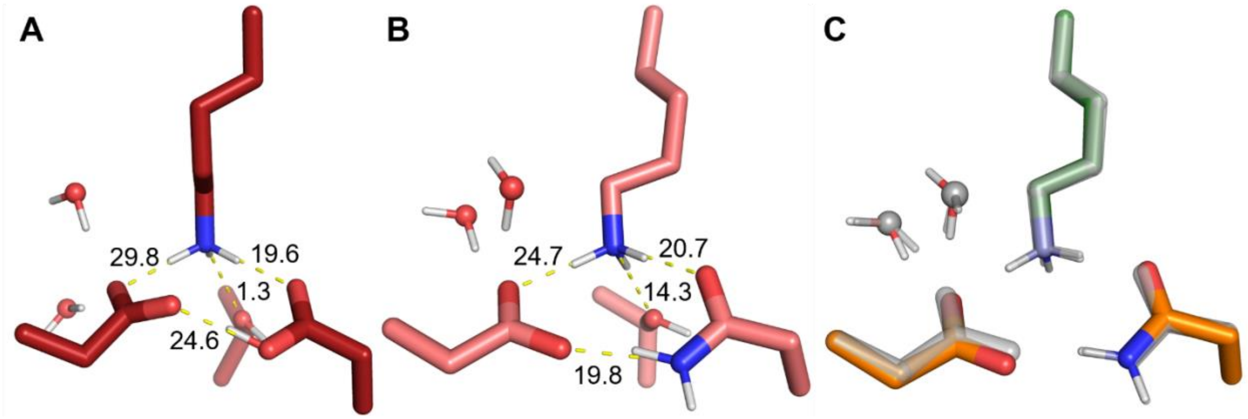

**Table S2.** The electronic energy of each component of the clusters, the sum of those energies, the energy for the full complex and the interaction energy. The interaction energy is defined as the difference in energy between the full complex and the sum of the component pieces and is given in kcal/mol. All other energies are given in Hartrees.

| **Component** | **Cluster 1, Complex 2** | **Cluster 1, Asn Mutant** |
|---|---|---|
| **Lys** | -591.2002 | -532.0012 |
| **Asp + 2$H_2O$** | -758.5579 | -758.5692 |
| **Thr + AspH/Asn** | -1138.1042 | -1118.2340 |
| **Total of Components** | -2487.8623 | -2468.0014 |
| **Full** | -2487.986 | -2468.116 |
| **Difference** | 77.8 | 71.8 |

The plausibility of proton transfer reactions between $D_{16}$ and $D_{49}$ in cluster 1 of complex 2 was initially evaluated via a relaxed surface scan. In this scan, the backbones were held fixed, the $O_{Asp}$-H distance was systematically varied, and the rest of the molecule was allowed to relax. In the presence of water, the relaxed surface scan indicated that the energy of the complex increased up to a plateau of approximately +5 kcal/mol as the proton was transferred (**Figure S13**). This suggests that solvent rearrangement is the primary barrier to proton transfer, which is consistent with our expectation that the newly formed Asp species would require hydrogen bond donors to help stabilize it.

**Figure S13**. Comparison of the energy profile for the relaxed surface scans that connect complex 2 and complex 3 via proton transfer. Inclusion of two explicit waters leads to no distinct maxima, whereas without water a clear maximum is observed.

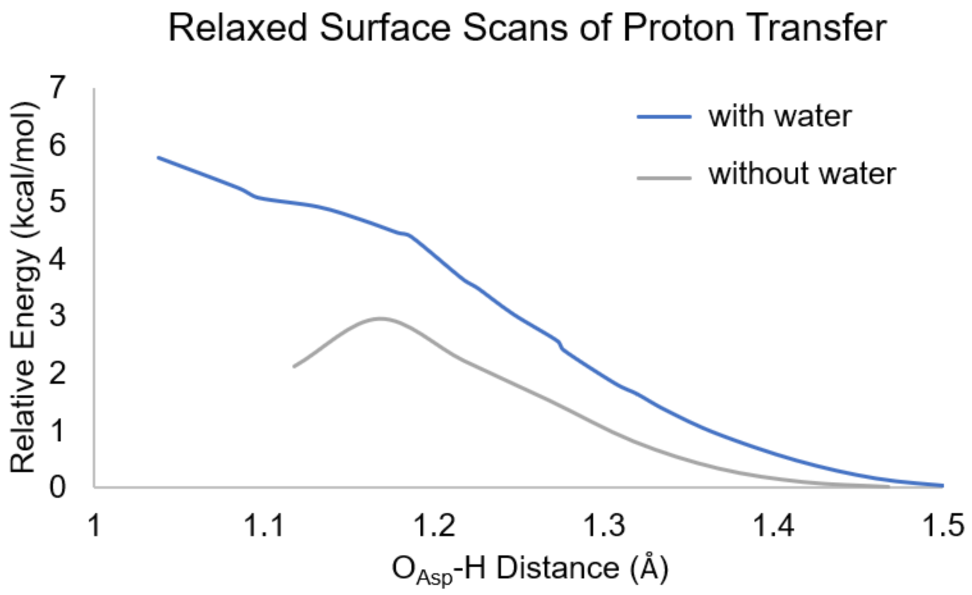

To further confirm that the solvent rearrangement was the primary barrier to proton transfer, the structure was relaxed in the absence of water. This led to a much shorter O–O distance for the carboxyl-carboxylate pair consistent with their improved p$K_a$ match (26). Then, the relaxed surface scan for proton transfer was again employed to investigate the reaction pathway. In this case, clear minima were observed at each endpoint (**Figure S13**). Each of these endpoints was optimized and used as input for a nudged elastic band-transition state calculation (27). This calculation evaluates both the minimum energy pathway and subsequently minimizes the transition state (**Figure S14**). A frequency calculation confirmed that the transition state had only one large imaginary frequency ($-477$ cm$^{-1}$) and that the barrier to proton transfer is minimal ($\Delta G^{\ddagger}$ = +2 kcal/mol). To confirm that the transition state was correctly identified an intrinsic reaction coordinate (IRC) calculation was performed (28). This calculation follows the transition state frequency in both directions to identify the reactants and products. As expected, the IRC confirms a proton transfer reaction (**Figure S15**). The small barrier to proton transfer in the absence of explicit waters is consistent with solvent rearrangement being the primary barrier to proton transfer.

**Figure S14.** The minimum energy pathway for proton transfer without explicit waters as calculated by the nudged elastic band method.

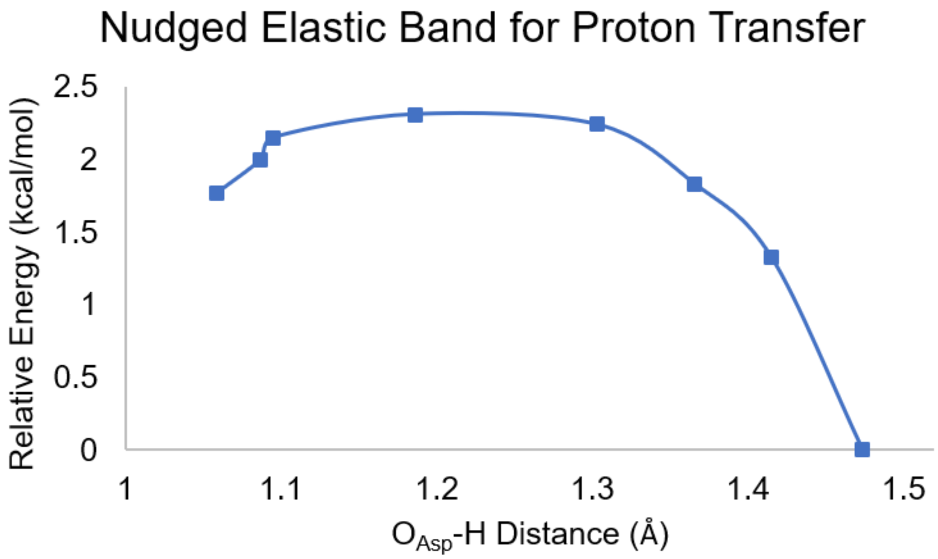

**Figure S15.** The transition state mode is followed in both the forward and backward direction to reveal the corresponding reactants and the products. As expected, the transition state describes the transfer of the proton between the two aspartate residues (i.e., the interconversion of complex2 ($D^{0}D^{-}K^{+}$) and complex3 ($D^{-}D^{0}K^{+}$)).

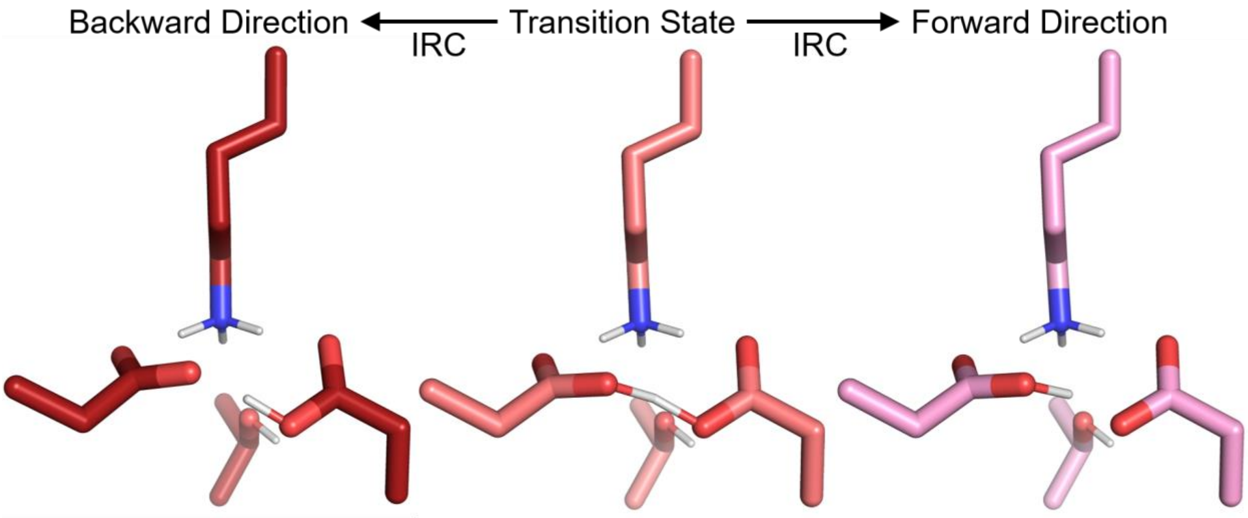

### VI. PDB CLUSTERS

**Figure S16.** The top eight clusters from the PDB search highlighting the number of members in each cluster and the carboxyl-carboxylate arrangement.

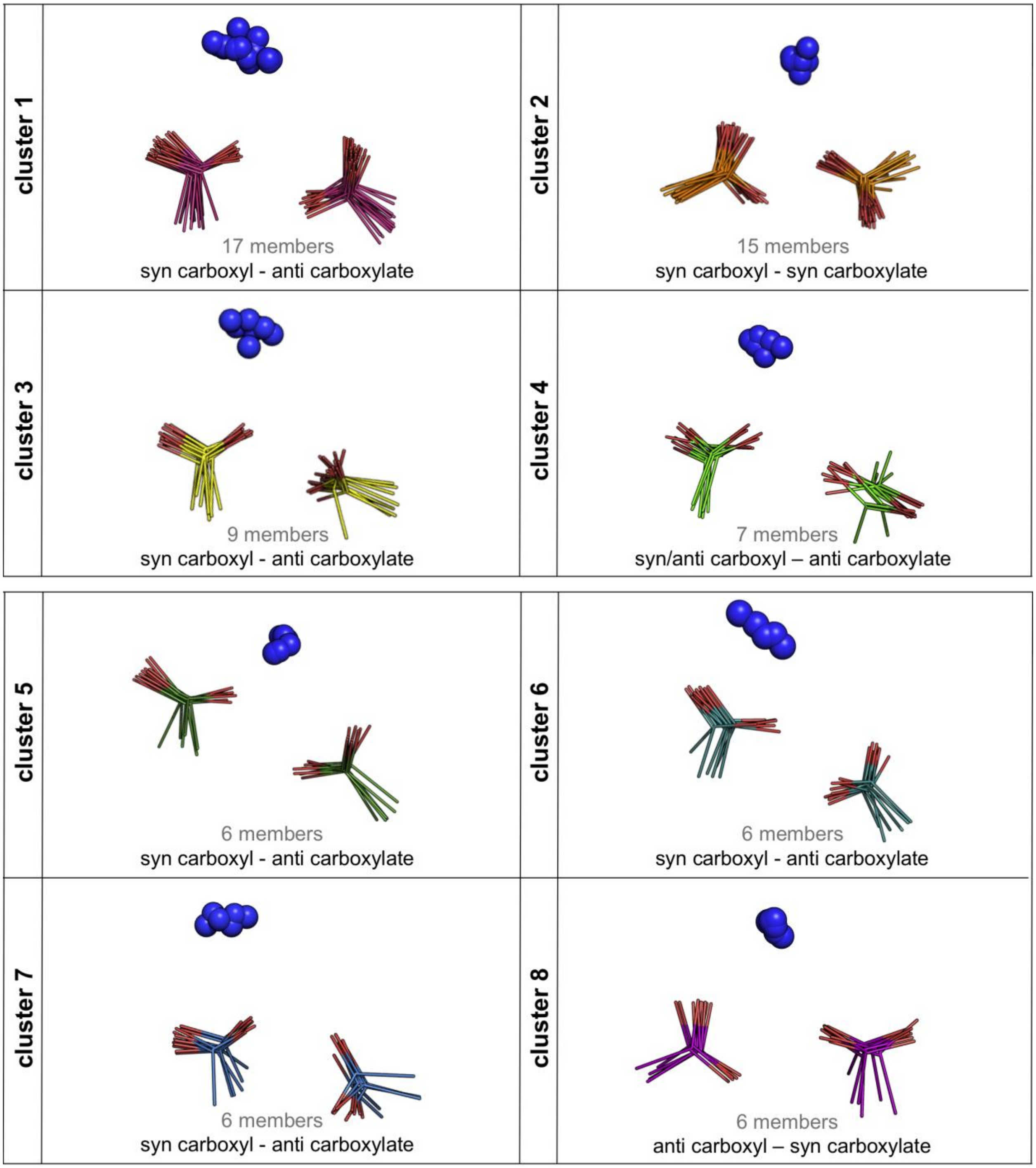

**Table S3.** PDB search counting statistics. From the non-redundant set of 13,026 protein crystal structures, 1,770 proteins had Glu/Asp carboxylate pairs with oxygen-oxygen distances less than 3.0Å. There were 2,492 carboxylate pairs in this set. Only the 1,583 pairs that had oxygen-oxygen distances less than 2.8 Å were identified as carboxyl-carboxylate pairs based on the close approach of the oxygen atoms which would be difficult if both carboxylates were deprotonated. Within this set of carboxyl-carboxylate pairs, 140 pairs interact with lysine NZ with an oxygen-nitrogen distance of less than 3.5 Å. A total of 9% of carboxyl-carboxylate pairs form salt bridges to Lys residues, based on this criterion. For comparison, a total of 11.2% (8439 - 140) of the Glu/Asp residues that are not engaged in carboxyl-carboxylate pairs form salt bridges with Lys residues. Thus, there is not a large difference in the propensity for Lys to interact with single Glu/Asp residues versus carboxyl-carboxylate pairs of Glu/Asp residues in proteins, which is consistent with the expectation that both have a net charge of -1.

| **Category** | **Count** |
|---|---|
| **Protein crystal structures used** | 13,026 |
| **Glu/Asp carboxylate pairs with O-O distance < 3.0Å** | 2,492 (within 1,770 proteins) |
| **Carboxyl-carboxylate pairs short O-O distance < 2.8Å** | 1,583 |
| **Lys-(carboxyl-carboxylate) with O-N distance < 3.5Å** | 140 |

### VII. EFFECT OF MUTATIONS ON POLAR RESIDUES

**Figure S17.** Effect of Asn mutation on polar triad arrangement. **A.** Six atom root-mean squared deviation (RMSD) comparison of cluster 1 centroid of Asn mutant ($N^0D^-K^+$) to cluster 1 centroid of wild-type complex 2 ($D^0D^-K^+$). **B.** Minimum distance between asparagine nitrogen and aspartate oxygen as function of simulation time. **C.** Asparagine-aspartate hydrogen bond persistence given as the proportion of time spent hydrogen bonded in each of three simulations. **D.** Structural centroids of polar residue clusters 2-4 and the amount of time spent in each of those geometric arrangements.

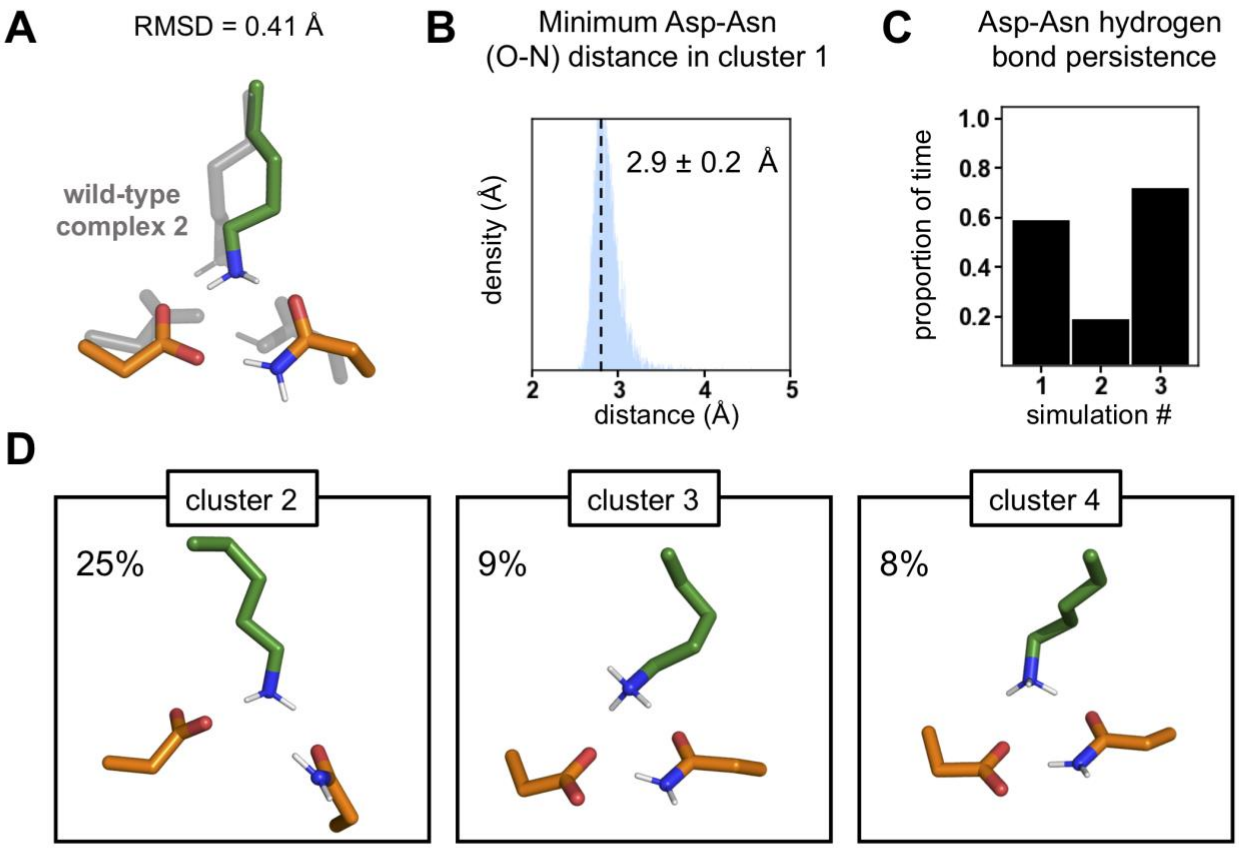

**Figure S18.** The role of threonine in DAP12-NKG2C assembly. **A.** Quantification of threonine-aspartate hydrogen bond persistence for each of the four protonation states complex 1 ($D^{-}D^{-}K^{+}$), complex 2 ($D^{0}D^{-}K^{+}$), complex 3 ($D^{-}D^{0}K^{+}$), and complex 4 ($D^{0}D^{0}K^{0}$). Values are given as the proportion of time spent hydrogen bonded out of 3 **μ**s for DAP12 monomer 1 (left) and monomer 2 (right) in each case. **B.** Quantification of threonine-lysine hydrogen bond persistence given as the proportion of time spent hydrogen bonded in each of three simulations for each of the four protonation states.

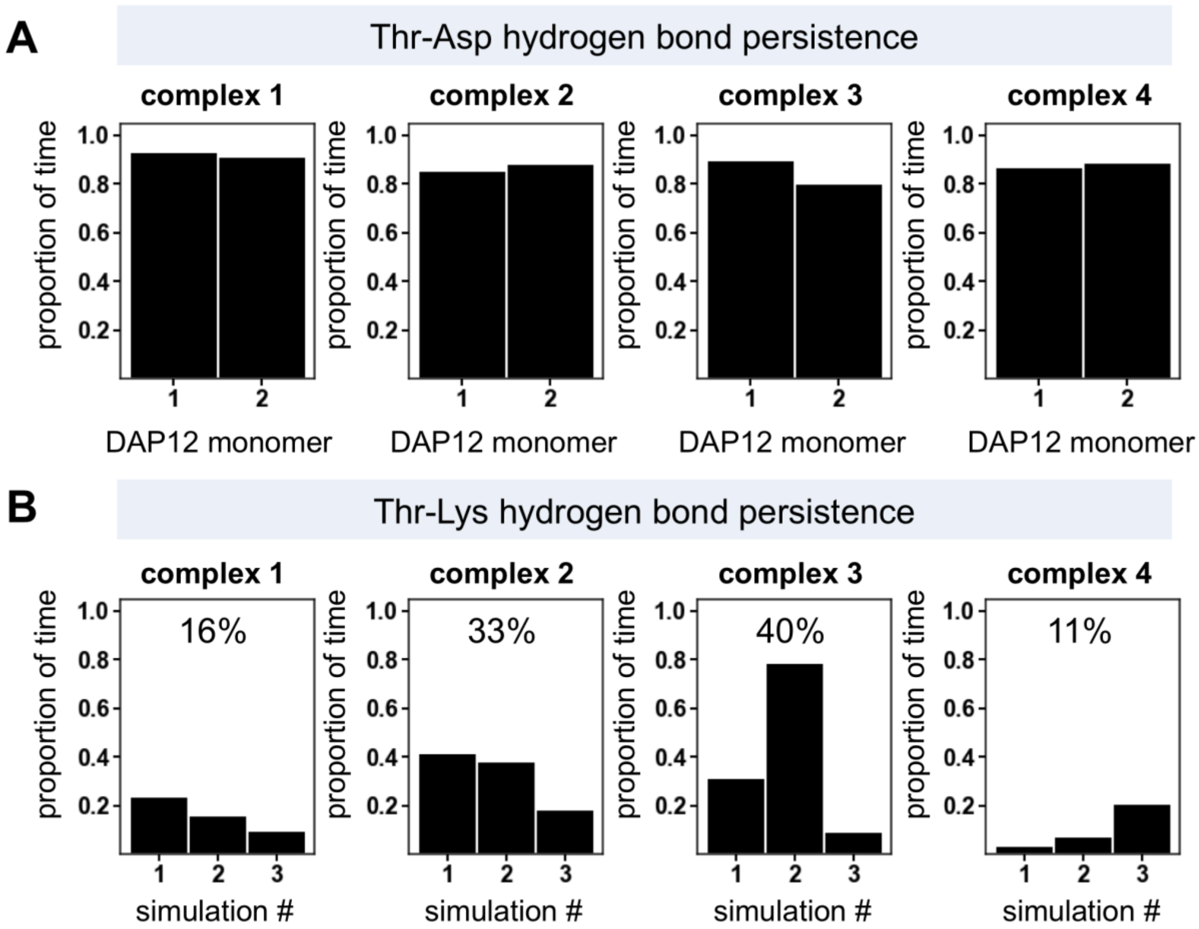

**Figure S19.** The effect of T20A mutation on polar triad conformational stability and hydration. **A.** Comparison of the centroid of cluster 1 (top) and cluster 2 (bottom) of T20A mutants to the centroid of cluster 1 wild type (gray) for each of the four protonation states: complex 1 ($D^{-}D^{-}K^{+}$), complex 2 ($D^{0}D^{-}K^{+}$), complex 3 ($D^{-}D^{0}K^{+}$), and complex 4 ($D^{0}D^{0}K^{0}$). **B.** Number of water molecules interacting with lysine for both wild type (gray) and T20A mutants (black) as a function of protonation state. Plots show the proportion of time a given number of waters is hydrogen bonded to lysine. In the case of complex 2 ($D^{0}D^{-}K^{+}$) and complex 3 ($D^{-}D^{0}K^{+}$), T20A mutation causes an increase in the number of waters that hydrogen bond lysine, highlighting the importance of the threonine residue in these protonation states.

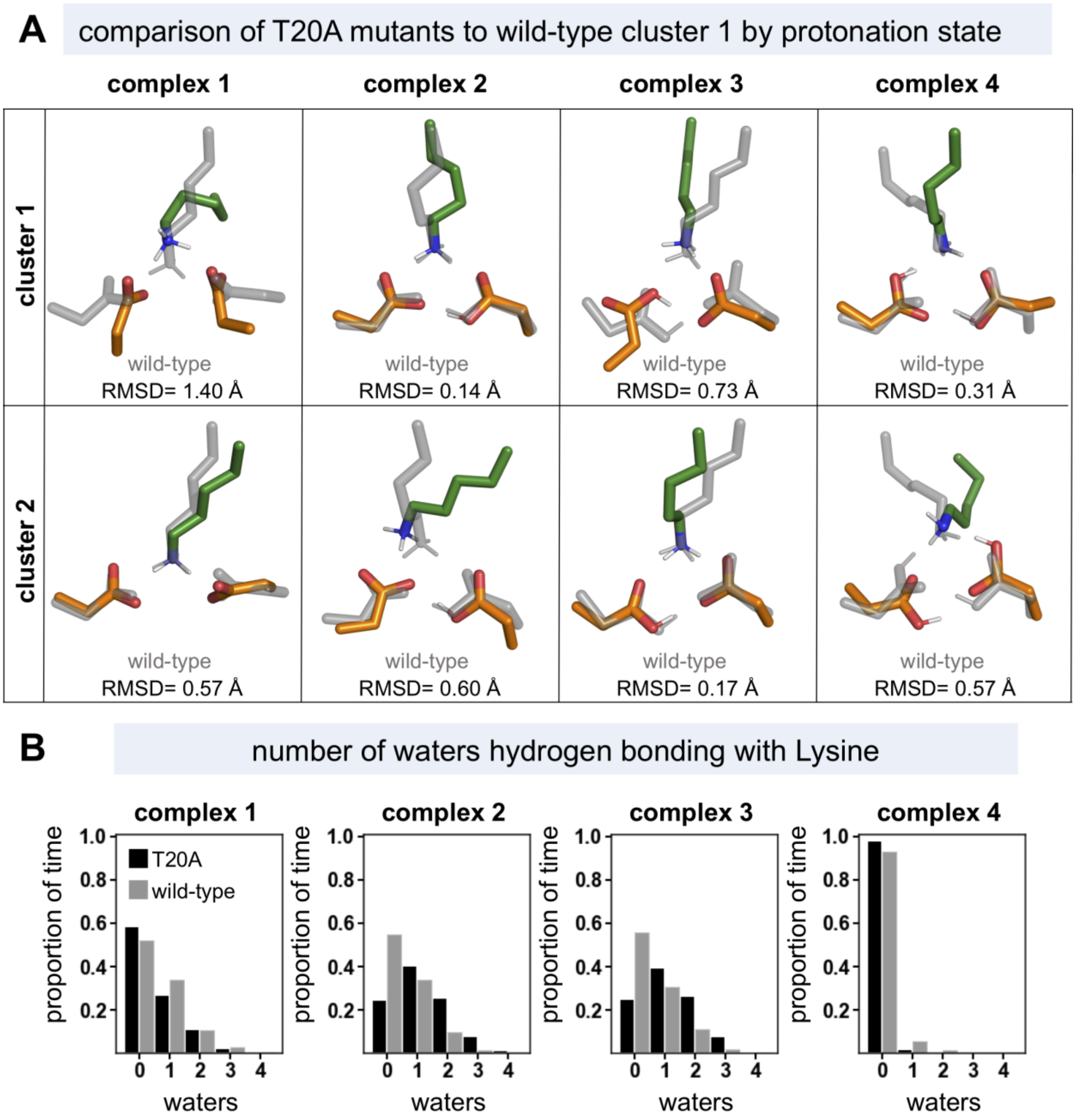

### VIII. HELIX DYNAMICS

**Figure S20.** Complex 2 ($D^0D^-K^+$) 5 µs of simulation **A.** Quantification of proportion of frames spent in each of the polar residue (Asp-Asp-Lys) clusters highlighting the percent of time spent in cluster 1 (left) and structural centroid of cluster 1 compared to the centroid of cluster 1 for 3 µs of simulation (right) **B.** 22 residue Cα root-mean squared deviation (RMSD) from cluster 1 centroid for helix pairs within the trimer (DAP12 homodimer, DAP12-1 NKG2C, DAP12-2 NKG2C). The residues used were the two polar residues in the helix pair and the 10 surrounding each of these residues.

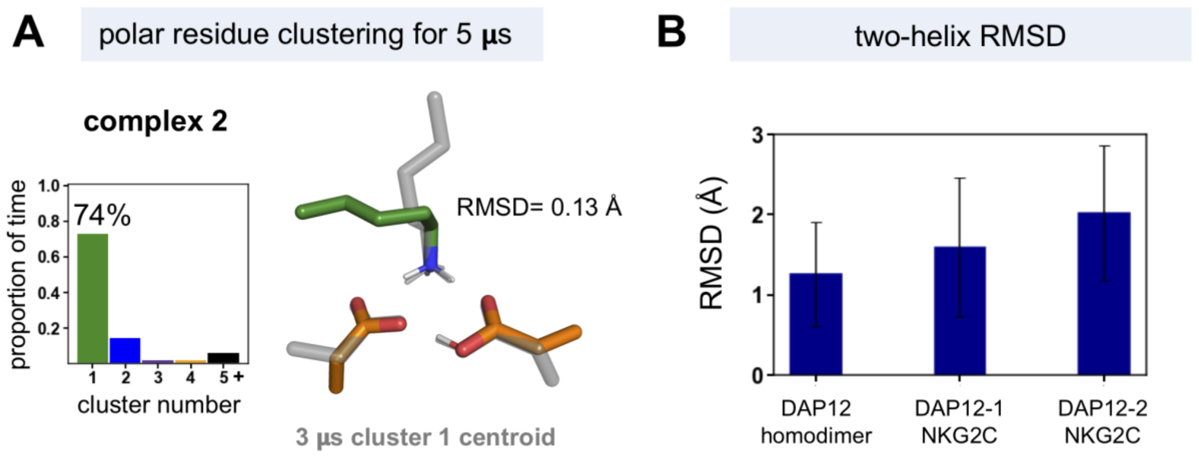

### IX. REFERENCES


1. M. E. Call, K. W. Wucherpfennig, J. J. Chou, The structural basis for intramembrane assembly of an activating immunoreceptor complex. *Nat. Immunol.* 11, 1023-1029 (2010).
2. S. Jo, T. Kim, W. Im, Automated builder and database of protein/membrane complexes for molecular dynamics simulations. *PloS one* 2, e880-e880 (2007).
3. D.A. Case *et al.* AMBER 2016. University of California, San Francisco (2016).
4. J. A. Maier *et al.*, ff14SB: Improving the accuracy of protein side chain and backbone parameters from ff99SB. *J. Chem. Theory Comput.* 11, 3696-3713 (2015).
5. C. J. Dickson *et al.*, Lipid14: The Amber lipid force field. *J. Chem. Theory Comput.* 10, 865-879 (2014).
6. R. Salomon-Ferrer, A. W. Gotz, D. Poole, S. Le Grand, R. C. Walker, Routine microsecond molecular dynamics simulations with AMBER on GPUs. 2. explicit solvent particle mesh Ewald. *J. Chem. Theory Comput.* 9, 3878-3888 (2013).
7. S. Miyamoto, P. A. Kollman, Settle: An analytical version of the SHAKE and RATTLE algorithm for rigid water models. *J. Comput. Chem.* 13, 952-962 (1992).
8. J. Shao, S. W. Tanner, N. Thompson, T. E. Cheatham, Clustering molecular dynamics trajectories: 1. characterizing the performance of different clustering algorithms. *J. Chem. Theory Comput.* 3, 2312-2334 (2007).
9. D. R. Roe, C. Bergonzo, T. E. Cheatham, 3rd, Evaluation of enhanced sampling provided by accelerated molecular dynamics with Hamiltonian replica exchange methods. *J. Phys. Chem. B* 118, 3543-3552 (2014).
10. D. R. Roe, T. E. Cheatham, 3rd, PTRAJ and CPPTRAJ: Software for Processing and Analysis of Molecular Dynamics Trajectory Data. *J. Chem. Theory Comput.* 9, 3084-3095 (2013).
11. F. Neese, Software update: the ORCA program system, version 4.0. *WIREs Computational Molecular Science* 8, e1327 (2018).
12. A. D. Becke, Density-functional exchange-energy approximation with correct asymptotic behavior. *Phys. Rev. A* 38, 3098-3100 (1988).
13. F. Weigend, R. Ahlrichs, Balanced basis sets of split valence, triple zeta valence and quadruple zeta valence quality for H to Rn: Design and assessment of accuracy. *Phys. Chem. Chem. Phys.* 7, 3297-3305 (2005).
14. F. Weigend, Accurate Coulomb-fitting basis sets for H to Rn. *Phys. Chem. Chem. Phys.* 8, 1057-1065 (2006).
15. S. Grimme, J. Antony, S. Ehrlich, H. Krieg, A consistent and accurate ab initio parametrization of density functional dispersion correction (DFT-D) for the 94 elements H-Pu. *J. Chem. Phys.* 132, 154104 (2010).
16. V. Barone, M. Cossi, Quantum Calculation of Molecular energies and energy gradients in solution by a conductor solvent model. *J. Phys. Chem. A* 102, 1995-2001 (1998).
17. W. Huang, D. G. Levitt, Theoretical calculation of the dielectric constant of a bilayer membrane. *Biophys. J.* 17, 111-128 (1977).
18. H. M. Berman *et al.*, The Protein Data Bank. *Nucleic Acids Res.* 28, 235-242 (2000).
19. G. Wang, R. L. Dunbrack, Jr., PISCES: a protein sequence culling server. *Bioinformatics* 19, 1589-1591 (2003).

20. R. P. Joosten, F. Long, G. N. Murshudov, A. Perrakis, The PDB_REDO server for macromolecular structure model optimization. *IUCrJ* 1, 213-220 (2014).
21. S. K. Tan *et al.*, Modulating integrin alphaIIbbeta3 activity through mutagenesis of allosterically regulated intersubunit contacts. *Biochemistry* 58, 3251-3259 (2019).
22. P. Virtanen *et al.*, SciPy 1.0: fundamental algorithms for scientific computing in Python. *Nature Methods* 17, 261-272 (2020).
23. E. D. Glendening, A. E. Reed, J.E. Carpenter, F. Weinhold, NBO Version 3.1. Gaussian Inc., Pittsburgh (2003).
24. M. J. Frisch *et al.*, Gaussian 09, Revision E.01 (2015).
25. A. E. Reed, L. A. Curtiss, F. Weinhold, Intermolecular interactions from a natural bond orbital, donor-acceptor viewpoint. *Chem. Rev.* 88, 899-926 (1988).
26. P. A. Sigala *et al.*, Determination of hydrogen bond structure in water versus aprotic environments to test the relationship between length and stability. *J. Am. Chem. Soc.* 137, 5730-5740 (2015).
27. A. V. Ivanov, D. Dagbartsson, J. Tranchida, V. M. Uzdin, H. Jonsson, Efficient optimization method for finding minimum energy paths of magnetic transitions. *J. Phys. Condens. Matter* 32, 345901 (2020).
28. K. Ishida, K. Morokuma, A. Komornicki, The intrinsic reaction coordinate. An ab initio calculation for HNC→HCN and H−+CH4→CH4+H− *J. Chem. Phys.* 66, 2153-2156 (1977).
