## Supplementary material for "Elucidation of the molecular interactions that enable stable assembly and structural diversity in multicomponent immune receptors": SI PDB Cluster Codes

| Cluster Number | PDB | Residues |
|---|---|---|
| 1 | 2ww8 | Glu54-Chain A, Glu103-Chain A, Lys731-Chain A |
| | 6o6y | Lys231-Chain A, Glu250-Chain A, Glu422-Chain A |
| | 5gw9 | Lys24-Chain A, Asp28-Chain A, Asp110-Chain A |
| | 3oyv | Lys116-Chain A, Glu137-Chain A, Asp147-Chain A |
| | 5xwi | Glu279-Chain A, Lys330-Chain A, Asp334-Chain A |
| | 5k4b | Lys248-Chain A, Glu350-Chain A, Glu359-Chain B |
| | 4q8k | Glu188-Chain A, Glu221-Chain A, Lys349-Chain A |
| | 5ziq | Lys129-Chain B, Glu186-Chain B, Glu190-Chain B |
| | 4xxl | Glu72-Chain A, Lys108-Chain A, Asp112-Chain A |
| | 6os7 | Lys183-Chain A, Glu200-Chain A, Glu357-Chain B |
| | 3drf | Asp504-Chain A, Lys525-Chain A, Asp529-Chain A |
| | 3l1n | Lys30-Chain A, Asp34-Chain A, Asp49-Chain A |
| | 3k6o | Asp119-Chain A, Lys193-Chain B, Asp195-Chain B |
| | 6sa9 | Asp55-Chain A, Lys31-Chain B, Asp35-Chain B |
| | 5dl8 | Lys300-Chain B, Glu303-Chain B, Glu343-Chain B |
| | 3pty | Lys747-Chain A, Asp754-Chain A, Asp1051-Chain A |
| | 2pd2 | Glu33-Chain B, Glu94-Chain B, Lys98-Chain B |
| 2 | 2o16 | Asp134-Chain A, Lys63-Chain B, Asp134-Chain B |
| | 2zod | Asp136-Chain A, Lys140-Chain A, Asp136-Chain B |
| | 3cra | Lys119-Chain A, Glu122-Chain A, Asp61-Chain B |
| | 6lce | Asp129-Chain A, Lys134-Chain A, Glu151-Chain A |
| | 3cay | Glu7-Chain C, Glu18-Chain J, Lys22-Chain J |
| | 5xwi | Glu58-Chain A, Lys234-Chain A, Glu309-Chain A |
| | 5x93 | Glu299-Chain A, Lys303-Chain A, Glu1021-Chain A |
| | 3m6z | Asp31-Chain A, Glu123-Chain A, Lys282-Chain A |
| | 3lp5 | Lys73-Chain A, Asp77-Chain A, Glu153-Chain A |
| | 4u9v | Lys97-Chain B, Glu100-Chain B, Glu139-Chain B |
| | 2xu3 | Lys17-Chain A, Glu50-Chain A, Glu86-Chain A |
| | 3ot2 | Glu28-Chain A, Lys123-Chain A, Asp159-Chain B |
| | 5dwa | Glu173-Chain A, Asp223-Chain A, Lys224-Chain A |
| | 3m6z | Asp31-Chain B, Glu123-Chain B, Lys282-Chain B |
| | 3hr0 | Asp538-Chain A, Lys541-Chain A, Glu662-Chain A |
| 3 | 3lrk | Asp35-Chain A, Asp72-Chain A, Lys147-Chain A |
| | 5coz | Glu306-Chain A, Lys309-Chain A, Asp368-Chain A |
| | 2ygk | Glu77-Chain A, Lys304-Chain A, Asp308-Chain A |
| | 4whn | Glu9-Chain C, Glu13-Chain C, Lys49-Chain C |
| | 3sd7 | Asp11-Chain A, Lys144-Chain A, Asp172-Chain A |
| | 4yn8 | Asp8-Chain A, Asp50-Chain A, Lys99-Chain A |
| | 4i90 | Glu207-Chain A, Asp210-Chain A, Lys214-Chain A |
| | 6kqs | Asp100-Chain A, Lys453-Chain A, Glu497-Chain A |
| | 4ri2 | Lys34-Chain A, Glu37-Chain A, Asp98-Chain A |

| | | |
|---|---|---|
| 4 | 4zr8 | Glu92-Chain B, Glu95-Chain B, Lys98-Chain B |
| | 5fc2 | Glu1773-Chain B, Lys1776-Chain B, Glu1777-Chain B |
| | 4hye | Glu141-Chain A, Glu143-Chain B, Lys180-Chain B |
| | 3e6c | Lys62-Chain C, Asp64-Chain C, Glu72-Chain C |
| | 1c3c | Lys387-Chain A, Glu391-Chain A, Glu395-Chain A |
| | 4qdc | Lys166-Chain A, Glu298-Chain A, Glu302-Chain A |
| | 3mgg | Lys224-Chain A, Glu235-Chain A, Glu239-Chain A |
| 5 | 3nua | Glu91-Chain A, Lys178-Chain A, Glu180-Chain A |
| | 3sdb | Asp372-Chain A, Asp482-Chain A, Lys510-Chain A |
| | 4q6j | Glu36-Chain A, Glu122-Chain A, Lys173-Chain A |
| | 5gt5 | Lys271-Chain A, Asp272-Chain A, Asp296-Chain A |
| | 4wep | Asp92-Chain A, Asp95-Chain A, Lys107-Chain A |
| | 3q1n | Lys216-Chain A, Asp231-Chain A, Glu291-Chain A |
| 6 | 2zfn | Lys105-Chain A, Glu109-Chain A, Glu232-Chain A |
| | 4z39 | Glu21-Chain A, Lys28-Chain A, Asp116-Chain B |
| | 3i42 | Glu25-Chain A, Asp69-Chain A, Lys120-Chain A |
| | 5ytq | Lys69-Chain A, Glu72-Chain A, Glu39-Chain B |
| | 2qvg | Glu14-Chain A, Asp66-Chain A, Lys118-Chain A |
| | 5x5j | Glu19-Chain A, Asp63-Chain A, Lys112-Chain A |
| 7 | 4hzi | Glu236-Chain A, Glu240-Chain A, Lys261-Chain A |
| | 4i6k | Glu132-Chain A, Asp157-Chain A, Lys188-Chain A |
| | 5jaz | Lys205-Chain A, Asp231-Chain A, Glu318-Chain A |
| | 5i6r | Lys6-Chain A, Glu12-Chain A, Glu16-Chain A |
| | 2xxp | Lys172-Chain A, Glu195-Chain A, Asp454-Chain A |
| | 3o3o | Asp82-Chain A, Lys347-Chain A, Glu329-Chain B |
| 8 | 2yk4 | Glu159-Chain A, Lys161-Chain A, Asp209-Chain A |
| | 6w40 | Glu3-Chain A, Asp73-Chain B, Lys75-Chain B |
| | 2rc3 | Lys24-Chain A, Glu32-Chain A, Asp146-Chain A |
| | 6dyf | Glu18-Chain A, Glu92-Chain A, Lys95-Chain A |
| | 3mvp | Lys151-Chain A, Asp179-Chain A, Glu187-Chain B |
| | 5cdv | Glu246-Chain A, Asp329-Chain A, Lys346-Chain A |
