## Supplementary material for "Elucidation of the molecular interactions that enable stable assembly and structural diversity in multicomponent immune receptors": SI DFT Coordinates

**Table S4.** Relaxed coordinates of Complex 2

| | | | |
|---|---|---|---|
| C | 35.07600 | 29.61800 | 50.32400 |
| O | 35.29000 | 29.74100 | 49.12000 |
| N | 33.82600 | 29.84000 | 50.82400 |
| C | 32.80700 | 30.46300 | 50.04700 |
| C | 33.11400 | 31.92700 | 49.53200 |
| O | 32.91300 | 32.28500 | 48.36700 |
| C | 31.46320 | 30.42890 | 50.80920 |
| C | 30.29080 | 30.73800 | 49.89190 |
| O | 29.82090 | 31.87440 | 49.79440 |
| O | 29.87110 | 29.67260 | 49.23530 |
| H | 33.70520 | 29.79390 | 51.83460 |
| H | 32.69630 | 29.90030 | 49.10730 |
| H | 31.33160 | 29.41690 | 51.21570 |
| H | 31.46590 | 31.15580 | 51.63260 |
| H | 29.05410 | 29.84880 | 48.62640 |
| N | 33.63800 | 32.82200 | 50.35700 |
| H | 33.88250 | 32.60970 | 51.31960 |
| C | 33.90300 | 32.02700 | 44.81800 |
| O | 33.71700 | 32.08700 | 43.59100 |
| N | 33.39200 | 32.93300 | 45.68900 |
| C | 32.51000 | 34.06700 | 45.26000 |
| C | 33.32800 | 35.05900 | 44.40200 |
| O | 32.77200 | 35.51900 | 43.37500 |
| C | 31.81190 | 34.70740 | 46.47970 |
| C | 30.99590 | 35.92790 | 46.08200 |
| O | 30.90170 | 33.77310 | 47.09090 |
| H | 33.48010 | 32.73540 | 46.68870 |
| H | 31.74040 | 33.67540 | 44.58110 |
| H | 32.58680 | 34.99730 | 47.21560 |
| H | 31.43560 | 33.11480 | 47.58960 |
| H | 30.49690 | 36.33610 | 46.97120 |
| H | 30.23390 | 35.64980 | 45.34060 |
| H | 31.63030 | 36.70810 | 45.64280 |
| N | 34.61300 | 35.31200 | 44.78500 |
| H | 35.05150 | 34.70490 | 45.47020 |
| C | 23.94400 | 27.94100 | 45.50200 |
| O | 23.57900 | 28.09500 | 44.33100 |
| N | 25.20400 | 28.15800 | 45.87400 |
| C | 26.22300 | 28.52000 | 44.89200 |
| C | 26.47000 | 27.38700 | 43.85800 |
| O | 26.67800 | 27.71100 | 42.68900 |
| C | 27.57100 | 28.80890 | 45.58060 |
| C | 27.57550 | 30.01170 | 46.52430 |
| O | 27.36760 | 31.16380 | 46.01180 |
| O | 27.79320 | 29.82030 | 47.75570 |
| H | 25.46860 | 28.05610 | 46.85190 |
| H | 25.88590 | 29.38280 | 44.30400 |
| H | 27.90390 | 27.92000 | 46.13420 |
| H | 28.30260 | 29.01290 | 44.78440 |
| N | 26.43600 | 26.10900 | 44.22600 |
| H | 26.15480 | 25.82150 | 45.15760 |
| C | 22.02800 | 35.96100 | 51.62100 |
| O | 21.52500 | 35.67200 | 52.73400 |
| N | 23.04900 | 35.23800 | 51.14800 |
| C | 23.74700 | 34.28100 | 52.02300 |
| C | 24.10700 | 34.76800 | 53.38500 |
| O | 23.80400 | 34.05500 | 54.34000 |
| C | 25.02640 | 33.78580 | 51.29570 |
| C | 24.77450 | 33.00110 | 49.99780 |
| C | 26.05670 | 32.48790 | 49.31350 |
| C | 26.84850 | 33.58150 | 48.59250 |
| N | 28.05980 | 32.99730 | 47.92690 |
| H | 23.49660 | 35.52290 | 50.27890 |
| H | 23.07610 | 33.42890 | 52.20090 |
| H | 25.67380 | 34.65930 | 51.10480 |
| H | 25.57570 | 33.14380 | 52.00200 |
| H | 24.13650 | 32.13560 | 50.23490 |
| H | 24.20340 | 33.60590 | 49.27420 |
| H | 26.70450 | 31.99230 | 50.05580 |
| H | 25.78610 | 31.72060 | 48.57350 |
| H | 26.23390 | 34.03610 | 47.80390 |
| H | 27.19480 | 34.36340 | 49.28180 |
| H | 28.65540 | 32.48020 | 48.61770 |
| H | 28.67630 | 33.70380 | 47.49750 |
| H | 27.77660 | 32.29910 | 47.16970 |
| N | 24.74100 | 35.92700 | 53.53200 |
| H | 24.85570 | 36.57440 | 52.75890 |
| O | 25.02000 | 32.63400 | 45.94200 |
| H | 25.77730 | 32.00510 | 45.92680 |
| H | 25.01390 | 32.99680 | 45.03920 |
| O | 27.76100 | 32.56500 | 43.56800 |
| H | 27.92580 | 31.98250 | 42.80770 |
| H | 27.68420 | 31.95400 | 44.33640 |
| H | 35.81620 | 29.27140 | 51.07360 |
| H | 33.84670 | 33.74950 | 50.00060 |
| H | 34.49820 | 31.23070 | 45.31360 |
| H | 35.20960 | 35.82390 | 44.14210 |
| H | 23.26620 | 27.63570 | 46.32800 |
| H | 26.53700 | 25.39970 | 43.50750 |

| | | | |
|---|---|---|---|
| H | 21.65580 | 36.76440 | 50.95380 |
| H | 24.88670 | 36.28040 | 54.47230 |

**Table S5.** Step 19 of the relaxed surface scan for proton transfer from complex 2 to complex 3 with explicit waters.

| | | | |
|---|---|---|---|
| C | 35.07600 | 29.61800 | 50.32400 |
| O | 35.29000 | 29.74100 | 49.12000 |
| N | 33.82600 | 29.84000 | 50.82400 |
| C | 32.80700 | 30.46300 | 50.04700 |
| C | 33.11400 | 31.92700 | 49.53200 |
| O | 32.91300 | 32.28500 | 48.36700 |
| C | 31.48110 | 30.49190 | 50.83310 |
| C | 30.30680 | 30.93800 | 49.95250 |
| O | 29.79320 | 32.06940 | 50.14620 |
| O | 29.96270 | 30.08600 | 49.06470 |
| H | 33.69840 | 29.78390 | 51.83300 |
| H | 32.65940 | 29.89680 | 49.11590 |
| H | 31.27400 | 29.47170 | 51.18750 |
| H | 31.56070 | 31.16550 | 51.69860 |
| H | 28.81210 | 29.94560 | 48.38000 |
| N | 33.63800 | 32.82200 | 50.35700 |
| H | 33.85820 | 32.61530 | 51.32650 |
| C | 33.90300 | 32.02700 | 44.81800 |
| O | 33.71700 | 32.08700 | 43.59100 |
| N | 33.39200 | 32.93300 | 45.68900 |
| C | 32.51000 | 34.06700 | 45.26000 |
| C | 33.32800 | 35.05900 | 44.40200 |
| O | 32.77200 | 35.51900 | 43.37500 |
| C | 31.84400 | 34.72470 | 46.49240 |
| C | 31.00480 | 35.93290 | 46.10410 |
| O | 30.96800 | 33.80130 | 47.15670 |
| H | 33.46900 | 32.72700 | 46.68870 |
| H | 31.72900 | 33.67980 | 44.59220 |
| H | 32.64710 | 35.03730 | 47.18850 |
| H | 31.52580 | 33.16250 | 47.65870 |
| H | 30.56480 | 36.37300 | 47.00880 |
| H | 30.19040 | 35.62350 | 45.43480 |
| H | 31.60520 | 36.69590 | 45.59320 |
| N | 34.61300 | 35.31200 | 44.78500 |
| H | 35.05070 | 34.71040 | 45.47550 |
| C | 23.94400 | 27.94100 | 45.50200 |
| O | 23.57900 | 28.09500 | 44.33100 |
| N | 25.20400 | 28.15800 | 45.87400 |
| C | 26.22300 | 28.52000 | 44.89200 |
| C | 26.47000 | 27.38700 | 43.85800 |
| O | 26.67800 | 27.71100 | 42.68900 |
| C | 27.59240 | 28.71570 | 45.56520 |
| C | 27.74890 | 29.82520 | 46.59030 |
| O | 27.64440 | 31.02480 | 46.26760 |
| O | 28.05290 | 29.39060 | 47.78040 |
| H | 25.47340 | 28.02160 | 46.84630 |
| H | 25.90830 | 29.39820 | 44.31670 |
| H | 27.90090 | 27.77130 | 46.03260 |
| H | 28.31110 | 28.94220 | 44.76240 |
| N | 26.43600 | 26.10900 | 44.22600 |
| H | 26.15180 | 25.81730 | 45.15530 |
| C | 22.02800 | 35.96100 | 51.62100 |
| O | 21.52500 | 35.67200 | 52.73400 |
| N | 23.04900 | 35.23800 | 51.14800 |
| C | 23.74700 | 34.28100 | 52.02300 |
| C | 24.10700 | 34.76800 | 53.38500 |
| O | 23.80400 | 34.05500 | 54.34000 |
| C | 25.04120 | 33.80230 | 51.31440 |
| C | 24.83180 | 33.04100 | 49.99810 |
| C | 26.14120 | 32.53180 | 49.36810 |
| C | 27.01800 | 33.63990 | 48.77940 |
| N | 28.24140 | 33.03720 | 48.16140 |
| H | 23.49690 | 35.52320 | 50.27920 |
| H | 23.08110 | 33.42440 | 52.19660 |
| H | 25.68760 | 34.68210 | 51.15340 |
| H | 25.57830 | 33.15220 | 52.02280 |
| H | 24.17650 | 32.17750 | 50.19220 |
| H | 24.30240 | 33.66550 | 49.25940 |
| H | 26.71890 | 31.96020 | 50.11380 |
| H | 25.89290 | 31.83060 | 48.55520 |
| H | 26.46780 | 34.17550 | 47.99520 |
| H | 27.35600 | 34.35010 | 49.54550 |
| H | 28.84520 | 32.59090 | 48.92810 |
| H | 28.84050 | 33.70900 | 47.65430 |
| H | 27.97780 | 32.29190 | 47.47130 |
| N | 24.74100 | 35.92700 | 53.53200 |
| H | 24.86110 | 36.57330 | 52.75890 |
| O | 25.64500 | 33.31530 | 45.88490 |
| H | 25.71890 | 32.35200 | 46.00520 |
| H | 26.51920 | 33.54550 | 45.48800 |
| O | 28.32850 | 33.50680 | 45.08060 |
| H | 28.72650 | 33.79930 | 44.24300 |
| H | 28.40160 | 32.53060 | 45.07660 |
| H | 35.81760 | 29.27250 | 51.07320 |
| H | 33.82540 | 33.75690 | 50.00850 |

| | | | |
|---|---|---|---|
| H | 34.49740 | 31.23040 | 45.31340 |
| H | 35.20880 | 35.83150 | 44.14760 |
| H | 23.26740 | 27.63110 | 46.32730 |
| H | 26.54560 | 25.40130 | 43.50720 |
| H | 21.65630 | 36.76480 | 50.95400 |
| H | 24.89350 | 36.27710 | 54.47240 |

**Table S6.** Step 18 of the relaxed surface scan for proton transfer from complex 2 to complex 3 with explicit waters.

| | | | |
|---|---|---|---|
| C | 35.07600 | 29.61800 | 50.32400 |
| O | 35.29000 | 29.74100 | 49.12000 |
| N | 33.82600 | 29.84000 | 50.82400 |
| C | 32.80700 | 30.46300 | 50.04700 |
| C | 33.11400 | 31.92700 | 49.53200 |
| O | 32.91300 | 32.28500 | 48.36700 |
| C | 31.43520 | 30.40690 | 50.74560 |
| C | 30.29540 | 30.74800 | 49.77500 |
| O | 29.80860 | 31.89880 | 49.79120 |
| O | 29.94030 | 29.77580 | 49.00140 |
| H | 33.70460 | 29.79340 | 51.83460 |
| H | 32.72360 | 29.90670 | 49.09980 |
| H | 31.28630 | 29.38050 | 51.10910 |
| H | 31.39980 | 31.10290 | 51.59550 |
| H | 28.92300 | 29.89540 | 48.31760 |
| N | 33.63800 | 32.82200 | 50.35700 |
| H | 33.86950 | 32.61100 | 51.32300 |
| C | 33.90300 | 32.02700 | 44.81800 |
| O | 33.71700 | 32.08700 | 43.59100 |
| N | 33.39200 | 32.93300 | 45.68900 |
| C | 32.51000 | 34.06700 | 45.26000 |
| C | 33.32800 | 35.05900 | 44.40200 |
| O | 32.77200 | 35.51900 | 43.37500 |
| C | 31.82090 | 34.70930 | 46.48620 |
| C | 30.98410 | 35.91740 | 46.09280 |
| O | 30.93180 | 33.77440 | 47.11880 |
| H | 33.47350 | 32.73190 | 46.68920 |
| H | 31.73700 | 33.67660 | 44.58440 |
| H | 32.60690 | 35.01530 | 47.20390 |
| H | 31.47580 | 33.13120 | 47.62840 |
| H | 30.52250 | 36.34680 | 46.99190 |
| H | 30.18640 | 35.61030 | 45.40210 |
| H | 31.59140 | 36.68800 | 45.60170 |
| N | 34.61300 | 35.31200 | 44.78500 |
| H | 35.04980 | 34.71140 | 45.47700 |
| C | 23.94400 | 27.94100 | 45.50200 |
| O | 23.57900 | 28.09500 | 44.33100 |
| N | 25.20400 | 28.15800 | 45.87400 |
| C | 26.22300 | 28.52000 | 44.89200 |
| C | 26.47000 | 27.38700 | 43.85800 |
| O | 26.67800 | 27.71100 | 42.68900 |
| C | 27.56460 | 28.80540 | 45.59950 |
| C | 27.57730 | 30.02800 | 46.51570 |
| O | 27.24790 | 31.14440 | 46.05090 |
| O | 27.91230 | 29.84820 | 47.75260 |
| H | 25.47190 | 28.02860 | 46.84750 |
| H | 25.89100 | 29.38560 | 44.30680 |
| H | 27.88490 | 27.92190 | 46.16860 |
| H | 28.31430 | 28.99230 | 44.81490 |
| N | 26.43600 | 26.10900 | 44.22600 |
| H | 26.15080 | 25.82030 | 45.15600 |
| C | 22.02800 | 35.96100 | 51.62100 |
| O | 21.52500 | 35.67200 | 52.73400 |
| N | 23.04900 | 35.23800 | 51.14800 |
| C | 23.74700 | 34.28100 | 52.02300 |
| C | 24.10700 | 34.76800 | 53.38500 |
| O | 23.80400 | 34.05500 | 54.34000 |
| C | 25.03170 | 33.79360 | 51.30450 |
| C | 24.79210 | 33.03470 | 49.99190 |
| C | 26.07940 | 32.52040 | 49.32460 |
| C | 26.95220 | 33.62800 | 48.72910 |
| N | 28.10150 | 33.01930 | 47.98710 |
| H | 23.49680 | 35.52370 | 50.27920 |
| H | 23.07800 | 33.42700 | 52.19900 |
| H | 25.68300 | 34.66820 | 51.13400 |
| H | 25.57150 | 33.13900 | 52.00640 |
| H | 24.13770 | 32.17380 | 50.20050 |
| H | 24.24700 | 33.66230 | 49.26750 |
| H | 26.67030 | 31.92980 | 50.04410 |
| H | 25.80300 | 31.83270 | 48.51000 |
| H | 26.37140 | 34.22090 | 48.01080 |
| H | 27.37000 | 34.28820 | 49.50050 |
| H | 28.73270 | 32.49130 | 48.65870 |
| H | 28.69290 | 33.70290 | 47.48950 |
| H | 27.74620 | 32.33610 | 47.27010 |
| N | 24.74100 | 35.92700 | 53.53200 |
| H | 24.85860 | 36.57370 | 52.75880 |
| O | 25.37060 | 33.54270 | 45.88800 |
| H | 25.45930 | 32.57440 | 45.95450 |
| H | 26.21750 | 33.79800 | 45.45490 |
| O | 28.06560 | 33.62130 | 44.98720 |

| | | | |
|---|---|---|---|
| H | 28.49360 | 33.86720 | 44.14970 |
| H | 28.00470 | 32.64260 | 44.97240 |
| H | 35.81620 | 29.27030 | 51.07330 |
| H | 33.84230 | 33.75180 | 50.00420 |
| H | 34.49770 | 31.23050 | 45.31340 |
| H | 35.20960 | 35.82920 | 44.14660 |
| H | 23.26670 | 27.63250 | 46.32740 |
| H | 26.53460 | 25.40030 | 43.50640 |
| H | 21.65610 | 36.76450 | 50.95380 |
| H | 24.89020 | 36.27880 | 54.47230 |

**Table S7.** Step 17 of the relaxed surface scan for proton transfer from complex 2 to complex 3 with explicit waters.

| | | | |
|---|---|---|---|
| C | 35.07600 | 29.61800 | 50.32400 |
| O | 35.29000 | 29.74100 | 49.12000 |
| N | 33.82600 | 29.84000 | 50.82400 |
| C | 32.80700 | 30.46300 | 50.04700 |
| C | 33.11400 | 31.92700 | 49.53200 |
| O | 32.91300 | 32.28500 | 48.36700 |
| C | 31.46960 | 30.49900 | 50.81100 |
| C | 30.30440 | 30.96330 | 49.92280 |
| O | 29.61670 | 31.95340 | 50.28400 |
| O | 30.12730 | 30.27520 | 48.85950 |
| H | 33.70240 | 29.79140 | 51.83400 |
| H | 32.66470 | 29.89970 | 49.11390 |
| H | 31.23780 | 29.47610 | 51.14620 |
| H | 31.53940 | 31.15220 | 51.69210 |
| H | 28.93090 | 30.04270 | 48.25510 |
| N | 33.63800 | 32.82200 | 50.35700 |
| H | 33.86770 | 32.61190 | 51.32360 |
| C | 33.90300 | 32.02700 | 44.81800 |
| O | 33.71700 | 32.08700 | 43.59100 |
| N | 33.39200 | 32.93300 | 45.68900 |
| C | 32.51000 | 34.06700 | 45.26000 |
| C | 33.32800 | 35.05900 | 44.40200 |
| O | 32.77200 | 35.51900 | 43.37500 |
| C | 31.81410 | 34.70360 | 46.48360 |
| C | 30.97760 | 35.91070 | 46.08910 |
| O | 30.92500 | 33.75530 | 47.10620 |
| H | 33.47710 | 32.73250 | 46.68900 |
| H | 31.73980 | 33.67570 | 44.58160 |
| H | 32.59130 | 35.00600 | 47.21140 |
| H | 31.47780 | 33.10690 | 47.60190 |
| H | 30.48710 | 36.32110 | 46.98200 |
| H | 30.20780 | 35.61780 | 45.36150 |
| H | 31.59670 | 36.69410 | 45.63410 |
| N | 34.61300 | 35.31200 | 44.78500 |
| H | 35.05080 | 34.70760 | 45.47310 |
| C | 23.94400 | 27.94100 | 45.50200 |
| O | 23.57900 | 28.09500 | 44.33100 |
| N | 25.20400 | 28.15800 | 45.87400 |
| C | 26.22300 | 28.52000 | 44.89200 |
| C | 26.47000 | 27.38700 | 43.85800 |
| O | 26.67800 | 27.71100 | 42.68900 |
| C | 27.59000 | 28.74190 | 45.55500 |
| C | 27.75310 | 29.87220 | 46.55350 |
| O | 27.56910 | 31.06810 | 46.23960 |
| O | 28.16550 | 29.46570 | 47.72160 |
| H | 25.47050 | 28.02560 | 46.84740 |
| H | 25.89660 | 29.39110 | 44.31100 |
| H | 27.91670 | 27.81210 | 46.03880 |
| H | 28.30510 | 28.96050 | 44.74560 |
| N | 26.43600 | 26.10900 | 44.22600 |
| H | 26.15430 | 25.81770 | 45.15630 |
| C | 22.02800 | 35.96100 | 51.62100 |
| O | 21.52500 | 35.67200 | 52.73400 |
| N | 23.04900 | 35.23800 | 51.14800 |
| C | 23.74700 | 34.28100 | 52.02300 |
| C | 24.10700 | 34.76800 | 53.38500 |
| O | 23.80400 | 34.05500 | 54.34000 |
| C | 25.03350 | 33.79260 | 51.30630 |
| C | 24.80620 | 33.03170 | 49.99110 |
| C | 26.10590 | 32.50380 | 49.35410 |
| C | 26.96380 | 33.59310 | 48.70400 |
| N | 28.24250 | 32.99880 | 48.19510 |
| H | 23.49700 | 35.52360 | 50.27940 |
| H | 23.07800 | 33.42700 | 52.19870 |
| H | 25.68710 | 34.66650 | 51.14180 |
| H | 25.56850 | 33.13710 | 52.01100 |
| H | 24.14420 | 32.17530 | 50.19240 |
| H | 24.27540 | 33.65770 | 49.25510 |
| H | 26.70410 | 31.97640 | 50.11570 |
| H | 25.84860 | 31.76390 | 48.58040 |
| H | 26.43330 | 34.04330 | 47.85350 |
| H | 27.23370 | 34.38250 | 49.41770 |
| H | 28.77520 | 32.53530 | 49.00640 |
| H | 28.89220 | 33.68060 | 47.76800 |
| H | 28.03870 | 32.26990 | 47.46740 |
| N | 24.74100 | 35.92700 | 53.53200 |
| H | 24.86020 | 36.57340 | 52.75890 |

| | | | |
|---|---|---|---|
| O | 25.02000 | 32.63400 | 45.94200 |
| H | 25.77700 | 32.01550 | 45.99180 |
| H | 25.08730 | 32.99230 | 45.03900 |
| O | 27.76100 | 32.56500 | 43.56800 |
| H | 28.12660 | 32.02800 | 42.84460 |
| H | 27.81400 | 31.98460 | 44.35470 |
| H | 35.81760 | 29.27260 | 51.07330 |
| H | 33.83570 | 33.75390 | 50.00600 |
| H | 34.49750 | 31.23060 | 45.31370 |
| H | 35.20980 | 35.82520 | 44.14340 |
| H | 23.26750 | 27.63290 | 46.32770 |
| H | 26.54430 | 25.40100 | 43.50720 |
| H | 21.65600 | 36.76460 | 50.95390 |
| H | 24.89220 | 36.27780 | 54.47240 |

**Table S8.** Step 16 of the relaxed surface scan for proton transfer from complex 2 to complex 3 with explicit waters.

| | | | |
|---|---|---|---|
| C | 35.07600 | 29.61800 | 50.32400 |
| O | 35.29000 | 29.74100 | 49.12000 |
| N | 33.82600 | 29.84000 | 50.82400 |
| C | 32.80700 | 30.46300 | 50.04700 |
| C | 33.11400 | 31.92700 | 49.53200 |
| O | 32.91300 | 32.28500 | 48.36700 |
| C | 31.46680 | 30.49150 | 50.80610 |
| C | 30.29840 | 30.92780 | 49.91000 |
| O | 29.60230 | 31.91680 | 50.24780 |
| O | 30.12890 | 30.20950 | 48.86130 |
| H | 33.70330 | 29.79290 | 51.83420 |
| H | 32.66980 | 29.90000 | 49.11270 |
| H | 31.24740 | 29.46980 | 51.15310 |
| H | 31.52420 | 31.15640 | 51.67920 |
| H | 28.96880 | 30.03910 | 48.26920 |
| N | 33.63800 | 32.82200 | 50.35700 |
| H | 33.87240 | 32.61040 | 51.32220 |
| C | 33.90300 | 32.02700 | 44.81800 |
| O | 33.71700 | 32.08700 | 43.59100 |
| N | 33.39200 | 32.93300 | 45.68900 |
| C | 32.51000 | 34.06700 | 45.26000 |
| C | 33.32800 | 35.05900 | 44.40200 |
| O | 32.77200 | 35.51900 | 43.37500 |
| C | 31.81190 | 34.70310 | 46.48270 |
| C | 30.97480 | 35.90930 | 46.08680 |
| O | 30.92230 | 33.75420 | 47.10430 |
| H | 33.47690 | 32.73240 | 46.68890 |
| H | 31.74040 | 33.67540 | 44.58110 |
| H | 32.58780 | 35.00610 | 47.21180 |
| H | 31.47520 | 33.10690 | 47.60070 |
| H | 30.48330 | 36.31990 | 46.97900 |
| H | 30.20580 | 35.61500 | 45.35890 |
| H | 31.59360 | 36.69280 | 45.63160 |
| N | 34.61300 | 35.31200 | 44.78500 |
| H | 35.05080 | 34.70710 | 45.47270 |
| C | 23.94400 | 27.94100 | 45.50200 |
| O | 23.57900 | 28.09500 | 44.33100 |
| N | 25.20400 | 28.15800 | 45.87400 |
| C | 26.22300 | 28.52000 | 44.89200 |
| C | 26.47000 | 27.38700 | 43.85800 |
| O | 26.67800 | 27.71100 | 42.68900 |
| C | 27.58830 | 28.74780 | 45.55800 |
| C | 27.74250 | 29.88630 | 46.55170 |
| O | 27.55360 | 31.07900 | 46.21890 |
| O | 28.14100 | 29.49980 | 47.72700 |
| H | 25.47020 | 28.02800 | 46.84790 |
| H | 25.89530 | 29.39010 | 44.31000 |
| H | 27.91550 | 27.82180 | 46.04870 |
| H | 28.30520 | 28.96330 | 44.74940 |
| N | 26.43600 | 26.10900 | 44.22600 |
| H | 26.15590 | 25.81800 | 45.15680 |
| C | 22.02800 | 35.96100 | 51.62100 |
| O | 21.52500 | 35.67200 | 52.73400 |
| N | 23.04900 | 35.23800 | 51.14800 |
| C | 23.74700 | 34.28100 | 52.02300 |
| C | 24.10700 | 34.76800 | 53.38500 |
| O | 23.80400 | 34.05500 | 54.34000 |
| C | 25.03110 | 33.79050 | 51.30360 |
| C | 24.79750 | 33.02910 | 49.98940 |
| C | 26.09360 | 32.49980 | 49.34630 |
| C | 26.94310 | 33.58590 | 48.67950 |
| N | 28.22250 | 32.98960 | 48.17260 |
| H | 23.49640 | 35.52310 | 50.27890 |
| H | 23.07730 | 33.42770 | 52.19960 |
| H | 25.68600 | 34.66310 | 51.13660 |
| H | 25.56700 | 33.13420 | 52.00700 |
| H | 24.13570 | 32.17340 | 50.19450 |
| H | 24.26320 | 33.65480 | 49.25560 |
| H | 26.69930 | 31.98060 | 50.10770 |
| H | 25.83310 | 31.75190 | 48.58150 |
| H | 26.40690 | 34.02150 | 47.82510 |
| H | 27.21240 | 34.38530 | 49.38240 |
| H | 28.74570 | 32.51970 | 48.98310 |

| | | | |
|---|---|---|---|
| H | 28.87850 | 33.67020 | 47.75350 |
| H | 28.01720 | 32.26250 | 47.44060 |
| N | 24.74100 | 35.92700 | 53.53200 |
| H | 24.85840 | 36.57410 | 52.75920 |
| O | 25.02000 | 32.63400 | 45.94200 |
| H | 25.77420 | 32.01090 | 45.98190 |
| H | 25.08810 | 33.00510 | 45.04430 |
| O | 27.76100 | 32.56500 | 43.56800 |
| H | 28.12320 | 32.02230 | 42.84730 |
| H | 27.80380 | 31.98600 | 44.35680 |
| H | 35.81750 | 29.27250 | 51.07340 |
| H | 33.83980 | 33.75260 | 50.00480 |
| H | 34.49720 | 31.23030 | 45.31370 |
| H | 35.20990 | 35.82460 | 44.14300 |
| H | 23.26740 | 27.63300 | 46.32770 |
| H | 26.54450 | 25.40080 | 43.50750 |
| H | 21.65610 | 36.76460 | 50.95390 |
| H | 24.89070 | 36.27850 | 54.47240 |

**Table S9.** Step 15 of the relaxed surface scan for proton transfer from complex 2 to complex 3 with explicit waters.

| | | | |
|---|---|---|---|
| C | 35.07600 | 29.61800 | 50.32400 |
| O | 35.29000 | 29.74100 | 49.12000 |
| N | 33.82600 | 29.84000 | 50.82400 |
| C | 32.80700 | 30.46300 | 50.04700 |
| C | 33.11400 | 31.92700 | 49.53200 |
| O | 32.91300 | 32.28500 | 48.36700 |
| C | 31.46210 | 30.47530 | 50.79930 |
| C | 30.29120 | 30.87850 | 49.89270 |
| O | 29.63890 | 31.91580 | 50.15320 |
| O | 30.07760 | 30.07300 | 48.91230 |
| H | 33.70330 | 29.79230 | 51.83420 |
| H | 32.67870 | 29.89990 | 49.11100 |
| H | 31.26410 | 29.45390 | 51.15830 |
| H | 31.50210 | 31.15500 | 51.66170 |
| H | 28.96810 | 30.01010 | 48.31120 |
| N | 33.63800 | 32.82200 | 50.35700 |
| H | 33.87200 | 32.61070 | 51.32230 |
| C | 33.90300 | 32.02700 | 44.81800 |
| O | 33.71700 | 32.08700 | 43.59100 |
| N | 33.39200 | 32.93300 | 45.68900 |
| C | 32.51000 | 34.06700 | 45.26000 |
| C | 33.32800 | 35.05900 | 44.40200 |
| O | 32.77200 | 35.51900 | 43.37500 |
| C | 31.81330 | 34.70560 | 46.48170 |
| C | 30.97970 | 35.91400 | 46.08470 |
| O | 30.92070 | 33.76000 | 47.10360 |
| H | 33.47820 | 32.73280 | 46.68870 |
| H | 31.74010 | 33.67550 | 44.58150 |
| H | 32.58960 | 35.00740 | 47.21090 |
| H | 31.47160 | 33.11410 | 47.60340 |
| H | 30.48700 | 36.32470 | 46.97620 |
| H | 30.21190 | 35.62220 | 45.35460 |
| H | 31.60110 | 36.69680 | 45.63180 |
| N | 34.61300 | 35.31200 | 44.78500 |
| H | 35.05120 | 34.70660 | 45.47200 |
| C | 23.94400 | 27.94100 | 45.50200 |
| O | 23.57900 | 28.09500 | 44.33100 |
| N | 25.20400 | 28.15800 | 45.87400 |
| C | 26.22300 | 28.52000 | 44.89200 |
| C | 26.47000 | 27.38700 | 43.85800 |
| O | 26.67800 | 27.71100 | 42.68900 |
| C | 27.58520 | 28.76290 | 45.56260 |
| C | 27.71370 | 29.91460 | 46.54870 |
| O | 27.53310 | 31.10140 | 46.17880 |
| O | 28.06380 | 29.56320 | 47.74450 |
| H | 25.47050 | 28.03120 | 46.84820 |
| H | 25.89250 | 29.38810 | 44.30840 |
| H | 27.91550 | 27.84430 | 46.06520 |
| H | 28.30330 | 28.97780 | 44.75530 |
| N | 26.43600 | 26.10900 | 44.22600 |
| H | 26.15630 | 25.81870 | 45.15720 |
| C | 22.02800 | 35.96100 | 51.62100 |
| O | 21.52500 | 35.67200 | 52.73400 |
| N | 23.04900 | 35.23800 | 51.14800 |
| C | 23.74700 | 34.28100 | 52.02300 |
| C | 24.10700 | 34.76800 | 53.38500 |
| O | 23.80400 | 34.05500 | 54.34000 |
| C | 25.03030 | 33.78960 | 51.30240 |
| C | 24.79330 | 33.02270 | 49.99190 |
| C | 26.08720 | 32.50020 | 49.33890 |
| C | 26.92370 | 33.59120 | 48.66480 |
| N | 28.19180 | 33.00100 | 48.12380 |
| H | 23.49680 | 35.52340 | 50.27920 |
| H | 23.07700 | 33.42800 | 52.19980 |
| H | 25.68320 | 34.66260 | 51.13010 |
| H | 25.56930 | 33.13670 | 52.00660 |
| H | 24.13860 | 32.16340 | 50.20440 |
| H | 24.24930 | 33.64310 | 49.26070 |
| H | 26.70220 | 31.98360 | 50.09460 |

| | | | |
|---|---|---|---|
| H | 25.82480 | 31.75090 | 48.57630 |
| H | 26.37080 | 34.03330 | 47.82460 |
| H | 27.20670 | 34.38490 | 49.36890 |
| H | 28.73390 | 32.52230 | 48.91110 |
| H | 28.83720 | 33.68960 | 47.70320 |
| H | 27.97300 | 32.27970 | 47.38640 |
| N | 24.74100 | 35.92700 | 53.53200 |
| H | 24.85900 | 36.57370 | 52.75890 |
| O | 25.02000 | 32.63400 | 45.94200 |
| H | 25.77800 | 32.01410 | 45.97130 |
| H | 25.07480 | 33.00680 | 45.04420 |
| O | 27.76100 | 32.56500 | 43.56800 |
| H | 28.06720 | 32.00620 | 42.83380 |
| H | 27.78470 | 31.97620 | 44.35150 |
| H | 35.81700 | 29.27160 | 51.07330 |
| H | 33.84040 | 33.75240 | 50.00460 |
| H | 34.49760 | 31.23060 | 45.31370 |
| H | 35.20970 | 35.82480 | 44.14300 |
| H | 23.26720 | 27.63330 | 46.32770 |
| H | 26.54280 | 25.40060 | 43.50750 |
| H | 21.65590 | 36.76450 | 50.95380 |
| H | 24.89090 | 36.27850 | 54.47230 |

**Table S10.** Step 14 of the relaxed surface scan for proton transfer from complex 2 to complex 3 with explicit waters.

| | | | |
|---|---|---|---|
| C | 35.07600 | 29.61800 | 50.32400 |
| O | 35.29000 | 29.74100 | 49.12000 |
| N | 33.82600 | 29.84000 | 50.82400 |
| C | 32.80700 | 30.46300 | 50.04700 |
| C | 33.11400 | 31.92700 | 49.53200 |
| O | 32.91300 | 32.28500 | 48.36700 |
| C | 31.43500 | 30.40620 | 50.74630 |
| C | 30.29530 | 30.74690 | 49.77710 |
| O | 29.80500 | 31.89330 | 49.78970 |
| O | 29.94260 | 29.76860 | 49.00310 |
| H | 33.70530 | 29.79490 | 51.83480 |
| H | 32.72430 | 29.90660 | 49.09980 |
| H | 31.28630 | 29.37970 | 51.10980 |
| H | 31.39930 | 31.10170 | 51.59620 |
| H | 28.96730 | 29.89610 | 48.33870 |
| N | 33.63800 | 32.82200 | 50.35700 |
| H | 33.87070 | 32.61070 | 51.32270 |
| C | 33.90300 | 32.02700 | 44.81800 |
| O | 33.71700 | 32.08700 | 43.59100 |
| N | 33.39200 | 32.93300 | 45.68900 |
| C | 32.51000 | 34.06700 | 45.26000 |
| C | 33.32800 | 35.05900 | 44.40200 |
| O | 32.77200 | 35.51900 | 43.37500 |
| C | 31.81980 | 34.70890 | 46.48570 |
| C | 30.98300 | 35.91670 | 46.09150 |
| O | 30.93030 | 33.77330 | 47.11680 |
| H | 33.47360 | 32.73210 | 46.68910 |
| H | 31.73730 | 33.67650 | 44.58410 |
| H | 32.60490 | 35.01480 | 47.20440 |
| H | 31.47420 | 33.13030 | 47.62670 |
| H | 30.51880 | 36.34480 | 46.98990 |
| H | 30.18750 | 35.61010 | 45.39810 |
| H | 31.59120 | 36.68830 | 45.60310 |
| N | 34.61300 | 35.31200 | 44.78500 |
| H | 35.04980 | 34.71140 | 45.47690 |
| C | 23.94400 | 27.94100 | 45.50200 |
| O | 23.57900 | 28.09500 | 44.33100 |
| N | 25.20400 | 28.15800 | 45.87400 |
| C | 26.22300 | 28.52000 | 44.89200 |
| C | 26.47000 | 27.38700 | 43.85800 |
| O | 26.67800 | 27.71100 | 42.68900 |
| C | 27.56440 | 28.80760 | 45.59940 |
| C | 27.57600 | 30.02970 | 46.51910 |
| O | 27.24000 | 31.14560 | 46.05080 |
| O | 27.91900 | 29.85180 | 47.75100 |
| H | 25.47220 | 28.02910 | 46.84750 |
| H | 25.88990 | 29.38510 | 44.30660 |
| H | 27.88730 | 27.92370 | 46.16670 |
| H | 28.31310 | 28.99660 | 44.81440 |
| N | 26.43600 | 26.10900 | 44.22600 |
| H | 26.15080 | 25.82050 | 45.15610 |
| C | 22.02800 | 35.96100 | 51.62100 |
| O | 21.52500 | 35.67200 | 52.73400 |
| N | 23.04900 | 35.23800 | 51.14800 |
| C | 23.74700 | 34.28100 | 52.02300 |
| C | 24.10700 | 34.76800 | 53.38500 |
| O | 23.80400 | 34.05500 | 54.34000 |
| C | 25.03080 | 33.79250 | 51.30360 |
| C | 24.78900 | 33.03200 | 49.99240 |
| C | 26.07470 | 32.51720 | 49.32260 |
| C | 26.94510 | 33.62420 | 48.72300 |
| N | 28.09270 | 33.01510 | 47.97800 |
| H | 23.49710 | 35.52420 | 50.27950 |
| H | 23.07760 | 33.42740 | 52.19930 |
| H | 25.68230 | 34.66660 | 51.13150 |

| | | | |
|---|---|---|---|
| H | 25.57110 | 33.13840 | 52.00570 |
| H | 24.13520 | 32.17110 | 50.20340 |
| H | 24.24200 | 33.65850 | 49.26840 |
| H | 26.66770 | 31.92810 | 50.04160 |
| H | 25.79670 | 31.82800 | 48.51000 |
| H | 26.36170 | 34.21520 | 48.00510 |
| H | 27.36510 | 34.28610 | 49.49190 |
| H | 28.72130 | 32.48690 | 48.64820 |
| H | 28.68420 | 33.69990 | 47.48240 |
| H | 27.73600 | 32.33140 | 47.26010 |
| N | 24.74100 | 35.92700 | 53.53200 |
| H | 24.85830 | 36.57380 | 52.75880 |
| O | 25.35790 | 33.53290 | 45.89090 |
| H | 25.45070 | 32.56480 | 45.95850 |
| H | 26.20270 | 33.78960 | 45.45460 |
| O | 28.04890 | 33.61210 | 44.98370 |
| H | 28.47590 | 33.85590 | 44.14500 |
| H | 27.98750 | 32.63290 | 44.97210 |
| H | 35.81620 | 29.27040 | 51.07340 |
| H | 33.84350 | 33.75140 | 50.00380 |
| H | 34.49770 | 31.23050 | 45.31330 |
| H | 35.20970 | 35.82910 | 44.14650 |
| H | 23.26650 | 27.63290 | 46.32740 |
| H | 26.53480 | 25.40020 | 43.50660 |
| H | 21.65590 | 36.76430 | 50.95370 |
| H | 24.88990 | 36.27900 | 54.47230 |

**Table S11.** Step 13 of the relaxed surface scan for proton transfer from complex 2 to complex 3 with explicit waters.

| | | | |
|---|---|---|---|
| C | 35.07600 | 29.61800 | 50.32400 |
| O | 35.29000 | 29.74100 | 49.12000 |
| N | 33.82600 | 29.84000 | 50.82400 |
| C | 32.80700 | 30.46300 | 50.04700 |
| C | 33.11400 | 31.92700 | 49.53200 |
| O | 32.91300 | 32.28500 | 48.36700 |
| C | 31.45540 | 30.45790 | 50.78860 |
| C | 30.28990 | 30.83670 | 49.86620 |
| O | 29.69890 | 31.92560 | 50.03000 |
| O | 30.01920 | 29.94780 | 48.96960 |
| H | 33.70420 | 29.79370 | 51.83440 |
| H | 32.69090 | 29.90080 | 49.10830 |
| H | 31.27470 | 29.43670 | 51.15510 |
| H | 31.47630 | 31.14860 | 51.64280 |
| H | 28.97500 | 29.97390 | 48.35100 |
| N | 33.63800 | 32.82200 | 50.35700 |
| H | 33.87300 | 32.61070 | 51.32210 |
| C | 33.90300 | 32.02700 | 44.81800 |
| O | 33.71700 | 32.08700 | 43.59100 |
| N | 33.39200 | 32.93300 | 45.68900 |
| C | 32.51000 | 34.06700 | 45.26000 |
| C | 33.32800 | 35.05900 | 44.40200 |
| O | 32.77200 | 35.51900 | 43.37500 |
| C | 31.81780 | 34.71010 | 46.48230 |
| C | 30.99120 | 35.92350 | 46.08520 |
| O | 30.91980 | 33.77320 | 47.10790 |
| H | 33.47770 | 32.73370 | 46.68890 |
| H | 31.73850 | 33.67620 | 44.58290 |
| H | 32.59740 | 35.00860 | 47.20950 |
| H | 31.46510 | 33.12250 | 47.60680 |
| H | 30.50000 | 36.33590 | 46.97670 |
| H | 30.22240 | 35.63640 | 45.35420 |
| H | 31.61680 | 36.70350 | 45.63330 |
| N | 34.61300 | 35.31200 | 44.78500 |
| H | 35.05170 | 34.70490 | 45.47020 |
| C | 23.94400 | 27.94100 | 45.50200 |
| O | 23.57900 | 28.09500 | 44.33100 |
| N | 25.20400 | 28.15800 | 45.87400 |
| C | 26.22300 | 28.52000 | 44.89200 |
| C | 26.47000 | 27.38700 | 43.85800 |
| O | 26.67800 | 27.71100 | 42.68900 |
| C | 27.58220 | 28.77240 | 45.56940 |
| C | 27.68670 | 29.94120 | 46.54120 |
| O | 27.49620 | 31.11770 | 46.13350 |
| O | 28.00860 | 29.63420 | 47.75130 |
| H | 25.47060 | 28.03380 | 46.84860 |
| H | 25.89180 | 29.38720 | 44.30750 |
| H | 27.90980 | 27.86150 | 46.08790 |
| H | 28.30570 | 28.97920 | 44.76490 |
| N | 26.43600 | 26.10900 | 44.22600 |
| H | 26.15680 | 25.81910 | 45.15740 |
| C | 22.02800 | 35.96100 | 51.62100 |
| O | 21.52500 | 35.67200 | 52.73400 |
| N | 23.04900 | 35.23800 | 51.14800 |
| C | 23.74700 | 34.28100 | 52.02300 |
| C | 24.10700 | 34.76800 | 53.38500 |
| O | 23.80400 | 34.05500 | 54.34000 |
| C | 25.03070 | 33.78970 | 51.30210 |
| C | 24.79360 | 33.01470 | 49.99620 |
| C | 26.08770 | 32.50140 | 49.33530 |
| C | 26.90390 | 33.59730 | 48.64480 |

| | | | |
|---|---|---|---|
| N | 28.15690 | 33.01450 | 48.06200 |
| H | 23.49670 | 35.52340 | 50.27920 |
| H | 23.07720 | 33.42790 | 52.19970 |
| H | 25.68050 | 34.66390 | 51.12390 |
| H | 25.57340 | 33.14220 | 52.00840 |
| H | 24.14840 | 32.15030 | 50.21680 |
| H | 24.23850 | 33.62650 | 49.26610 |
| H | 26.71600 | 31.99650 | 50.08790 |
| H | 25.82670 | 31.74380 | 48.58060 |
| H | 26.32680 | 34.04070 | 47.82170 |
| H | 27.20470 | 34.38870 | 49.34390 |
| H | 28.72550 | 32.52790 | 48.81980 |
| H | 28.78700 | 33.71070 | 47.63250 |
| H | 27.91950 | 32.29910 | 47.32180 |
| N | 24.74100 | 35.92700 | 53.53200 |
| H | 24.85830 | 36.57400 | 52.75900 |
| O | 25.02000 | 32.63400 | 45.94200 |
| H | 25.77700 | 32.01140 | 45.96070 |
| H | 25.05490 | 32.99600 | 45.03880 |
| O | 27.76100 | 32.56500 | 43.56800 |
| H | 28.00350 | 31.99140 | 42.82160 |
| H | 27.75500 | 31.96750 | 44.34620 |
| H | 35.81670 | 29.27130 | 51.07340 |
| H | 33.84130 | 33.75200 | 50.00390 |
| H | 34.49780 | 31.23060 | 45.31360 |
| H | 35.20950 | 35.82450 | 44.14250 |
| H | 23.26710 | 27.63350 | 46.32770 |
| H | 26.54200 | 25.40040 | 43.50750 |
| H | 21.65600 | 36.76450 | 50.95380 |
| H | 24.89010 | 36.27880 | 54.47240 |

**Table S12.** Step 12 of the relaxed surface scan for proton transfer from complex 2 to complex 3 with explicit waters.

| | | | |
|---|---|---|---|
| C | 35.07600 | 29.61800 | 50.32400 |
| O | 35.29000 | 29.74100 | 49.12000 |
| N | 33.82600 | 29.84000 | 50.82400 |
| C | 32.80700 | 30.46300 | 50.04700 |
| C | 33.11400 | 31.92700 | 49.53200 |
| O | 32.91300 | 32.28500 | 48.36700 |
| C | 31.44970 | 30.44120 | 50.77850 |
| C | 30.28750 | 30.79530 | 49.84460 |
| O | 29.72440 | 31.90320 | 49.94550 |
| O | 29.98840 | 29.85840 | 49.00040 |
| H | 33.70440 | 29.79330 | 51.83450 |
| H | 32.70070 | 29.90220 | 49.10580 |
| H | 31.28480 | 29.42030 | 51.15240 |
| H | 31.45160 | 31.14070 | 51.62550 |
| H | 28.99690 | 29.93720 | 48.37490 |
| N | 33.63800 | 32.82200 | 50.35700 |
| H | 33.87440 | 32.61030 | 51.32170 |
| C | 33.90300 | 32.02700 | 44.81800 |
| O | 33.71700 | 32.08700 | 43.59100 |
| N | 33.39200 | 32.93300 | 45.68900 |
| C | 32.51000 | 34.06700 | 45.26000 |
| C | 33.32800 | 35.05900 | 44.40200 |
| O | 32.77200 | 35.51900 | 43.37500 |
| C | 31.81550 | 34.70890 | 46.48140 |
| C | 30.99200 | 35.92430 | 46.08410 |
| O | 30.91380 | 33.77210 | 47.10170 |
| H | 33.47850 | 32.73420 | 46.68880 |
| H | 31.73920 | 33.67600 | 44.58220 |
| H | 32.59320 | 35.00460 | 47.21170 |
| H | 31.45590 | 33.11990 | 47.60160 |
| H | 30.49810 | 36.33520 | 46.97490 |
| H | 30.22560 | 35.64010 | 45.34960 |
| H | 31.62040 | 36.70460 | 45.63650 |
| N | 34.61300 | 35.31200 | 44.78500 |
| H | 35.05080 | 34.70730 | 45.47280 |
| C | 23.94400 | 27.94100 | 45.50200 |
| O | 23.57900 | 28.09500 | 44.33100 |
| N | 25.20400 | 28.15800 | 45.87400 |
| C | 26.22300 | 28.52000 | 44.89200 |
| C | 26.47000 | 27.38700 | 43.85800 |
| O | 26.67800 | 27.71100 | 42.68900 |
| C | 27.57940 | 28.77840 | 45.57790 |
| C | 27.66470 | 29.96450 | 46.53330 |
| O | 27.45140 | 31.12870 | 46.09380 |
| O | 27.97800 | 29.70340 | 47.75180 |
| H | 25.47030 | 28.03720 | 46.84910 |
| H | 25.89290 | 29.38780 | 44.30780 |
| H | 27.90060 | 27.87460 | 46.11320 |
| H | 28.31050 | 28.97350 | 44.77770 |
| N | 26.43600 | 26.10900 | 44.22600 |
| H | 26.15600 | 25.81960 | 45.15740 |
| C | 22.02800 | 35.96100 | 51.62100 |
| O | 21.52500 | 35.67200 | 52.73400 |
| N | 23.04900 | 35.23800 | 51.14800 |
| C | 23.74700 | 34.28100 | 52.02300 |
| C | 24.10700 | 34.76800 | 53.38500 |
| O | 23.80400 | 34.05500 | 54.34000 |

| | | | |
|---|---|---|---|
| C | 25.02970 | 33.78870 | 51.30050 |
| C | 24.78860 | 33.01070 | 49.99700 |
| C | 26.07940 | 32.49870 | 49.32870 |
| C | 26.89000 | 33.59590 | 48.63410 |
| N | 28.12860 | 33.01300 | 48.02120 |
| H | 23.49710 | 35.52370 | 50.27950 |
| H | 23.07680 | 33.42820 | 52.20000 |
| H | 25.67840 | 34.66300 | 51.11850 |
| H | 25.57440 | 33.14330 | 52.00720 |
| H | 24.14620 | 32.14540 | 50.22250 |
| H | 24.22840 | 33.62030 | 49.26900 |
| H | 26.71260 | 31.99280 | 50.07650 |
| H | 25.81480 | 31.74170 | 48.57490 |
| H | 26.30120 | 34.04930 | 47.82510 |
| H | 27.20790 | 34.37920 | 49.33490 |
| H | 28.70810 | 32.51150 | 48.75620 |
| H | 28.75530 | 33.71290 | 47.59380 |
| H | 27.87470 | 32.30760 | 47.27500 |
| N | 24.74100 | 35.92700 | 53.53200 |
| H | 24.85780 | 36.57400 | 52.75890 |
| O | 25.02000 | 32.63400 | 45.94200 |
| H | 25.77440 | 32.00670 | 45.94810 |
| H | 25.04690 | 33.00280 | 45.04150 |
| O | 27.76100 | 32.56500 | 43.56800 |
| H | 27.98350 | 31.99050 | 42.81620 |
| H | 27.73560 | 31.96070 | 44.34140 |
| H | 35.81640 | 29.27100 | 51.07340 |
| H | 33.84270 | 33.75150 | 50.00350 |
| H | 34.49790 | 31.23070 | 45.31370 |
| H | 35.20970 | 35.82510 | 44.14320 |
| H | 23.26680 | 27.63380 | 46.32770 |
| H | 26.54050 | 25.40020 | 43.50740 |
| H | 21.65590 | 36.76440 | 50.95380 |
| H | 24.88950 | 36.27910 | 54.47240 |

**Table S14.** Step 11 of the relaxed surface scan for proton transfer from complex 2 to complex 3 with explicit waters.

| | | | |
|---|---|---|---|
| C | 35.07600 | 29.61800 | 50.32400 |
| O | 35.29000 | 29.74100 | 49.12000 |
| N | 33.82600 | 29.84000 | 50.82400 |
| C | 32.80700 | 30.46300 | 50.04700 |
| C | 33.11400 | 31.92700 | 49.53200 |
| O | 32.91300 | 32.28500 | 48.36700 |
| C | 31.43500 | 30.39620 | 50.74760 |
| C | 30.29530 | 30.72640 | 49.77980 |
| O | 29.86350 | 31.89040 | 49.70580 |
| O | 29.87490 | 29.70600 | 49.08890 |
| H | 33.70480 | 29.79300 | 51.83460 |
| H | 32.72790 | 29.90750 | 49.09870 |
| H | 31.29720 | 29.37090 | 51.11730 |
| H | 31.39360 | 31.09740 | 51.59290 |
| H | 28.95890 | 29.87050 | 48.42230 |
| N | 33.63800 | 32.82200 | 50.35700 |
| H | 33.87300 | 32.61070 | 51.32210 |
| C | 33.90300 | 32.02700 | 44.81800 |
| O | 33.71700 | 32.08700 | 43.59100 |
| N | 33.39200 | 32.93300 | 45.68900 |
| C | 32.51000 | 34.06700 | 45.26000 |
| C | 33.32800 | 35.05900 | 44.40200 |
| O | 32.77200 | 35.51900 | 43.37500 |
| C | 31.81620 | 34.70720 | 46.48450 |
| C | 30.98000 | 35.91500 | 46.08830 |
| O | 30.92360 | 33.77180 | 47.11140 |
| H | 33.47610 | 32.73440 | 46.68930 |
| H | 31.73870 | 33.67580 | 44.58290 |
| H | 32.59890 | 35.01240 | 47.20600 |
| H | 31.46350 | 33.12470 | 47.61950 |
| H | 30.51420 | 36.34330 | 46.98580 |
| H | 30.18570 | 35.60740 | 45.39380 |
| H | 31.58860 | 36.68660 | 45.60030 |
| N | 34.61300 | 35.31200 | 44.78500 |
| H | 35.04970 | 34.71080 | 45.47650 |
| C | 23.94400 | 27.94100 | 45.50200 |
| O | 23.57900 | 28.09500 | 44.33100 |
| N | 25.20400 | 28.15800 | 45.87400 |
| C | 26.22300 | 28.52000 | 44.89200 |
| C | 26.47000 | 27.38700 | 43.85800 |
| O | 26.67800 | 27.71100 | 42.68900 |
| C | 27.56030 | 28.81900 | 45.60060 |
| C | 27.55180 | 30.04500 | 46.51650 |
| O | 27.17510 | 31.15050 | 46.04050 |
| O | 27.91900 | 29.89530 | 47.73600 |
| H | 25.47100 | 28.03330 | 46.84830 |
| H | 25.88710 | 29.38340 | 44.30560 |
| H | 27.89020 | 27.94040 | 46.17250 |
| H | 28.30970 | 29.01250 | 44.81750 |
| N | 26.43600 | 26.10900 | 44.22600 |
| H | 26.14880 | 25.82110 | 45.15570 |
| C | 22.02800 | 35.96100 | 51.62100 |
| O | 21.52500 | 35.67200 | 52.73400 |

| | | | |
|---|---|---|---|
| N | 23.04900 | 35.23800 | 51.14800 |
| C | 23.74700 | 34.28100 | 52.02300 |
| C | 24.10700 | 34.76800 | 53.38500 |
| O | 23.80400 | 34.05500 | 54.34000 |
| C | 25.03140 | 33.79250 | 51.30440 |
| C | 24.78980 | 33.02210 | 49.99900 |
| C | 26.07610 | 32.51020 | 49.32760 |
| C | 26.93170 | 33.61820 | 48.70830 |
| N | 28.06770 | 33.01320 | 47.94300 |
| H | 23.49550 | 35.52190 | 50.27800 |
| H | 23.07790 | 33.42720 | 52.19940 |
| H | 25.67980 | 34.66750 | 51.12550 |
| H | 25.57510 | 33.14460 | 52.00970 |
| H | 24.14180 | 32.15890 | 50.21800 |
| H | 24.23630 | 33.64040 | 49.27290 |
| H | 26.67850 | 31.93470 | 50.04980 |
| H | 25.79910 | 31.80900 | 48.52510 |
| H | 26.33210 | 34.20020 | 47.99650 |
| H | 27.35980 | 34.28830 | 49.46540 |
| H | 28.71940 | 32.49340 | 48.59130 |
| H | 28.63730 | 33.69830 | 47.42320 |
| H | 27.69620 | 32.32030 | 47.23810 |
| N | 24.74100 | 35.92700 | 53.53200 |
| H | 24.85720 | 36.57410 | 52.75890 |
| O | 25.29160 | 33.47740 | 45.90310 |
| H | 25.47030 | 32.52040 | 45.97220 |
| H | 26.09710 | 33.79740 | 45.43910 |
| O | 27.99540 | 33.60870 | 44.96220 |
| H | 28.43090 | 33.80750 | 44.11610 |
| H | 27.90330 | 32.63120 | 44.98590 |
| H | 35.81600 | 29.27050 | 51.07350 |
| H | 33.84470 | 33.75070 | 50.00270 |
| H | 34.49790 | 31.23060 | 45.31330 |
| H | 35.20990 | 35.82810 | 44.14580 |
| H | 23.26620 | 27.63380 | 46.32760 |
| H | 26.53180 | 25.40020 | 43.50620 |
| H | 21.65600 | 36.76440 | 50.95370 |
| H | 24.88910 | 36.27930 | 54.47230 |

**Table S14.** Step 10 of the relaxed surface scan for proton transfer from complex 2 to complex 3 with explicit waters.

| | | | |
|---|---|---|---|
| C | 35.07600 | 29.61800 | 50.32400 |
| O | 35.29000 | 29.74100 | 49.12000 |
| N | 33.82600 | 29.84000 | 50.82400 |
| C | 32.80700 | 30.46300 | 50.04700 |
| C | 33.11400 | 31.92700 | 49.53200 |
| O | 32.91300 | 32.28500 | 48.36700 |
| C | 31.44670 | 30.42870 | 50.77300 |
| C | 30.28710 | 30.76680 | 49.83200 |
| O | 29.74510 | 31.88600 | 49.88530 |
| O | 29.96690 | 29.79680 | 49.02770 |
| H | 33.70470 | 29.79350 | 51.83460 |
| H | 32.70730 | 29.90320 | 49.10410 |
| H | 31.29330 | 29.40780 | 51.15080 |
| H | 31.43580 | 31.13290 | 51.61610 |
| H | 29.01520 | 29.90550 | 48.39720 |
| N | 33.63800 | 32.82200 | 50.35700 |
| H | 33.87550 | 32.61010 | 51.32140 |
| C | 33.90300 | 32.02700 | 44.81800 |
| O | 33.71700 | 32.08700 | 43.59100 |
| N | 33.39200 | 32.93300 | 45.68900 |
| C | 32.51000 | 34.06700 | 45.26000 |
| C | 33.32800 | 35.05900 | 44.40200 |
| O | 32.77200 | 35.51900 | 43.37500 |
| C | 31.81480 | 34.70830 | 46.48120 |
| C | 30.99420 | 35.92590 | 46.08440 |
| O | 30.91000 | 33.77260 | 47.09870 |
| H | 33.47920 | 32.73520 | 46.68890 |
| H | 31.73950 | 33.67590 | 44.58200 |
| H | 32.59180 | 35.00160 | 47.21320 |
| H | 31.44910 | 33.11780 | 47.59800 |
| H | 30.49870 | 36.33550 | 46.97480 |
| H | 30.22920 | 35.64440 | 45.34730 |
| H | 31.62480 | 36.70610 | 45.64000 |
| N | 34.61300 | 35.31200 | 44.78500 |
| H | 35.05080 | 34.70720 | 45.47270 |
| C | 23.94400 | 27.94100 | 45.50200 |
| O | 23.57900 | 28.09500 | 44.33100 |
| N | 25.20400 | 28.15800 | 45.87400 |
| C | 26.22300 | 28.52000 | 44.89200 |
| C | 26.47000 | 27.38700 | 43.85800 |
| O | 26.67800 | 27.71100 | 42.68900 |
| C | 27.57800 | 28.78190 | 45.58220 |
| C | 27.65020 | 29.97840 | 46.52720 |
| O | 27.41820 | 31.13360 | 46.06620 |
| O | 27.96150 | 29.74990 | 47.74870 |
| H | 25.46990 | 28.03950 | 46.84940 |
| H | 25.89380 | 29.38810 | 44.30790 |
| H | 27.89650 | 27.88260 | 46.12670 |
| H | 28.31280 | 28.97120 | 44.78410 |

| | | | |
|---|---|---|---|
| N | 26.43600 | 26.10900 | 44.22600 |
| H | 26.15580 | 25.81990 | 45.15740 |
| C | 22.02800 | 35.96100 | 51.62100 |
| O | 21.52500 | 35.67200 | 52.73400 |
| N | 23.04900 | 35.23800 | 51.14800 |
| C | 23.74700 | 34.28100 | 52.02300 |
| C | 24.10700 | 34.76800 | 53.38500 |
| O | 23.80400 | 34.05500 | 54.34000 |
| C | 25.02920 | 33.78850 | 51.29930 |
| C | 24.78560 | 33.00940 | 49.99680 |
| C | 26.07380 | 32.49760 | 49.32310 |
| C | 26.88120 | 33.59440 | 48.62430 |
| N | 28.10800 | 33.00960 | 47.98960 |
| H | 23.49740 | 35.52410 | 50.27970 |
| H | 23.07680 | 33.42820 | 52.20010 |
| H | 25.67740 | 34.66290 | 51.11540 |
| H | 25.57520 | 33.14380 | 52.00560 |
| H | 24.14480 | 32.14360 | 50.22510 |
| H | 24.22190 | 33.61770 | 49.27040 |
| H | 26.71030 | 31.99110 | 50.06760 |
| H | 25.80550 | 31.74100 | 48.57040 |
| H | 26.28440 | 34.05480 | 47.82520 |
| H | 27.21240 | 34.37220 | 49.32520 |
| H | 28.69430 | 32.49850 | 48.70830 |
| H | 28.73230 | 33.71080 | 47.56140 |
| H | 27.84120 | 32.31010 | 47.24040 |
| N | 24.74100 | 35.92700 | 53.53200 |
| H | 24.85750 | 36.57410 | 52.75890 |
| O | 25.02000 | 32.63400 | 45.94200 |
| H | 25.77140 | 32.00220 | 45.93840 |
| H | 25.04160 | 33.00830 | 45.04380 |
| O | 27.76100 | 32.56500 | 43.56800 |
| H | 27.94370 | 31.98650 | 42.80860 |
| H | 27.71780 | 31.95610 | 44.33760 |
| H | 35.81630 | 29.27090 | 51.07350 |
| H | 33.84440 | 33.75090 | 50.00280 |
| H | 34.49800 | 31.23070 | 45.31360 |
| H | 35.20980 | 35.82480 | 44.14300 |
| H | 23.26680 | 27.63390 | 46.32770 |
| H | 26.53960 | 25.40010 | 43.50740 |
| H | 21.65600 | 36.76450 | 50.95380 |
| H | 24.88900 | 36.27930 | 54.47230 |

**Table S15.** Step 9 of the relaxed surface scan for proton transfer from complex 2 to complex 3 with explicit waters.

| | | | |
|---|---|---|---|
| C | 35.07600 | 29.61800 | 50.32400 |
| O | 35.29000 | 29.74100 | 49.12000 |
| N | 33.82600 | 29.84000 | 50.82400 |
| C | 32.80700 | 30.46300 | 50.04700 |
| C | 33.11400 | 31.92700 | 49.53200 |
| O | 32.91300 | 32.28500 | 48.36700 |
| C | 31.43720 | 30.39770 | 50.75300 |
| C | 30.29580 | 30.72580 | 49.78920 |
| O | 29.86770 | 31.88770 | 49.70040 |
| O | 29.86810 | 29.69500 | 49.11130 |
| H | 33.70490 | 29.79310 | 51.83460 |
| H | 32.72570 | 29.90700 | 49.09920 |
| H | 31.30060 | 29.37370 | 51.12630 |
| H | 31.39860 | 31.10180 | 51.59600 |
| H | 28.98120 | 29.86550 | 48.45400 |
| N | 33.63800 | 32.82200 | 50.35700 |
| H | 33.87420 | 32.61070 | 51.32180 |
| C | 33.90300 | 32.02700 | 44.81800 |
| O | 33.71700 | 32.08700 | 43.59100 |
| N | 33.39200 | 32.93300 | 45.68900 |
| C | 32.51000 | 34.06700 | 45.26000 |
| C | 33.32800 | 35.05900 | 44.40200 |
| O | 32.77200 | 35.51900 | 43.37500 |
| C | 31.81440 | 34.70610 | 46.48380 |
| C | 30.97780 | 35.91350 | 46.08710 |
| O | 30.92150 | 33.76950 | 47.10850 |
| H | 33.47650 | 32.73450 | 46.68930 |
| H | 31.73930 | 33.67550 | 44.58230 |
| H | 32.59580 | 35.01150 | 47.20680 |
| H | 31.46110 | 33.12270 | 47.61710 |
| H | 30.51060 | 36.34120 | 46.98420 |
| H | 30.18470 | 35.60560 | 45.39140 |
| H | 31.58650 | 36.68560 | 45.60010 |
| N | 34.61300 | 35.31200 | 44.78500 |
| H | 35.04990 | 34.71050 | 45.47600 |
| C | 23.94400 | 27.94100 | 45.50200 |
| O | 23.57900 | 28.09500 | 44.33100 |
| N | 25.20400 | 28.15800 | 45.87400 |
| C | 26.22300 | 28.52000 | 44.89200 |
| C | 26.47000 | 27.38700 | 43.85800 |
| O | 26.67800 | 27.71100 | 42.68900 |
| C | 27.55950 | 28.82260 | 45.59940 |
| C | 27.54530 | 30.04720 | 46.51850 |
| O | 27.16710 | 31.15330 | 46.03820 |
| O | 27.91050 | 29.89850 | 47.73550 |

| | | | |
|---|---|---|---|
| H | 25.47140 | 28.03420 | 46.84840 |
| H | 25.88560 | 29.38260 | 44.30520 |
| H | 27.89240 | 27.94440 | 46.17020 |
| H | 28.30710 | 29.02000 | 44.81570 |
| N | 26.43600 | 26.10900 | 44.22600 |
| H | 26.14890 | 25.82150 | 45.15580 |
| C | 22.02800 | 35.96100 | 51.62100 |
| O | 21.52500 | 35.67200 | 52.73400 |
| N | 23.04900 | 35.23800 | 51.14800 |
| C | 23.74700 | 34.28100 | 52.02300 |
| C | 24.10700 | 34.76800 | 53.38500 |
| O | 23.80400 | 34.05500 | 54.34000 |
| C | 25.03070 | 33.79180 | 51.30370 |
| C | 24.78750 | 33.01940 | 49.99970 |
| C | 26.07280 | 32.50730 | 49.32670 |
| C | 26.92460 | 33.61490 | 48.70140 |
| N | 28.05890 | 33.01010 | 47.93370 |
| H | 23.49520 | 35.52160 | 50.27770 |
| H | 23.07760 | 33.42740 | 52.19960 |
| H | 25.67900 | 34.66660 | 51.12290 |
| H | 25.57530 | 33.14490 | 52.00930 |
| H | 24.14040 | 32.15630 | 50.22120 |
| H | 24.23240 | 33.63630 | 49.27360 |
| H | 26.67810 | 31.93510 | 50.04920 |
| H | 25.79540 | 31.80290 | 48.52720 |
| H | 26.32130 | 34.19310 | 47.98960 |
| H | 27.35370 | 34.28840 | 49.45490 |
| H | 28.71080 | 32.49110 | 48.57920 |
| H | 28.62710 | 33.69580 | 47.41330 |
| H | 27.68600 | 32.31550 | 47.22880 |
| N | 24.74100 | 35.92700 | 53.53200 |
| H | 24.85700 | 36.57410 | 52.75890 |
| O | 25.28010 | 33.46610 | 45.90150 |
| H | 25.46640 | 32.51040 | 45.97130 |
| H | 26.08230 | 33.79010 | 45.43450 |
| O | 27.98320 | 33.60310 | 44.95460 |
| H | 28.41790 | 33.79670 | 44.10690 |
| H | 27.88930 | 32.62550 | 44.98380 |
| H | 35.81610 | 29.27080 | 51.07350 |
| H | 33.84500 | 33.75050 | 50.00240 |
| H | 34.49800 | 31.23060 | 45.31330 |
| H | 35.20980 | 35.82800 | 44.14570 |
| H | 23.26620 | 27.63360 | 46.32750 |
| H | 26.53140 | 25.40010 | 43.50630 |
| H | 21.65590 | 36.76430 | 50.95370 |
| H | 24.88880 | 36.27940 | 54.47230 |

**Table S16.** Step 8 of the relaxed surface scan for proton transfer from complex 2 to complex 3 with explicit waters.

| | | | |
|---|---|---|---|
| C | 35.07600 | 29.61800 | 50.32400 |
| O | 35.29000 | 29.74100 | 49.12000 |
| N | 33.82600 | 29.84000 | 50.82400 |
| C | 32.80700 | 30.46300 | 50.04700 |
| C | 33.11400 | 31.92700 | 49.53200 |
| O | 32.91300 | 32.28500 | 48.36700 |
| C | 31.44630 | 30.42250 | 50.77310 |
| C | 30.28660 | 30.74870 | 49.83030 |
| O | 29.76610 | 31.87730 | 49.84280 |
| O | 29.94180 | 29.75260 | 49.06380 |
| H | 33.70500 | 29.79380 | 51.83460 |
| H | 32.71000 | 29.90360 | 49.10350 |
| H | 31.30110 | 29.40280 | 51.15680 |
| H | 31.43020 | 31.13190 | 51.61170 |
| H | 29.02120 | 29.88080 | 48.43170 |
| N | 33.63800 | 32.82200 | 50.35700 |
| H | 33.87680 | 32.60990 | 51.32100 |
| C | 33.90300 | 32.02700 | 44.81800 |
| O | 33.71700 | 32.08700 | 43.59100 |
| N | 33.39200 | 32.93300 | 45.68900 |
| C | 32.51000 | 34.06700 | 45.26000 |
| C | 33.32800 | 35.05900 | 44.40200 |
| O | 32.77200 | 35.51900 | 43.37500 |
| C | 31.81420 | 34.70790 | 46.48090 |
| C | 30.99580 | 35.92720 | 46.08430 |
| O | 30.90690 | 33.77310 | 47.09590 |
| H | 33.47940 | 32.73520 | 46.68890 |
| H | 31.73970 | 33.67580 | 44.58180 |
| H | 32.59070 | 34.99910 | 47.21430 |
| H | 31.44360 | 33.11620 | 47.59450 |
| H | 30.49900 | 36.33600 | 46.97430 |
| H | 30.23200 | 35.64780 | 45.34520 |
| H | 31.62840 | 36.70730 | 45.64240 |
| N | 34.61300 | 35.31200 | 44.78500 |
| H | 35.05070 | 34.70680 | 45.47240 |
| C | 23.94400 | 27.94100 | 45.50200 |
| O | 23.57900 | 28.09500 | 44.33100 |
| N | 25.20400 | 28.15800 | 45.87400 |
| C | 26.22300 | 28.52000 | 44.89200 |
| C | 26.47000 | 27.38700 | 43.85800 |
| O | 26.67800 | 27.71100 | 42.68900 |
| C | 27.57670 | 28.78420 | 45.58550 |

| | | | |
|---|---|---|---|
| C | 27.63930 | 29.98830 | 46.52280 |
| O | 27.39480 | 31.13710 | 46.04720 |
| O | 27.94900 | 29.78130 | 47.74530 |
| H | 25.46970 | 28.04140 | 46.84970 |
| H | 25.89420 | 29.38820 | 44.30800 |
| H | 27.89220 | 27.88820 | 46.13730 |
| H | 28.31510 | 28.96830 | 44.78960 |
| N | 26.43600 | 26.10900 | 44.22600 |
| H | 26.15530 | 25.82010 | 45.15730 |
| C | 22.02800 | 35.96100 | 51.62100 |
| O | 21.52500 | 35.67200 | 52.73400 |
| N | 23.04900 | 35.23800 | 51.14800 |
| C | 23.74700 | 34.28100 | 52.02300 |
| C | 24.10700 | 34.76800 | 53.38500 |
| O | 23.80400 | 34.05500 | 54.34000 |
| C | 25.02910 | 33.78830 | 51.29860 |
| C | 24.78430 | 33.00820 | 49.99680 |
| C | 26.07130 | 32.49610 | 49.32060 |
| C | 26.87690 | 33.59260 | 48.61920 |
| N | 28.09520 | 33.00690 | 47.96940 |
| H | 23.49720 | 35.52370 | 50.27950 |
| H | 23.07690 | 33.42820 | 52.20010 |
| H | 25.67670 | 34.66280 | 51.11350 |
| H | 25.57580 | 33.14430 | 52.00490 |
| H | 24.14420 | 32.14230 | 50.22660 |
| H | 24.21930 | 33.61590 | 49.27100 |
| H | 26.70940 | 31.98990 | 50.06390 |
| H | 25.80150 | 31.73920 | 48.56900 |
| H | 26.27470 | 34.05830 | 47.82730 |
| H | 27.21720 | 34.36640 | 49.32020 |
| H | 28.68820 | 32.49020 | 48.67590 |
| H | 28.71680 | 33.70910 | 47.53930 |
| H | 27.81890 | 32.31120 | 47.21880 |
| N | 24.74100 | 35.92700 | 53.53200 |
| H | 24.85720 | 36.57410 | 52.75890 |
| O | 25.02000 | 32.63400 | 45.94200 |
| H | 25.76960 | 31.99930 | 45.93210 |
| H | 25.03510 | 33.00850 | 45.04380 |
| O | 27.76100 | 32.56500 | 43.56800 |
| H | 27.92770 | 31.98460 | 42.80640 |
| H | 27.70670 | 31.95490 | 44.33640 |
| H | 35.81620 | 29.27080 | 51.07350 |
| H | 33.84570 | 33.75030 | 50.00210 |
| H | 34.49810 | 31.23070 | 45.31360 |
| H | 35.20980 | 35.82440 | 44.14270 |
| H | 23.26660 | 27.63410 | 46.32770 |
| H | 26.53910 | 25.40010 | 43.50740 |
| H | 21.65590 | 36.76450 | 50.95380 |
| H | 24.88870 | 36.27950 | 54.47230 |

**Table S17.** Step 7 of the relaxed surface scan for proton transfer from complex 2 to complex 3 with explicit waters.

| | | | |
|---|---|---|---|
| C | 35.07600 | 29.61800 | 50.32400 |
| O | 35.29000 | 29.74100 | 49.12000 |
| N | 33.82600 | 29.84000 | 50.82400 |
| C | 32.80700 | 30.46300 | 50.04700 |
| C | 33.11400 | 31.92700 | 49.53200 |
| O | 32.91300 | 32.28500 | 48.36700 |
| C | 31.44790 | 30.42320 | 50.77660 |
| C | 30.28710 | 30.74870 | 49.83730 |
| O | 29.76740 | 31.87580 | 49.84230 |
| O | 29.93920 | 29.74560 | 49.07650 |
| H | 33.70500 | 29.79370 | 51.83460 |
| H | 32.70840 | 29.90320 | 49.10400 |
| H | 31.30350 | 29.40420 | 51.16240 |
| H | 31.43350 | 31.13450 | 51.61360 |
| H | 29.03570 | 29.88090 | 48.45090 |
| N | 33.63800 | 32.82200 | 50.35700 |
| H | 33.87740 | 32.61000 | 51.32090 |
| C | 33.90300 | 32.02700 | 44.81800 |
| O | 33.71700 | 32.08700 | 43.59100 |
| N | 33.39200 | 32.93300 | 45.68900 |
| C | 32.51000 | 34.06700 | 45.26000 |
| C | 33.32800 | 35.05900 | 44.40200 |
| O | 32.77200 | 35.51900 | 43.37500 |
| C | 31.81370 | 34.70790 | 46.48060 |
| C | 30.99600 | 35.92740 | 46.08380 |
| O | 30.90560 | 33.77300 | 47.09450 |
| H | 33.47930 | 32.73510 | 46.68880 |
| H | 31.73980 | 33.67580 | 44.58160 |
| H | 32.58970 | 34.99860 | 47.21470 |
| H | 31.44200 | 33.11600 | 47.59310 |
| H | 30.49830 | 36.33590 | 46.97350 |
| H | 30.23290 | 35.64860 | 45.34370 |
| H | 31.62900 | 36.70780 | 45.64300 |
| N | 34.61300 | 35.31200 | 44.78500 |
| H | 35.05070 | 34.70680 | 45.47240 |
| C | 23.94400 | 27.94100 | 45.50200 |
| O | 23.57900 | 28.09500 | 44.33100 |
| N | 25.20400 | 28.15800 | 45.87400 |

| | | | |
|---|---|---|---|
| C | 26.22300 | 28.52000 | 44.89200 |
| C | 26.47000 | 27.38700 | 43.85800 |
| O | 26.67800 | 27.71100 | 42.68900 |
| C | 27.57640 | 28.78790 | 45.58390 |
| C | 27.63400 | 29.99090 | 46.52370 |
| O | 27.38970 | 31.14060 | 46.04510 |
| O | 27.93980 | 29.78550 | 47.74500 |
| H | 25.46970 | 28.04290 | 46.84990 |
| H | 25.89280 | 29.38740 | 44.30750 |
| H | 27.89540 | 27.89250 | 46.13470 |
| H | 28.31270 | 28.97540 | 44.78700 |
| N | 26.43600 | 26.10900 | 44.22600 |
| H | 26.15540 | 25.82030 | 45.15740 |
| C | 22.02800 | 35.96100 | 51.62100 |
| O | 21.52500 | 35.67200 | 52.73400 |
| N | 23.04900 | 35.23800 | 51.14800 |
| C | 23.74700 | 34.28100 | 52.02300 |
| C | 24.10700 | 34.76800 | 53.38500 |
| O | 23.80400 | 34.05500 | 54.34000 |
| C | 25.02830 | 33.78770 | 51.29780 |
| C | 24.78150 | 33.00680 | 49.99670 |
| C | 26.06700 | 32.49430 | 49.31810 |
| C | 26.87130 | 33.59030 | 48.61470 |
| N | 28.08880 | 33.00440 | 47.96320 |
| H | 23.49720 | 35.52370 | 50.27950 |
| H | 23.07660 | 33.42840 | 52.20040 |
| H | 25.67600 | 34.66190 | 51.11150 |
| H | 25.57560 | 33.14370 | 52.00380 |
| H | 24.14180 | 32.14110 | 50.22820 |
| H | 24.21500 | 33.61410 | 49.27160 |
| H | 26.70640 | 31.98810 | 50.06030 |
| H | 25.79580 | 31.73700 | 48.56730 |
| H | 26.26810 | 34.05510 | 47.82300 |
| H | 27.21230 | 34.36480 | 49.31460 |
| H | 28.68060 | 32.48790 | 48.66830 |
| H | 28.71060 | 33.70700 | 47.53400 |
| H | 27.81180 | 32.30790 | 47.21120 |
| N | 24.74100 | 35.92700 | 53.53200 |
| H | 24.85720 | 36.57410 | 52.75890 |
| O | 25.02000 | 32.63400 | 45.94200 |
| H | 25.76990 | 31.99910 | 45.93170 |
| H | 25.03320 | 33.00690 | 45.04320 |
| O | 27.76100 | 32.56500 | 43.56800 |
| H | 27.91950 | 31.98260 | 42.80610 |
| H | 27.70320 | 31.95560 | 44.33700 |
| H | 35.81630 | 29.27120 | 51.07360 |
| H | 33.84560 | 33.75030 | 50.00200 |
| H | 34.49810 | 31.23070 | 45.31360 |
| H | 35.20980 | 35.82430 | 44.14270 |
| H | 23.26660 | 27.63420 | 46.32770 |
| H | 26.53870 | 25.40000 | 43.50740 |
| H | 21.65600 | 36.76450 | 50.95380 |
| H | 24.88850 | 36.27960 | 54.47230 |

**Table S18.** Step 6 of the relaxed surface scan for proton transfer from complex 2 to complex 3 with explicit waters.

| | | | |
|---|---|---|---|
| C | 35.07600 | 29.61800 | 50.32400 |
| O | 35.29000 | 29.74100 | 49.12000 |
| N | 33.82600 | 29.84000 | 50.82400 |
| C | 32.80700 | 30.46300 | 50.04700 |
| C | 33.11400 | 31.92700 | 49.53200 |
| O | 32.91300 | 32.28500 | 48.36700 |
| C | 31.43970 | 30.39980 | 50.75880 |
| C | 30.29650 | 30.72550 | 49.79920 |
| O | 29.87170 | 31.88560 | 49.69620 |
| O | 29.86170 | 29.68530 | 49.13440 |
| H | 33.70490 | 29.79300 | 51.83460 |
| H | 32.72330 | 29.90640 | 49.09990 |
| H | 31.30430 | 29.37710 | 51.13590 |
| H | 31.40450 | 31.10670 | 51.59940 |
| H | 28.99800 | 29.86110 | 48.48470 |
| N | 33.63800 | 32.82200 | 50.35700 |
| H | 33.87520 | 32.61060 | 51.32160 |
| C | 33.90300 | 32.02700 | 44.81800 |
| O | 33.71700 | 32.08700 | 43.59100 |
| N | 33.39200 | 32.93300 | 45.68900 |
| C | 32.51000 | 34.06700 | 45.26000 |
| C | 33.32800 | 35.05900 | 44.40200 |
| O | 32.77200 | 35.51900 | 43.37500 |
| C | 31.81270 | 34.70490 | 46.48330 |
| C | 30.97570 | 35.91190 | 46.08630 |
| O | 30.91960 | 33.76690 | 47.10580 |
| H | 33.47700 | 32.73480 | 46.68920 |
| H | 31.74000 | 33.67520 | 44.58180 |
| H | 32.59280 | 35.01020 | 47.20770 |
| H | 31.45900 | 33.12050 | 47.61500 |
| H | 30.50720 | 36.33890 | 46.98300 |
| H | 30.18360 | 35.60370 | 45.38950 |
| H | 31.58440 | 36.68460 | 45.60030 |
| N | 34.61300 | 35.31200 | 44.78500 |

| | | | |
|---|---|---|---|
| H | 35.04970 | 34.71070 | 45.47630 |
| C | 23.94400 | 27.94100 | 45.50200 |
| O | 23.57900 | 28.09500 | 44.33100 |
| N | 25.20400 | 28.15800 | 45.87400 |
| C | 26.22300 | 28.52000 | 44.89200 |
| C | 26.47000 | 27.38700 | 43.85800 |
| O | 26.67800 | 27.71100 | 42.68900 |
| C | 27.55860 | 28.82650 | 45.59800 |
| C | 27.53680 | 30.04970 | 46.51960 |
| O | 27.15910 | 31.15640 | 46.03460 |
| O | 27.89530 | 29.90120 | 47.73590 |
| H | 25.47140 | 28.03550 | 46.84860 |
| H | 25.88420 | 29.38170 | 44.30460 |
| H | 27.89540 | 27.94900 | 46.16770 |
| H | 28.30420 | 29.02820 | 44.81370 |
| N | 26.43600 | 26.10900 | 44.22600 |
| H | 26.14890 | 25.82170 | 45.15590 |
| C | 22.02800 | 35.96100 | 51.62100 |
| O | 21.52500 | 35.67200 | 52.73400 |
| N | 23.04900 | 35.23800 | 51.14800 |
| C | 23.74700 | 34.28100 | 52.02300 |
| C | 24.10700 | 34.76800 | 53.38500 |
| O | 23.80400 | 34.05500 | 54.34000 |
| C | 25.03040 | 33.79120 | 51.30380 |
| C | 24.78600 | 33.01620 | 50.00160 |
| C | 26.07090 | 32.50460 | 49.32730 |
| C | 26.91840 | 33.61200 | 48.69580 |
| N | 28.05110 | 33.00700 | 47.92580 |
| H | 23.49500 | 35.52120 | 50.27750 |
| H | 23.07750 | 33.42760 | 52.19980 |
| H | 25.67780 | 34.66610 | 51.12060 |
| H | 25.57600 | 33.14600 | 52.01020 |
| H | 24.14060 | 32.15250 | 50.22590 |
| H | 24.22860 | 33.63110 | 49.27580 |
| H | 26.67920 | 31.93640 | 50.05050 |
| H | 25.79380 | 31.79650 | 48.53120 |
| H | 26.31150 | 34.18620 | 47.98390 |
| H | 27.34850 | 34.28950 | 49.44520 |
| H | 28.70300 | 32.48840 | 48.56880 |
| H | 28.61840 | 33.69300 | 47.40480 |
| H | 27.67670 | 32.31140 | 47.22060 |
| N | 24.74100 | 35.92700 | 53.53200 |
| H | 24.85680 | 36.57420 | 52.75890 |
| O | 25.26980 | 33.45560 | 45.89820 |
| H | 25.46280 | 32.50100 | 45.96830 |
| H | 26.06920 | 33.78400 | 45.42950 |
| O | 27.97230 | 33.60030 | 44.94820 |
| H | 28.40680 | 33.79040 | 44.09970 |
| H | 27.87710 | 32.62250 | 44.98110 |
| H | 35.81600 | 29.27070 | 51.07350 |
| H | 33.84530 | 33.75040 | 50.00210 |
| H | 34.49800 | 31.23060 | 45.31330 |
| H | 35.20990 | 35.82780 | 44.14560 |
| H | 23.26620 | 27.63370 | 46.32760 |
| H | 26.53110 | 25.40000 | 43.50640 |
| H | 21.65590 | 36.76430 | 50.95360 |
| H | 24.88860 | 36.27950 | 54.47230 |

**Table S19.** Step 5 of the relaxed surface scan for proton transfer from complex 2 to complex 3 with explicit waters.

| | | | |
|---|---|---|---|
| C | 35.07600 | 29.61800 | 50.32400 |
| O | 35.29000 | 29.74100 | 49.12000 |
| N | 33.82600 | 29.84000 | 50.82400 |
| C | 32.80700 | 30.46300 | 50.04700 |
| C | 33.11400 | 31.92700 | 49.53200 |
| O | 32.91300 | 32.28500 | 48.36700 |
| C | 31.45010 | 30.42220 | 50.78170 |
| C | 30.28660 | 30.74150 | 49.84560 |
| O | 29.78420 | 31.87500 | 49.82160 |
| O | 29.91630 | 29.72110 | 49.11540 |
| H | 33.70520 | 29.79420 | 51.83470 |
| H | 32.70760 | 29.90290 | 49.10420 |
| H | 31.31030 | 29.40450 | 51.17210 |
| H | 31.43730 | 31.13670 | 51.61600 |
| H | 29.03320 | 29.86720 | 48.49090 |
| N | 33.63800 | 32.82200 | 50.35700 |
| H | 33.87860 | 32.60980 | 51.32060 |
| C | 33.90300 | 32.02700 | 44.81800 |
| O | 33.71700 | 32.08700 | 43.59100 |
| N | 33.39200 | 32.93300 | 45.68900 |
| C | 32.51000 | 34.06700 | 45.26000 |
| C | 33.32800 | 35.05900 | 44.40200 |
| O | 32.77200 | 35.51900 | 43.37500 |
| C | 31.81290 | 34.70690 | 46.48060 |
| C | 30.99620 | 35.92730 | 46.08430 |
| O | 30.90370 | 33.77170 | 47.09230 |
| H | 33.47950 | 32.73560 | 46.68890 |
| H | 31.74000 | 33.67570 | 44.58140 |
| H | 32.58820 | 34.99650 | 47.21590 |
| H | 31.43870 | 33.11340 | 47.59060 |

| | | | |
|---|---|---|---|
| H | 30.49790 | 36.33520 | 46.97400 |
| H | 30.23360 | 35.64960 | 45.34330 |
| H | 31.63020 | 36.70770 | 45.64490 |
| N | 34.61300 | 35.31200 | 44.78500 |
| H | 35.05130 | 34.70540 | 45.47080 |
| C | 23.94400 | 27.94100 | 45.50200 |
| O | 23.57900 | 28.09500 | 44.33100 |
| N | 25.20400 | 28.15800 | 45.87400 |
| C | 26.22300 | 28.52000 | 44.89200 |
| C | 26.47000 | 27.38700 | 43.85800 |
| O | 26.67800 | 27.71100 | 42.68900 |
| C | 27.57520 | 28.78970 | 45.58500 |
| C | 27.62670 | 29.99630 | 46.52110 |
| O | 27.38110 | 31.14380 | 46.03410 |
| O | 27.92700 | 29.79980 | 47.74300 |
| H | 25.46960 | 28.04440 | 46.85010 |
| H | 25.89240 | 29.38720 | 44.30730 |
| H | 27.89340 | 27.89610 | 46.13940 |
| H | 28.31340 | 28.97530 | 44.78940 |
| N | 26.43600 | 26.10900 | 44.22600 |
| H | 26.15540 | 25.82050 | 45.15740 |
| C | 22.02800 | 35.96100 | 51.62100 |
| O | 21.52500 | 35.67200 | 52.73400 |
| N | 23.04900 | 35.23800 | 51.14800 |
| C | 23.74700 | 34.28100 | 52.02300 |
| C | 24.10700 | 34.76800 | 53.38500 |
| O | 23.80400 | 34.05500 | 54.34000 |
| C | 25.02840 | 33.78750 | 51.29780 |
| C | 24.78170 | 33.00560 | 49.99730 |
| C | 26.06720 | 32.49290 | 49.31850 |
| C | 26.86960 | 33.58860 | 48.61230 |
| N | 28.08280 | 33.00260 | 47.95290 |
| H | 23.49720 | 35.52360 | 50.27940 |
| H | 23.07660 | 33.42840 | 52.20030 |
| H | 25.67590 | 34.66180 | 51.11080 |
| H | 25.57590 | 33.14430 | 52.00430 |
| H | 24.14230 | 32.13990 | 50.22950 |
| H | 24.21490 | 33.61220 | 49.27200 |
| H | 26.70770 | 31.98860 | 50.06110 |
| H | 25.79620 | 31.73410 | 48.56930 |
| H | 26.26280 | 34.05430 | 47.82400 |
| H | 27.21490 | 34.36220 | 49.31100 |
| H | 28.67920 | 32.48450 | 48.65130 |
| H | 28.70200 | 33.70620 | 47.52190 |
| H | 27.80110 | 32.30700 | 47.20090 |
| N | 24.74100 | 35.92700 | 53.53200 |
| H | 24.85680 | 36.57420 | 52.75890 |
| O | 25.02000 | 32.63400 | 45.94200 |
| H | 25.76980 | 31.99850 | 45.92670 |
| H | 25.02980 | 33.00980 | 45.04440 |
| O | 27.76100 | 32.56500 | 43.56800 |
| H | 27.89030 | 31.98160 | 42.80140 |
| H | 27.69540 | 31.95370 | 44.33540 |
| H | 35.81610 | 29.27080 | 51.07350 |
| H | 33.84600 | 33.75010 | 50.00160 |
| H | 34.49810 | 31.23070 | 45.31360 |
| H | 35.20970 | 35.82390 | 44.14220 |
| H | 23.26650 | 27.63440 | 46.32770 |
| H | 26.53890 | 25.39990 | 43.50750 |
| H | 21.65590 | 36.76440 | 50.95380 |
| H | 24.88820 | 36.27970 | 54.47230 |

**Table S20.** Step 4 of the relaxed surface scan for proton transfer from complex 2 to complex 3 with explicit waters.

| | | | |
|---|---|---|---|
| C | 35.07600 | 29.61800 | 50.32400 |
| O | 35.29000 | 29.74100 | 49.12000 |
| N | 33.82600 | 29.84000 | 50.82400 |
| C | 32.80700 | 30.46300 | 50.04700 |
| C | 33.11400 | 31.92700 | 49.53200 |
| O | 32.91300 | 32.28500 | 48.36700 |
| C | 31.44240 | 30.40280 | 50.76550 |
| C | 30.29700 | 30.72670 | 49.81070 |
| O | 29.87650 | 31.88470 | 49.69260 |
| O | 29.85240 | 29.67670 | 49.16060 |
| H | 33.70470 | 29.79270 | 51.83450 |
| H | 32.71990 | 29.90550 | 49.10080 |
| H | 31.30760 | 29.38190 | 51.14720 |
| H | 31.41140 | 31.11360 | 51.60290 |
| H | 29.00920 | 29.85850 | 48.51580 |
| N | 33.63800 | 32.82200 | 50.35700 |
| H | 33.87620 | 32.61070 | 51.32130 |
| C | 33.90300 | 32.02700 | 44.81800 |
| O | 33.71700 | 32.08700 | 43.59100 |
| N | 33.39200 | 32.93300 | 45.68900 |
| C | 32.51000 | 34.06700 | 45.26000 |
| C | 33.32800 | 35.05900 | 44.40200 |
| O | 32.77200 | 35.51900 | 43.37500 |
| C | 31.81080 | 34.70350 | 46.48290 |
| C | 30.97370 | 35.91020 | 46.08580 |
| O | 30.91730 | 33.76360 | 47.10230 |

| | | | |
|---|---|---|---|
| H | 33.47730 | 32.73490 | 46.68910 |
| H | 31.74060 | 33.67490 | 44.58120 |
| H | 32.58950 | 35.00820 | 47.20900 |
| H | 31.45620 | 33.11770 | 47.61230 |
| H | 30.50360 | 36.33620 | 46.98220 |
| H | 30.18280 | 35.60230 | 45.38760 |
| H | 31.58280 | 36.68370 | 45.60150 |
| N | 34.61300 | 35.31200 | 44.78500 |
| H | 35.04980 | 34.71060 | 45.47620 |
| C | 23.94400 | 27.94100 | 45.50200 |
| O | 23.57900 | 28.09500 | 44.33100 |
| N | 25.20400 | 28.15800 | 45.87400 |
| C | 26.22300 | 28.52000 | 44.89200 |
| C | 26.47000 | 27.38700 | 43.85800 |
| O | 26.67800 | 27.71100 | 42.68900 |
| C | 27.55780 | 28.83160 | 45.59600 |
| C | 27.52620 | 30.05310 | 46.52050 |
| O | 27.15000 | 31.16000 | 46.02980 |
| O | 27.87430 | 29.90490 | 47.73730 |
| H | 25.47120 | 28.03830 | 46.84900 |
| H | 25.88230 | 29.38060 | 44.30390 |
| H | 27.89980 | 27.95530 | 46.16440 |
| H | 28.30020 | 29.03900 | 44.81030 |
| N | 26.43600 | 26.10900 | 44.22600 |
| H | 26.14890 | 25.82200 | 45.15600 |
| C | 22.02800 | 35.96100 | 51.62100 |
| O | 21.52500 | 35.67200 | 52.73400 |
| N | 23.04900 | 35.23800 | 51.14800 |
| C | 23.74700 | 34.28100 | 52.02300 |
| C | 24.10700 | 34.76800 | 53.38500 |
| O | 23.80400 | 34.05500 | 54.34000 |
| C | 25.03010 | 33.79080 | 51.30370 |
| C | 24.78500 | 33.01300 | 50.00320 |
| C | 26.06970 | 32.50250 | 49.32810 |
| C | 26.91180 | 33.60970 | 48.68910 |
| N | 28.04330 | 33.00460 | 47.91700 |
| H | 23.49500 | 35.52110 | 50.27740 |
| H | 23.07740 | 33.42770 | 52.19990 |
| H | 25.67690 | 34.66580 | 51.11810 |
| H | 25.57670 | 33.14720 | 52.01060 |
| H | 24.14130 | 32.14870 | 50.23020 |
| H | 24.22530 | 33.62570 | 49.27720 |
| H | 26.68150 | 31.93950 | 50.05230 |
| H | 25.79380 | 31.78970 | 48.53580 |
| H | 26.30050 | 34.17800 | 47.97610 |
| H | 27.34330 | 34.29250 | 49.43280 |
| H | 28.69510 | 32.48690 | 48.55780 |
| H | 28.60970 | 33.69060 | 47.39510 |
| H | 27.66750 | 32.30760 | 47.21180 |
| N | 24.74100 | 35.92700 | 53.53200 |
| H | 24.85670 | 36.57410 | 52.75880 |
| O | 25.25940 | 33.44660 | 45.89610 |
| H | 25.45870 | 32.49290 | 45.96530 |
| H | 26.05630 | 33.77870 | 45.42600 |
| O | 27.96210 | 33.59820 | 44.94260 |
| H | 28.39660 | 33.78560 | 44.09350 |
| H | 27.86520 | 32.62040 | 44.97800 |
| H | 35.81610 | 29.27130 | 51.07370 |
| H | 33.84510 | 33.75030 | 50.00180 |
| H | 34.49810 | 31.23060 | 45.31330 |
| H | 35.20990 | 35.82760 | 44.14550 |
| H | 23.26610 | 27.63410 | 46.32760 |
| H | 26.53070 | 25.39980 | 43.50640 |
| H | 21.65590 | 36.76430 | 50.95360 |
| H | 24.88850 | 36.27960 | 54.47230 |

**Table S21.** Step 3 of the relaxed surface scan for proton transfer from complex 2 to complex 3 with explicit waters.

| | | | |
|---|---|---|---|
| C | 35.07600 | 29.61800 | 50.32400 |
| O | 35.29000 | 29.74100 | 49.12000 |
| N | 33.82600 | 29.84000 | 50.82400 |
| C | 32.80700 | 30.46300 | 50.04700 |
| C | 33.11400 | 31.92700 | 49.53200 |
| O | 32.91300 | 32.28500 | 48.36700 |
| C | 31.44600 | 30.40720 | 50.77360 |
| C | 30.29860 | 30.72790 | 49.82290 |
| O | 29.88030 | 31.88370 | 49.69170 |
| O | 29.84880 | 29.66970 | 49.18500 |
| H | 33.70500 | 29.79320 | 51.83460 |
| H | 32.71590 | 29.90460 | 49.10190 |
| H | 31.31180 | 29.38840 | 51.16110 |
| H | 31.42030 | 31.12240 | 51.60730 |
| H | 29.02150 | 29.85440 | 48.54490 |
| N | 33.63800 | 32.82200 | 50.35700 |
| H | 33.87670 | 32.61090 | 51.32130 |
| C | 33.90300 | 32.02700 | 44.81800 |
| O | 33.71700 | 32.08700 | 43.59100 |
| N | 33.39200 | 32.93300 | 45.68900 |
| C | 32.51000 | 34.06700 | 45.26000 |
| C | 33.32800 | 35.05900 | 44.40200 |

| | | | |
|---|---|---|---|
| O | 32.77200 | 35.51900 | 43.37500 |
| C | 31.80930 | 34.70210 | 46.48260 |
| C | 30.97200 | 35.90880 | 46.08570 |
| O | 30.91560 | 33.76060 | 47.09950 |
| H | 33.47750 | 32.73490 | 46.68910 |
| H | 31.74100 | 33.67470 | 44.58080 |
| H | 32.58670 | 35.00640 | 47.21020 |
| H | 31.45430 | 33.11530 | 47.61020 |
| H | 30.50060 | 36.33370 | 46.98190 |
| H | 30.18210 | 35.60110 | 45.38620 |
| H | 31.58130 | 36.68300 | 45.60270 |
| N | 34.61300 | 35.31200 | 44.78500 |
| H | 35.04980 | 34.71060 | 45.47620 |
| C | 23.94400 | 27.94100 | 45.50200 |
| O | 23.57900 | 28.09500 | 44.33100 |
| N | 25.20400 | 28.15800 | 45.87400 |
| C | 26.22300 | 28.52000 | 44.89200 |
| C | 26.47000 | 27.38700 | 43.85800 |
| O | 26.67800 | 27.71100 | 42.68900 |
| C | 27.55670 | 28.83580 | 45.59410 |
| C | 27.51590 | 30.05590 | 46.52030 |
| O | 27.14220 | 31.16360 | 46.02570 |
| O | 27.85340 | 29.90620 | 47.73780 |
| H | 25.47100 | 28.04030 | 46.84930 |
| H | 25.88040 | 29.37940 | 44.30320 |
| H | 27.90250 | 27.96070 | 46.16220 |
| H | 28.29750 | 29.04700 | 44.80810 |
| N | 26.43600 | 26.10900 | 44.22600 |
| H | 26.14880 | 25.82220 | 45.15600 |
| C | 22.02800 | 35.96100 | 51.62100 |
| O | 21.52500 | 35.67200 | 52.73400 |
| N | 23.04900 | 35.23800 | 51.14800 |
| C | 23.74700 | 34.28100 | 52.02300 |
| C | 24.10700 | 34.76800 | 53.38500 |
| O | 23.80400 | 34.05500 | 54.34000 |
| C | 25.03000 | 33.79070 | 51.30370 |
| C | 24.78460 | 33.01160 | 50.00430 |
| C | 26.06990 | 32.50230 | 49.32940 |
| C | 26.90840 | 33.60880 | 48.68550 |
| N | 28.03880 | 33.00210 | 47.91210 |
| H | 23.49470 | 35.52050 | 50.27700 |
| H | 23.07740 | 33.42770 | 52.20000 |
| H | 25.67660 | 34.66550 | 51.11710 |
| H | 25.57700 | 33.14780 | 52.01110 |
| H | 24.14210 | 32.14670 | 50.23210 |
| H | 24.22400 | 33.62290 | 49.27780 |
| H | 26.68380 | 31.94300 | 50.05480 |
| H | 25.79570 | 31.78620 | 48.53950 |
| H | 26.29480 | 34.17310 | 47.97120 |
| H | 27.34190 | 34.29670 | 49.42390 |
| H | 28.69020 | 32.48440 | 48.55160 |
| H | 28.60540 | 33.68740 | 47.38950 |
| H | 27.66120 | 32.30500 | 47.20670 |
| N | 24.74100 | 35.92700 | 53.53200 |
| H | 24.85670 | 36.57410 | 52.75880 |
| O | 25.25410 | 33.44310 | 45.89480 |
| H | 25.45620 | 32.48960 | 45.96260 |
| H | 26.05010 | 33.77730 | 45.42490 |
| O | 27.95670 | 33.59880 | 44.94150 |
| H | 28.39180 | 33.78550 | 44.09270 |
| H | 27.85790 | 32.62080 | 44.97710 |
| H | 35.81610 | 29.27120 | 51.07360 |
| H | 33.84470 | 33.75040 | 50.00170 |
| H | 34.49810 | 31.23060 | 45.31330 |
| H | 35.21000 | 35.82740 | 44.14540 |
| H | 23.26600 | 27.63450 | 46.32770 |
| H | 26.53060 | 25.39980 | 43.50650 |
| H | 21.65580 | 36.76420 | 50.95360 |
| H | 24.88850 | 36.27950 | 54.47230 |

**Table S22.** Step 2 of the relaxed surface scan for proton transfer from complex 2 to complex 3 with explicit waters.

| | | | |
|---|---|---|---|
| C | 35.07600 | 29.61800 | 50.32400 |
| O | 35.29000 | 29.74100 | 49.12000 |
| N | 33.82600 | 29.84000 | 50.82400 |
| C | 32.80700 | 30.46300 | 50.04700 |
| C | 33.11400 | 31.92700 | 49.53200 |
| O | 32.91300 | 32.28500 | 48.36700 |
| C | 31.45110 | 30.41320 | 50.78390 |
| C | 30.30130 | 30.72800 | 49.83640 |
| O | 29.88670 | 31.88220 | 49.68960 |
| O | 29.84650 | 29.66130 | 49.21280 |
| H | 33.70580 | 29.79500 | 51.83480 |
| H | 32.71120 | 29.90330 | 49.10330 |
| H | 31.31790 | 29.39730 | 51.17910 |
| H | 31.43220 | 31.13400 | 51.61270 |
| H | 29.03400 | 29.84600 | 48.57590 |
| N | 33.63800 | 32.82200 | 50.35700 |
| H | 33.87650 | 32.61120 | 51.32140 |
| C | 33.90300 | 32.02700 | 44.81800 |

| | | | |
|---|---|---|---|
| O | 33.71700 | 32.08700 | 43.59100 |
| N | 33.39200 | 32.93300 | 45.68900 |
| C | 32.51000 | 34.06700 | 45.26000 |
| C | 33.32800 | 35.05900 | 44.40200 |
| O | 32.77200 | 35.51900 | 43.37500 |
| C | 31.80710 | 34.70010 | 46.48200 |
| C | 30.96840 | 35.90590 | 46.08580 |
| O | 30.91380 | 33.75580 | 47.09540 |
| H | 33.47760 | 32.73490 | 46.68900 |
| H | 31.74160 | 33.67440 | 44.58020 |
| H | 32.58280 | 35.00390 | 47.21150 |
| H | 31.45260 | 33.11140 | 47.60740 |
| H | 30.49450 | 36.32840 | 46.98190 |
| H | 30.18020 | 35.59800 | 45.38450 |
| H | 31.57720 | 36.68190 | 45.60500 |
| N | 34.61300 | 35.31200 | 44.78500 |
| H | 35.04960 | 34.71100 | 45.47670 |
| C | 23.94400 | 27.94100 | 45.50200 |
| O | 23.57900 | 28.09500 | 44.33100 |
| N | 25.20400 | 28.15800 | 45.87400 |
| C | 26.22300 | 28.52000 | 44.89200 |
| C | 26.47000 | 27.38700 | 43.85800 |
| O | 26.67800 | 27.71100 | 42.68900 |
| C | 27.55530 | 28.83780 | 45.59270 |
| C | 27.50650 | 30.05620 | 46.52010 |
| O | 27.13540 | 31.16510 | 46.02320 |
| O | 27.83370 | 29.90280 | 47.73890 |
| H | 25.47100 | 28.04130 | 46.84950 |
| H | 25.87890 | 29.37840 | 44.30260 |
| H | 27.90340 | 27.96360 | 46.16070 |
| H | 28.29650 | 29.05190 | 44.80770 |
| N | 26.43600 | 26.10900 | 44.22600 |
| H | 26.14850 | 25.82230 | 45.15590 |
| C | 22.02800 | 35.96100 | 51.62100 |
| O | 21.52500 | 35.67200 | 52.73400 |
| N | 23.04900 | 35.23800 | 51.14800 |
| C | 23.74700 | 34.28100 | 52.02300 |
| C | 24.10700 | 34.76800 | 53.38500 |
| O | 23.80400 | 34.05500 | 54.34000 |
| C | 25.03200 | 33.79430 | 51.30520 |
| C | 24.79030 | 33.01450 | 50.00620 |
| C | 26.07730 | 32.50780 | 49.33440 |
| C | 26.91080 | 33.61100 | 48.67940 |
| N | 28.03860 | 32.99890 | 47.90680 |
| H | 23.49450 | 35.52030 | 50.27690 |
| H | 23.07880 | 33.42650 | 52.19910 |
| H | 25.67660 | 34.67090 | 51.11930 |
| H | 25.57990 | 33.15310 | 52.01350 |
| H | 24.14990 | 32.14780 | 50.23390 |
| H | 24.22860 | 33.62330 | 49.27850 |
| H | 26.69410 | 31.95410 | 50.06160 |
| H | 25.80440 | 31.78610 | 48.54900 |
| H | 26.29590 | 34.16960 | 47.96140 |
| H | 27.35070 | 34.30830 | 49.40220 |
| H | 28.68950 | 32.48150 | 48.54540 |
| H | 28.60590 | 33.68240 | 47.38300 |
| H | 27.65790 | 32.30120 | 47.20320 |
| N | 24.74100 | 35.92700 | 53.53200 |
| H | 24.85650 | 36.57420 | 52.75890 |
| O | 25.25470 | 33.43890 | 45.89710 |
| H | 25.45790 | 32.48560 | 45.96370 |
| H | 26.05000 | 33.77440 | 45.42650 |
| O | 27.95450 | 33.59640 | 44.94340 |
| H | 28.39040 | 33.78240 | 44.09480 |
| H | 27.85300 | 32.61840 | 44.97840 |
| H | 35.81600 | 29.27050 | 51.07340 |
| H | 33.84470 | 33.75030 | 50.00160 |
| H | 34.49810 | 31.23060 | 45.31330 |
| H | 35.21010 | 35.82680 | 44.14500 |
| H | 23.26590 | 27.63460 | 46.32770 |
| H | 26.53080 | 25.39970 | 43.50660 |
| H | 21.65600 | 36.76430 | 50.95370 |
| H | 24.88900 | 36.27930 | 54.47230 |

**Table S23.** Step 1 of the relaxed surface scan for proton transfer from complex 2 to complex 3 with explicit waters.

| | | | |
|---|---|---|---|
| C | 35.07600 | 29.61800 | 50.32400 |
| O | 35.29000 | 29.74100 | 49.12000 |
| N | 33.82600 | 29.84000 | 50.82400 |
| C | 32.80700 | 30.46300 | 50.04700 |
| C | 33.11400 | 31.92700 | 49.53200 |
| O | 32.91300 | 32.28500 | 48.36700 |
| C | 31.46520 | 30.44600 | 50.81140 |
| C | 30.30320 | 30.77930 | 49.88810 |
| O | 29.83990 | 31.92040 | 49.80990 |
| O | 29.89280 | 29.73020 | 49.20430 |
| H | 33.70130 | 29.78520 | 51.83350 |
| H | 32.69010 | 29.89950 | 49.10890 |
| H | 31.31480 | 29.43490 | 51.21390 |
| H | 31.47960 | 31.16990 | 51.63690 |

| | | | |
|---|---|---|---|
| H | 29.07560 | 29.90700 | 48.58700 |
| N | 33.63800 | 32.82200 | 50.35700 |
| H | 33.87760 | 32.61110 | 51.32110 |
| C | 33.90300 | 32.02700 | 44.81800 |
| O | 33.71700 | 32.08700 | 43.59100 |
| N | 33.39200 | 32.93300 | 45.68900 |
| C | 32.51000 | 34.06700 | 45.26000 |
| C | 33.32800 | 35.05900 | 44.40200 |
| O | 32.77200 | 35.51900 | 43.37500 |
| C | 31.81410 | 34.70660 | 46.48300 |
| C | 30.97520 | 35.91190 | 46.08670 |
| O | 30.92420 | 33.76720 | 47.10960 |
| H | 33.47950 | 32.73590 | 46.68890 |
| H | 31.73960 | 33.67520 | 44.58220 |
| H | 32.59560 | 35.01420 | 47.20510 |
| H | 31.47100 | 33.12320 | 47.61350 |
| H | 30.50950 | 36.34010 | 46.98420 |
| H | 30.18090 | 35.60260 | 45.39320 |
| H | 31.58220 | 36.68400 | 45.59760 |
| N | 34.61300 | 35.31200 | 44.78500 |
| H | 35.05030 | 34.71020 | 45.47560 |
| C | 23.94400 | 27.94100 | 45.50200 |
| O | 23.57900 | 28.09500 | 44.33100 |
| N | 25.20400 | 28.15800 | 45.87400 |
| C | 26.22300 | 28.52000 | 44.89200 |
| C | 26.47000 | 27.38700 | 43.85800 |
| O | 26.67800 | 27.71100 | 42.68900 |
| C | 27.55890 | 28.82400 | 45.58890 |
| C | 27.52750 | 30.01760 | 46.54900 |
| O | 27.22500 | 31.15750 | 46.06930 |
| O | 27.81280 | 29.80950 | 47.76840 |
| H | 25.47270 | 28.03770 | 46.84870 |
| H | 25.88040 | 29.37980 | 44.30350 |
| H | 27.90700 | 27.93550 | 46.13420 |
| H | 28.29390 | 29.05310 | 44.80240 |
| N | 26.43600 | 26.10900 | 44.22600 |
| H | 26.15000 | 25.82190 | 45.15630 |
| C | 22.02800 | 35.96100 | 51.62100 |
| O | 21.52500 | 35.67200 | 52.73400 |
| N | 23.04900 | 35.23800 | 51.14800 |
| C | 23.74700 | 34.28100 | 52.02300 |
| C | 24.10700 | 34.76800 | 53.38500 |
| O | 23.80400 | 34.05500 | 54.34000 |
| C | 25.02530 | 33.78520 | 51.30000 |
| C | 24.77110 | 32.99730 | 50.00600 |
| C | 26.05390 | 32.49060 | 49.32320 |
| C | 26.86270 | 33.59660 | 48.64210 |
| N | 28.04570 | 33.00550 | 47.93920 |
| H | 23.49930 | 35.52670 | 50.28140 |
| H | 23.07480 | 33.43000 | 52.20170 |
| H | 25.67210 | 34.65830 | 51.10480 |
| H | 25.57560 | 33.14600 | 52.00830 |
| H | 24.13490 | 32.13100 | 50.24630 |
| H | 24.19900 | 33.60210 | 49.28280 |
| H | 26.68810 | 31.96440 | 50.05650 |
| H | 25.78450 | 31.74740 | 48.55730 |
| H | 26.24930 | 34.09640 | 47.88000 |
| H | 27.23870 | 34.33980 | 49.35870 |
| H | 28.69360 | 32.54880 | 48.62340 |
| H | 28.59900 | 33.68990 | 47.40160 |
| H | 27.72750 | 32.25620 | 47.25070 |
| N | 24.74100 | 35.92700 | 53.53200 |
| H | 24.85730 | 36.57380 | 52.75870 |
| O | 25.29280 | 33.34540 | 45.78310 |
| H | 25.57750 | 32.41520 | 45.87750 |
| H | 26.07320 | 33.73950 | 45.33680 |
| O | 28.04520 | 33.57770 | 44.95180 |
| H | 28.52250 | 33.73430 | 44.11940 |
| H | 27.95100 | 32.60120 | 45.01960 |
| H | 35.81640 | 29.27160 | 51.07350 |
| H | 33.84010 | 33.75180 | 50.00270 |
| H | 34.49840 | 31.23080 | 45.31320 |
| H | 35.20970 | 35.82790 | 44.14550 |
| H | 23.26590 | 27.63480 | 46.32780 |
| H | 26.53380 | 25.39970 | 43.50700 |
| H | 21.65580 | 36.76420 | 50.95360 |
| H | 24.88740 | 36.28030 | 54.47220 |

**Table S24.** Step 8 of the relaxed surface scan for proton transfer from complex 2 to complex 3 without explicit waters.

| | | | |
|---|---|---|---|
| C | 35.07600 | 29.61800 | 50.32400 |
| O | 35.29000 | 29.74100 | 49.12000 |
| N | 33.82600 | 29.84000 | 50.82400 |
| C | 32.80700 | 30.46300 | 50.04700 |
| C | 33.11400 | 31.92700 | 49.53200 |
| O | 32.91300 | 32.28500 | 48.36700 |
| C | 31.48460 | 30.49620 | 50.83720 |
| C | 30.29740 | 30.92260 | 49.96630 |
| O | 29.62900 | 31.93650 | 50.29930 |
| O | 30.08810 | 30.18350 | 48.94610 |

| | | | |
|---|---|---|---|
| H | 33.70050 | 29.78780 | 51.83340 |
| H | 32.65410 | 29.89670 | 49.11750 |
| H | 31.27670 | 29.47630 | 51.19610 |
| H | 31.56560 | 31.16760 | 51.70370 |
| H | 28.90100 | 29.92610 | 48.34590 |
| N | 33.63800 | 32.82200 | 50.35700 |
| H | 33.86840 | 32.61160 | 51.32340 |
| C | 33.90300 | 32.02700 | 44.81800 |
| O | 33.71700 | 32.08700 | 43.59100 |
| N | 33.39200 | 32.93300 | 45.68900 |
| C | 32.51000 | 34.06700 | 45.26000 |
| C | 33.32800 | 35.05900 | 44.40200 |
| O | 32.77200 | 35.51900 | 43.37500 |
| C | 31.82690 | 34.71310 | 46.48660 |
| C | 30.99620 | 35.92500 | 46.09410 |
| O | 30.93810 | 33.77580 | 47.12450 |
| H | 33.47400 | 32.72980 | 46.68860 |
| H | 31.73620 | 33.67750 | 44.58500 |
| H | 32.61330 | 35.01500 | 47.20490 |
| H | 31.49240 | 33.12900 | 47.62100 |
| H | 30.51240 | 36.33940 | 46.98880 |
| H | 30.22120 | 35.63610 | 45.37020 |
| H | 31.61760 | 36.70400 | 45.63480 |
| N | 34.61300 | 35.31200 | 44.78500 |
| H | 35.05070 | 34.70560 | 45.47150 |
| C | 23.94400 | 27.94100 | 45.50200 |
| O | 23.57900 | 28.09500 | 44.33100 |
| N | 25.20400 | 28.15800 | 45.87400 |
| C | 26.22300 | 28.52000 | 44.89200 |
| C | 26.47000 | 27.38700 | 43.85800 |
| O | 26.67800 | 27.71100 | 42.68900 |
| C | 27.58940 | 28.71530 | 45.57010 |
| C | 27.74460 | 29.80340 | 46.62220 |
| O | 27.56830 | 31.00260 | 46.35800 |
| O | 28.14340 | 29.32980 | 47.77990 |
| H | 25.47230 | 28.02430 | 46.84680 |
| H | 25.90710 | 29.39840 | 44.31830 |
| H | 27.90990 | 27.76440 | 46.01610 |
| H | 28.30890 | 28.96620 | 44.77500 |
| N | 26.43600 | 26.10900 | 44.22600 |
| H | 26.16100 | 25.81720 | 45.15800 |
| C | 22.02800 | 35.96100 | 51.62100 |
| O | 21.52500 | 35.67200 | 52.73400 |
| N | 23.04900 | 35.23800 | 51.14800 |
| C | 23.74700 | 34.28100 | 52.02300 |
| C | 24.10700 | 34.76800 | 53.38500 |
| O | 23.80400 | 34.05500 | 54.34000 |
| C | 25.03350 | 33.79120 | 51.31040 |
| C | 24.80670 | 33.03090 | 49.99570 |
| C | 26.10370 | 32.48740 | 49.36850 |
| C | 27.00220 | 33.56570 | 48.75400 |
| N | 28.24930 | 32.93580 | 48.22000 |
| H | 23.50160 | 35.53050 | 50.28410 |
| H | 23.07720 | 33.42780 | 52.19920 |
| H | 25.68960 | 34.66330 | 51.14690 |
| H | 25.56540 | 33.13430 | 52.01630 |
| H | 24.13270 | 32.18290 | 50.19330 |
| H | 24.28720 | 33.66570 | 49.25820 |
| H | 26.67810 | 31.93390 | 50.12980 |
| H | 25.84780 | 31.76350 | 48.57890 |
| H | 26.49050 | 34.08010 | 47.92870 |
| H | 27.30390 | 34.31710 | 49.49610 |
| H | 28.78410 | 32.47870 | 49.03570 |
| H | 28.90070 | 33.59700 | 47.76570 |
| H | 28.00870 | 32.18620 | 47.51150 |
| N | 24.74100 | 35.92700 | 53.53200 |
| H | 24.86230 | 36.57340 | 52.75920 |
| H | 35.81770 | 29.27180 | 51.07290 |
| H | 33.83590 | 33.75370 | 50.00580 |
| H | 34.49680 | 31.23010 | 45.31380 |
| H | 35.20980 | 35.82470 | 44.14290 |
| H | 23.26690 | 27.63170 | 46.32740 |
| H | 26.55280 | 25.40060 | 43.50910 |
| H | 21.65660 | 36.76530 | 50.95430 |
| H | 24.89450 | 36.27670 | 54.47240 |

**Table S25.** Step 7 of the relaxed surface scan for proton transfer from complex 2 to complex 3 without explicit waters.

| | | | |
|---|---|---|---|
| C | 35.07600 | 29.61800 | 50.32400 |
| O | 35.29000 | 29.74100 | 49.12000 |
| N | 33.82600 | 29.84000 | 50.82400 |
| C | 32.80700 | 30.46300 | 50.04700 |
| C | 33.11400 | 31.92700 | 49.53200 |
| O | 32.91300 | 32.28500 | 48.36700 |
| C | 31.43500 | 30.40890 | 50.74680 |
| C | 30.28820 | 30.72500 | 49.77380 |
| O | 29.75220 | 31.85530 | 49.82280 |
| O | 29.97610 | 29.76540 | 48.97060 |
| H | 33.70370 | 29.79140 | 51.83430 |
| H | 32.72220 | 29.90620 | 49.10040 |

| | | | |
|---|---|---|---|
| H | 31.29180 | 29.38590 | 51.12320 |
| H | 31.39660 | 31.11360 | 51.58880 |
| H | 28.92560 | 29.86080 | 48.31450 |
| N | 33.63800 | 32.82200 | 50.35700 |
| H | 33.87330 | 32.60950 | 51.32180 |
| C | 33.90300 | 32.02700 | 44.81800 |
| O | 33.71700 | 32.08700 | 43.59100 |
| N | 33.39200 | 32.93300 | 45.68900 |
| C | 32.51000 | 34.06700 | 45.26000 |
| C | 33.32800 | 35.05900 | 44.40200 |
| O | 32.77200 | 35.51900 | 43.37500 |
| C | 31.82200 | 34.70880 | 46.48550 |
| C | 31.01650 | 35.93940 | 46.09760 |
| O | 30.90790 | 33.78030 | 47.09860 |
| H | 33.47360 | 32.73190 | 46.68920 |
| H | 31.73740 | 33.67780 | 44.58330 |
| H | 32.60340 | 34.98890 | 47.21790 |
| H | 31.43810 | 33.11750 | 47.59740 |
| H | 30.52110 | 36.34520 | 46.98980 |
| H | 30.25230 | 35.67510 | 45.35320 |
| H | 31.65800 | 36.71710 | 45.66440 |
| N | 34.61300 | 35.31200 | 44.78500 |
| H | 35.05060 | 34.70590 | 45.47180 |
| C | 23.94400 | 27.94100 | 45.50200 |
| O | 23.57900 | 28.09500 | 44.33100 |
| N | 25.20400 | 28.15800 | 45.87400 |
| C | 26.22300 | 28.52000 | 44.89200 |
| C | 26.47000 | 27.38700 | 43.85800 |
| O | 26.67800 | 27.71100 | 42.68900 |
| C | 27.56090 | 28.81000 | 45.60630 |
| C | 27.55000 | 30.01740 | 46.54830 |
| O | 27.21040 | 31.13340 | 46.11170 |
| O | 27.88160 | 29.80270 | 47.79390 |
| H | 25.47200 | 28.03040 | 46.84780 |
| H | 25.88960 | 29.38570 | 44.30780 |
| H | 27.89090 | 27.92030 | 46.16040 |
| H | 28.31020 | 29.02130 | 44.82820 |
| N | 26.43600 | 26.10900 | 44.22600 |
| H | 26.16160 | 25.82020 | 45.15910 |
| C | 22.02800 | 35.96100 | 51.62100 |
| O | 21.52500 | 35.67200 | 52.73400 |
| N | 23.04900 | 35.23800 | 51.14800 |
| C | 23.74700 | 34.28100 | 52.02300 |
| C | 24.10700 | 34.76800 | 53.38500 |
| O | 23.80400 | 34.05500 | 54.34000 |
| C | 25.02890 | 33.78780 | 51.30290 |
| C | 24.78240 | 33.01640 | 49.99820 |
| C | 26.06390 | 32.48660 | 49.32980 |
| C | 26.93990 | 33.57880 | 48.70860 |
| N | 28.09690 | 32.95900 | 47.99030 |
| H | 23.50200 | 35.53080 | 50.28440 |
| H | 23.07610 | 33.42880 | 52.20020 |
| H | 25.68060 | 34.66030 | 51.12300 |
| H | 25.57100 | 33.13820 | 52.00760 |
| H | 24.12570 | 32.16020 | 50.21840 |
| H | 24.23240 | 33.63980 | 49.27310 |
| H | 26.66110 | 31.91590 | 50.05960 |
| H | 25.78630 | 31.77750 | 48.53500 |
| H | 26.36710 | 34.17210 | 47.98170 |
| H | 27.34640 | 34.26070 | 49.46750 |
| H | 28.69750 | 32.40610 | 48.67710 |
| H | 28.73050 | 33.64430 | 47.55090 |
| H | 27.74760 | 32.29280 | 47.24020 |
| N | 24.74100 | 35.92700 | 53.53200 |
| H | 24.86020 | 36.57370 | 52.75910 |
| H | 35.81620 | 29.26970 | 51.07310 |
| H | 33.84530 | 33.75070 | 50.00320 |
| H | 34.49740 | 31.23040 | 45.31360 |
| H | 35.21010 | 35.82380 | 44.14260 |
| H | 23.26630 | 27.63290 | 46.32740 |
| H | 26.54020 | 25.39960 | 43.50810 |
| H | 21.65650 | 36.76520 | 50.95430 |
| H | 24.89210 | 36.27790 | 54.47240 |

**Table S26.** Step 6 of the relaxed surface scan for proton transfer from complex 2 to complex 3 without explicit waters.

| | | | |
|---|---|---|---|
| C | 35.07600 | 29.61800 | 50.32400 |
| O | 35.29000 | 29.74100 | 49.12000 |
| N | 33.82600 | 29.84000 | 50.82400 |
| C | 32.80700 | 30.46300 | 50.04700 |
| C | 33.11400 | 31.92700 | 49.53200 |
| O | 32.91300 | 32.28500 | 48.36700 |
| C | 31.43370 | 30.40030 | 50.74340 |
| C | 30.28880 | 30.71410 | 49.77260 |
| O | 29.75750 | 31.84220 | 49.79650 |
| O | 29.96900 | 29.73510 | 48.98410 |
| H | 33.70490 | 29.79290 | 51.83460 |
| H | 32.72630 | 29.90710 | 49.09920 |
| H | 31.29640 | 29.37610 | 51.11800 |
| H | 31.38680 | 31.10480 | 51.58520 |

| | | | |
|---|---|---|---|
| H | 28.98250 | 29.85250 | 48.34640 |
| N | 33.63800 | 32.82200 | 50.35700 |
| H | 33.87540 | 32.60930 | 51.32120 |
| C | 33.90300 | 32.02700 | 44.81800 |
| O | 33.71700 | 32.08700 | 43.59100 |
| N | 33.39200 | 32.93300 | 45.68900 |
| C | 32.51000 | 34.06700 | 45.26000 |
| C | 33.32800 | 35.05900 | 44.40200 |
| O | 32.77200 | 35.51900 | 43.37500 |
| C | 31.81810 | 34.70610 | 46.48420 |
| C | 31.01480 | 35.93830 | 46.09650 |
| O | 30.90060 | 33.77550 | 47.08960 |
| H | 33.47390 | 32.73230 | 46.68920 |
| H | 31.73870 | 33.67730 | 44.58210 |
| H | 32.59610 | 34.98360 | 47.22110 |
| H | 31.42840 | 33.11250 | 47.58970 |
| H | 30.51530 | 36.34120 | 46.98770 |
| H | 30.25410 | 35.67670 | 45.34750 |
| H | 31.65880 | 36.71720 | 45.66910 |
| N | 34.61300 | 35.31200 | 44.78500 |
| H | 35.05060 | 34.70570 | 45.47160 |
| C | 23.94400 | 27.94100 | 45.50200 |
| O | 23.57900 | 28.09500 | 44.33100 |
| N | 25.20400 | 28.15800 | 45.87400 |
| C | 26.22300 | 28.52000 | 44.89200 |
| C | 26.47000 | 27.38700 | 43.85800 |
| O | 26.67800 | 27.71100 | 42.68900 |
| C | 27.55910 | 28.81460 | 45.60780 |
| C | 27.54220 | 30.02320 | 46.55250 |
| O | 27.17200 | 31.13240 | 46.10840 |
| O | 27.90270 | 29.82690 | 47.78340 |
| H | 25.47190 | 28.03380 | 46.84820 |
| H | 25.88930 | 29.38570 | 44.30810 |
| H | 27.89220 | 27.92550 | 46.16160 |
| H | 28.30760 | 29.02870 | 44.82960 |
| N | 26.43600 | 26.10900 | 44.22600 |
| H | 26.16150 | 25.82090 | 45.15930 |
| C | 22.02800 | 35.96100 | 51.62100 |
| O | 21.52500 | 35.67200 | 52.73400 |
| N | 23.04900 | 35.23800 | 51.14800 |
| C | 23.74700 | 34.28100 | 52.02300 |
| C | 24.10700 | 34.76800 | 53.38500 |
| O | 23.80400 | 34.05500 | 54.34000 |
| C | 25.02570 | 33.78380 | 51.30000 |
| C | 24.76990 | 33.00920 | 49.99860 |
| C | 26.04500 | 32.47700 | 49.32010 |
| C | 26.91580 | 33.56840 | 48.69060 |
| N | 28.06620 | 32.94950 | 47.96090 |
| H | 23.50140 | 35.53020 | 50.28390 |
| H | 23.07480 | 33.43000 | 52.20150 |
| H | 25.67810 | 34.65460 | 51.11490 |
| H | 25.56940 | 33.13470 | 52.00390 |
| H | 24.11440 | 32.15400 | 50.22620 |
| H | 24.21450 | 33.63100 | 49.27630 |
| H | 26.64780 | 31.90610 | 50.04510 |
| H | 25.76070 | 31.76760 | 48.52790 |
| H | 26.33680 | 34.16100 | 47.96810 |
| H | 27.32840 | 34.25110 | 49.44560 |
| H | 28.66730 | 32.39590 | 48.63710 |
| H | 28.69740 | 33.63730 | 47.52270 |
| H | 27.71030 | 32.28030 | 47.20840 |
| N | 24.74100 | 35.92700 | 53.53200 |
| H | 24.85990 | 36.57370 | 52.75900 |
| H | 35.81600 | 29.27010 | 51.07340 |
| H | 33.84720 | 33.75000 | 50.00240 |
| H | 34.49750 | 31.23040 | 45.31360 |
| H | 35.21010 | 35.82370 | 44.14240 |
| H | 23.26600 | 27.63330 | 46.32750 |
| H | 26.53910 | 25.39930 | 43.50820 |
| H | 21.65660 | 36.76520 | 50.95430 |
| H | 24.89140 | 36.27830 | 54.47240 |

**Table S27.** Step 5 of the relaxed surface scan for proton transfer from complex 2 to complex 3 without explicit waters.

| | | | |
|---|---|---|---|
| C | 35.07600 | 29.61800 | 50.32400 |
| O | 35.29000 | 29.74100 | 49.12000 |
| N | 33.82600 | 29.84000 | 50.82400 |
| C | 32.80700 | 30.46300 | 50.04700 |
| C | 33.11400 | 31.92700 | 49.53200 |
| O | 32.91300 | 32.28500 | 48.36700 |
| C | 31.43930 | 30.40570 | 50.75590 |
| C | 30.28920 | 30.70260 | 49.79060 |
| O | 29.76590 | 31.83070 | 49.77120 |
| O | 29.95540 | 29.69030 | 49.04240 |
| H | 33.70460 | 29.79210 | 51.83450 |
| H | 32.72120 | 29.90570 | 49.10070 |
| H | 31.30730 | 29.38690 | 51.14630 |
| H | 31.39810 | 31.12240 | 51.58760 |
| H | 29.02700 | 29.83040 | 48.39890 |
| N | 33.63800 | 32.82200 | 50.35700 |

| | | | |
|---|---|---|---|
| H | 33.87730 | 32.60920 | 51.32080 |
| C | 33.90300 | 32.02700 | 44.81800 |
| O | 33.71700 | 32.08700 | 43.59100 |
| N | 33.39200 | 32.93300 | 45.68900 |
| C | 32.51000 | 34.06700 | 45.26000 |
| C | 33.32800 | 35.05900 | 44.40200 |
| O | 32.77200 | 35.51900 | 43.37500 |
| C | 31.81590 | 34.70420 | 46.48380 |
| C | 31.01430 | 35.93760 | 46.09670 |
| O | 30.89640 | 33.77290 | 47.08550 |
| H | 33.47510 | 32.73310 | 46.68920 |
| H | 31.73950 | 33.67690 | 44.58140 |
| H | 32.59250 | 34.97980 | 47.22290 |
| H | 31.42260 | 33.10900 | 47.58560 |
| H | 30.51240 | 36.33860 | 46.98740 |
| H | 30.25570 | 35.67840 | 45.34480 |
| H | 31.66010 | 36.71720 | 45.67310 |
| N | 34.61300 | 35.31200 | 44.78500 |
| H | 35.05070 | 34.70540 | 45.47130 |
| C | 23.94400 | 27.94100 | 45.50200 |
| O | 23.57900 | 28.09500 | 44.33100 |
| N | 25.20400 | 28.15800 | 45.87400 |
| C | 26.22300 | 28.52000 | 44.89200 |
| C | 26.47000 | 27.38700 | 43.85800 |
| O | 26.67800 | 27.71100 | 42.68900 |
| C | 27.55550 | 28.82140 | 45.60730 |
| C | 27.53320 | 30.03350 | 46.54840 |
| O | 27.12820 | 31.13510 | 46.10280 |
| O | 27.93080 | 29.85740 | 47.76210 |
| H | 25.47140 | 28.03610 | 46.84850 |
| H | 25.88690 | 29.38410 | 44.30710 |
| H | 27.89240 | 27.93570 | 46.16460 |
| H | 28.30540 | 29.03570 | 44.83040 |
| N | 26.43600 | 26.10900 | 44.22600 |
| H | 26.16060 | 25.82140 | 45.15920 |
| C | 22.02800 | 35.96100 | 51.62100 |
| O | 21.52500 | 35.67200 | 52.73400 |
| N | 23.04900 | 35.23800 | 51.14800 |
| C | 23.74700 | 34.28100 | 52.02300 |
| C | 24.10700 | 34.76800 | 53.38500 |
| O | 23.80400 | 34.05500 | 54.34000 |
| C | 25.02430 | 33.78130 | 51.29920 |
| C | 24.76430 | 32.99960 | 50.00280 |
| C | 26.03700 | 32.46800 | 49.31890 |
| C | 26.89590 | 33.55850 | 48.67190 |
| N | 28.04060 | 32.94050 | 47.93290 |
| H | 23.50240 | 35.53130 | 50.28470 |
| H | 23.07390 | 33.43080 | 52.20210 |
| H | 25.67580 | 34.65150 | 51.10770 |
| H | 25.57040 | 33.13610 | 52.00480 |
| H | 24.11270 | 32.14370 | 50.23830 |
| H | 24.20230 | 33.61640 | 49.28130 |
| H | 26.64930 | 31.90740 | 50.04420 |
| H | 25.75040 | 31.75010 | 48.53540 |
| H | 26.30620 | 34.14110 | 47.94990 |
| H | 27.31140 | 34.25070 | 49.41660 |
| H | 28.63850 | 32.37810 | 48.59760 |
| H | 28.67370 | 33.63090 | 47.50210 |
| H | 27.67690 | 32.27140 | 47.17570 |
| N | 24.74100 | 35.92700 | 53.53200 |
| H | 24.85930 | 36.57370 | 52.75900 |
| H | 35.81610 | 29.27020 | 51.07340 |
| H | 33.84890 | 33.74930 | 50.00150 |
| H | 34.49750 | 31.23030 | 45.31350 |
| H | 35.21020 | 35.82320 | 44.14220 |
| H | 23.26590 | 27.63350 | 46.32750 |
| H | 26.53800 | 25.39920 | 43.50820 |
| H | 21.65650 | 36.76510 | 50.95430 |
| H | 24.89060 | 36.27880 | 54.47230 |

**Table S28.** Step 4 of the relaxed surface scan for proton transfer from complex 2 to complex 3 without explicit waters.

| | | | |
|---|---|---|---|
| C | 35.07600 | 29.61800 | 50.32400 |
| O | 35.29000 | 29.74100 | 49.12000 |
| N | 33.82600 | 29.84000 | 50.82400 |
| C | 32.80700 | 30.46300 | 50.04700 |
| C | 33.11400 | 31.92700 | 49.53200 |
| O | 32.91300 | 32.28500 | 48.36700 |
| C | 31.44240 | 30.40850 | 50.76290 |
| C | 30.28950 | 30.70150 | 49.80300 |
| O | 29.76380 | 31.82540 | 49.77340 |
| O | 29.95520 | 29.67820 | 49.06280 |
| H | 33.70470 | 29.79220 | 51.83450 |
| H | 32.71810 | 29.90500 | 49.10160 |
| H | 31.31200 | 29.39140 | 51.15840 |
| H | 31.40480 | 31.12900 | 51.59130 |
| H | 29.05630 | 29.82300 | 48.43040 |
| N | 33.63800 | 32.82200 | 50.35700 |
| H | 33.87840 | 32.60920 | 51.32050 |
| C | 33.90300 | 32.02700 | 44.81800 |

| | | | |
|---|---|---|---|
| O | 33.71700 | 32.08700 | 43.59100 |
| N | 33.39200 | 32.93300 | 45.68900 |
| C | 32.51000 | 34.06700 | 45.26000 |
| C | 33.32800 | 35.05900 | 44.40200 |
| O | 32.77200 | 35.51900 | 43.37500 |
| C | 31.81500 | 34.70390 | 46.48330 |
| C | 31.01350 | 35.93730 | 46.09580 |
| O | 30.89520 | 33.77210 | 47.08380 |
| H | 33.47560 | 32.73330 | 46.68910 |
| H | 31.73990 | 33.67680 | 44.58110 |
| H | 32.59090 | 34.97950 | 47.22320 |
| H | 31.42140 | 33.10840 | 47.58390 |
| H | 30.51070 | 36.33790 | 46.98620 |
| H | 30.25560 | 35.67810 | 45.34320 |
| H | 31.65950 | 36.71710 | 45.67300 |
| N | 34.61300 | 35.31200 | 44.78500 |
| H | 35.05080 | 34.70530 | 45.47110 |
| C | 23.94400 | 27.94100 | 45.50200 |
| O | 23.57900 | 28.09500 | 44.33100 |
| N | 25.20400 | 28.15800 | 45.87400 |
| C | 26.22300 | 28.52000 | 44.89200 |
| C | 26.47000 | 27.38700 | 43.85800 |
| O | 26.67800 | 27.71100 | 42.68900 |
| C | 27.55410 | 28.82590 | 45.60630 |
| C | 27.52480 | 30.03630 | 46.55100 |
| O | 27.11710 | 31.13820 | 46.10130 |
| O | 27.92130 | 29.86120 | 47.76160 |
| H | 25.47130 | 28.03850 | 46.84890 |
| H | 25.88520 | 29.38310 | 44.30650 |
| H | 27.89540 | 27.94090 | 46.16210 |
| H | 28.30200 | 29.04510 | 44.82880 |
| N | 26.43600 | 26.10900 | 44.22600 |
| H | 26.16030 | 25.82180 | 45.15920 |
| C | 22.02800 | 35.96100 | 51.62100 |
| O | 21.52500 | 35.67200 | 52.73400 |
| N | 23.04900 | 35.23800 | 51.14800 |
| C | 23.74700 | 34.28100 | 52.02300 |
| C | 24.10700 | 34.76800 | 53.38500 |
| O | 23.80400 | 34.05500 | 54.34000 |
| C | 25.02260 | 33.77930 | 51.29780 |
| C | 24.75870 | 32.99530 | 50.00330 |
| C | 26.02880 | 32.46210 | 49.31600 |
| C | 26.88430 | 33.55150 | 48.66260 |
| N | 28.02890 | 32.93400 | 47.92360 |
| H | 23.50260 | 35.53150 | 50.28490 |
| H | 23.07320 | 33.43140 | 52.20270 |
| H | 25.67450 | 34.64850 | 51.10340 |
| H | 25.56970 | 33.13470 | 52.00340 |
| H | 24.10750 | 32.14000 | 50.24260 |
| H | 24.19430 | 33.61090 | 49.28270 |
| H | 26.64430 | 31.90390 | 50.04040 |
| H | 25.74030 | 31.74150 | 48.53570 |
| H | 26.29190 | 34.12970 | 47.93940 |
| H | 27.29910 | 34.24790 | 49.40390 |
| H | 28.62300 | 32.36940 | 48.58570 |
| H | 28.66360 | 33.62490 | 47.49660 |
| H | 27.66660 | 32.26290 | 47.16260 |
| N | 24.74100 | 35.92700 | 53.53200 |
| H | 24.85920 | 36.57370 | 52.75890 |
| H | 35.81610 | 29.27040 | 51.07340 |
| H | 33.84910 | 33.74910 | 50.00110 |
| H | 34.49750 | 31.23030 | 45.31350 |
| H | 35.21020 | 35.82310 | 44.14210 |
| H | 23.26570 | 27.63390 | 46.32750 |
| H | 26.53750 | 25.39910 | 43.50820 |
| H | 21.65640 | 36.76510 | 50.95420 |
| H | 24.89030 | 36.27900 | 54.47230 |

**Table S29.** Step 3 of the relaxed surface scan for proton transfer from complex 2 to complex 3 without explicit waters.

| | | | |
|---|---|---|---|
| C | 35.07600 | 29.61800 | 50.32400 |
| O | 35.29000 | 29.74100 | 49.12000 |
| N | 33.82600 | 29.84000 | 50.82400 |
| C | 32.80700 | 30.46300 | 50.04700 |
| C | 33.11400 | 31.92700 | 49.53200 |
| O | 32.91300 | 32.28500 | 48.36700 |
| C | 31.44590 | 30.41170 | 50.77040 |
| C | 30.28980 | 30.70050 | 49.81620 |
| O | 29.76220 | 31.82090 | 49.77640 |
| O | 29.95420 | 29.66670 | 49.08480 |
| H | 33.70460 | 29.79200 | 51.83450 |
| H | 32.71470 | 29.90410 | 49.10250 |
| H | 31.31720 | 29.39660 | 51.17140 |
| H | 31.41240 | 31.13640 | 51.59520 |
| H | 29.07790 | 29.81520 | 48.46160 |
| N | 33.63800 | 32.82200 | 50.35700 |
| H | 33.87940 | 32.60920 | 51.32030 |
| C | 33.90300 | 32.02700 | 44.81800 |
| O | 33.71700 | 32.08700 | 43.59100 |
| N | 33.39200 | 32.93300 | 45.68900 |

| | | | |
|---|---|---|---|
| C | 32.51000 | 34.06700 | 45.26000 |
| C | 33.32800 | 35.05900 | 44.40200 |
| O | 32.77200 | 35.51900 | 43.37500 |
| C | 31.81440 | 34.70390 | 46.48290 |
| C | 31.01300 | 35.93720 | 46.09490 |
| O | 30.89410 | 33.77220 | 47.08290 |
| H | 33.47610 | 32.73350 | 46.68900 |
| H | 31.74000 | 33.67660 | 44.58100 |
| H | 32.58990 | 34.97960 | 47.22320 |
| H | 31.42050 | 33.10850 | 47.58280 |
| H | 30.50970 | 36.33790 | 46.98490 |
| H | 30.25550 | 35.67780 | 45.34200 |
| H | 31.65910 | 36.71700 | 45.67230 |
| N | 34.61300 | 35.31200 | 44.78500 |
| H | 35.05090 | 34.70520 | 45.47100 |
| C | 23.94400 | 27.94100 | 45.50200 |
| O | 23.57900 | 28.09500 | 44.33100 |
| N | 25.20400 | 28.15800 | 45.87400 |
| C | 26.22300 | 28.52000 | 44.89200 |
| C | 26.47000 | 27.38700 | 43.85800 |
| O | 26.67800 | 27.71100 | 42.68900 |
| C | 27.55290 | 28.83050 | 45.60500 |
| C | 27.51620 | 30.04020 | 46.55130 |
| O | 27.11240 | 31.14340 | 46.09500 |
| O | 27.90350 | 29.86550 | 47.76180 |
| H | 25.47120 | 28.04070 | 46.84920 |
| H | 25.88350 | 29.38210 | 44.30580 |
| H | 27.89770 | 27.94680 | 46.16070 |
| H | 28.29890 | 29.05320 | 44.82690 |
| N | 26.43600 | 26.10900 | 44.22600 |
| H | 26.16000 | 25.82210 | 45.15920 |
| C | 22.02800 | 35.96100 | 51.62100 |
| O | 21.52500 | 35.67200 | 52.73400 |
| N | 23.04900 | 35.23800 | 51.14800 |
| C | 23.74700 | 34.28100 | 52.02300 |
| C | 24.10700 | 34.76800 | 53.38500 |
| O | 23.80400 | 34.05500 | 54.34000 |
| C | 25.02140 | 33.77800 | 51.29650 |
| C | 24.75440 | 32.99400 | 50.00240 |
| C | 26.02230 | 32.45900 | 49.31230 |
| C | 26.87660 | 33.54700 | 48.65510 |
| N | 28.02040 | 32.92880 | 47.91540 |
| H | 23.50270 | 35.53160 | 50.28500 |
| H | 23.07280 | 33.43180 | 52.20300 |
| H | 25.67400 | 34.64650 | 51.10100 |
| H | 25.56870 | 33.13300 | 52.00140 |
| H | 24.10280 | 32.13960 | 50.24320 |
| H | 24.18920 | 33.61000 | 49.28270 |
| H | 26.63940 | 31.90140 | 50.03570 |
| H | 25.73170 | 31.73730 | 48.53380 |
| H | 26.28300 | 34.12320 | 47.93120 |
| H | 27.29180 | 34.24550 | 49.39430 |
| H | 28.60960 | 32.36010 | 48.57520 |
| H | 28.65810 | 33.62000 | 47.49350 |
| H | 27.65760 | 32.25840 | 47.14990 |
| N | 24.74100 | 35.92700 | 53.53200 |
| H | 24.85900 | 36.57370 | 52.75890 |
| H | 35.81610 | 29.27040 | 51.07340 |
| H | 33.84920 | 33.74900 | 50.00090 |
| H | 34.49760 | 31.23040 | 45.31350 |
| H | 35.21020 | 35.82300 | 44.14200 |
| H | 23.26560 | 27.63420 | 46.32760 |
| H | 26.53680 | 25.39900 | 43.50820 |
| H | 21.65650 | 36.76510 | 50.95430 |
| H | 24.88990 | 36.27920 | 54.47230 |

**Table S30.** Step 2 of the relaxed surface scan for proton transfer from complex 2 to complex 3 without explicit waters.

| | | | |
|---|---|---|---|
| C | 35.07600 | 29.61800 | 50.32400 |
| O | 35.29000 | 29.74100 | 49.12000 |
| N | 33.82600 | 29.84000 | 50.82400 |
| C | 32.80700 | 30.46300 | 50.04700 |
| C | 33.11400 | 31.92700 | 49.53200 |
| O | 32.91300 | 32.28500 | 48.36700 |
| C | 31.44980 | 30.41670 | 50.77910 |
| C | 30.28980 | 30.70160 | 49.83140 |
| O | 29.76000 | 31.81810 | 49.78250 |
| O | 29.95150 | 29.65740 | 49.10890 |
| H | 33.70460 | 29.79190 | 51.83440 |
| H | 32.71020 | 29.90300 | 49.10380 |
| H | 31.32180 | 29.40420 | 51.18690 |
| H | 31.42170 | 31.14640 | 51.59940 |
| H | 29.09340 | 29.80900 | 48.49320 |
| N | 33.63800 | 32.82200 | 50.35700 |
| H | 33.88030 | 32.60920 | 51.32000 |
| C | 33.90300 | 32.02700 | 44.81800 |
| O | 33.71700 | 32.08700 | 43.59100 |
| N | 33.39200 | 32.93300 | 45.68900 |
| C | 32.51000 | 34.06700 | 45.26000 |
| C | 33.32800 | 35.05900 | 44.40200 |

| | | | |
|---|---|---|---|
| O | 32.77200 | 35.51900 | 43.37500 |
| C | 31.81410 | 34.70430 | 46.48270 |
| C | 31.01260 | 35.93720 | 46.09400 |
| O | 30.89380 | 33.77260 | 47.08260 |
| H | 33.47670 | 32.73360 | 46.68900 |
| H | 31.73970 | 33.67660 | 44.58140 |
| H | 32.58940 | 34.98010 | 47.22320 |
| H | 31.42020 | 33.10890 | 47.58220 |
| H | 30.50910 | 36.33840 | 46.98360 |
| H | 30.25540 | 35.67710 | 45.34110 |
| H | 31.65860 | 36.71700 | 45.67120 |
| N | 34.61300 | 35.31200 | 44.78500 |
| H | 35.05100 | 34.70490 | 45.47070 |
| C | 23.94400 | 27.94100 | 45.50200 |
| O | 23.57900 | 28.09500 | 44.33100 |
| N | 25.20400 | 28.15800 | 45.87400 |
| C | 26.22300 | 28.52000 | 44.89200 |
| C | 26.47000 | 27.38700 | 43.85800 |
| O | 26.67800 | 27.71100 | 42.68900 |
| C | 27.55180 | 28.83640 | 45.60300 |
| C | 27.50600 | 30.04550 | 46.55050 |
| O | 27.10820 | 31.14980 | 46.08590 |
| O | 27.87970 | 29.87170 | 47.76270 |
| H | 25.47110 | 28.04330 | 46.84960 |
| H | 25.88130 | 29.38090 | 44.30510 |
| H | 27.90150 | 27.95460 | 46.15890 |
| H | 28.29470 | 29.06330 | 44.82330 |
| N | 26.43600 | 26.10900 | 44.22600 |
| H | 26.15950 | 25.82240 | 45.15920 |
| C | 22.02800 | 35.96100 | 51.62100 |
| O | 21.52500 | 35.67200 | 52.73400 |
| N | 23.04900 | 35.23800 | 51.14800 |
| C | 23.74700 | 34.28100 | 52.02300 |
| C | 24.10700 | 34.76800 | 53.38500 |
| O | 23.80400 | 34.05500 | 54.34000 |
| C | 25.02000 | 33.77660 | 51.29490 |
| C | 24.74920 | 32.99380 | 50.00070 |
| C | 26.01430 | 32.45600 | 49.30750 |
| C | 26.86840 | 33.54260 | 48.64790 |
| N | 28.01140 | 32.92370 | 47.90780 |
| H | 23.50250 | 35.53130 | 50.28480 |
| H | 23.07230 | 33.43230 | 52.20360 |
| H | 25.67360 | 34.64420 | 51.09910 |
| H | 25.56700 | 33.13000 | 51.99850 |
| H | 24.09560 | 32.14100 | 50.24230 |
| H | 24.18420 | 33.61140 | 49.28220 |
| H | 26.63230 | 31.89780 | 50.02980 |
| H | 25.72110 | 31.73420 | 48.53030 |
| H | 26.27450 | 34.11780 | 47.92350 |
| H | 27.28400 | 34.24220 | 49.38600 |
| H | 28.59480 | 32.35000 | 48.56520 |
| H | 28.65250 | 33.61520 | 47.49150 |
| H | 27.64880 | 32.25510 | 47.13650 |
| N | 24.74100 | 35.92700 | 53.53200 |
| H | 24.85880 | 36.57360 | 52.75880 |
| H | 35.81620 | 29.27070 | 51.07340 |
| H | 33.84900 | 33.74900 | 50.00070 |
| H | 34.49770 | 31.23040 | 45.31340 |
| H | 35.21020 | 35.82300 | 44.14190 |
| H | 23.26550 | 27.63460 | 46.32760 |
| H | 26.53590 | 25.39890 | 43.50820 |
| H | 21.65640 | 36.76510 | 50.95420 |
| H | 24.88960 | 36.27940 | 54.47230 |

**Table S31.** Step 1 of the relaxed surface scan for proton transfer from complex 2 to complex 3 without explicit waters.

| | | | |
|---|---|---|---|
| C | 35.07600 | 29.61800 | 50.32400 |
| O | 35.29000 | 29.74100 | 49.12000 |
| N | 33.82600 | 29.84000 | 50.82400 |
| C | 32.80700 | 30.46300 | 50.04700 |
| C | 33.11400 | 31.92700 | 49.53200 |
| O | 32.91300 | 32.28500 | 48.36700 |
| C | 31.45510 | 30.42310 | 50.78900 |
| C | 30.29180 | 30.70380 | 49.84620 |
| O | 29.76050 | 31.81790 | 49.78970 |
| O | 29.95470 | 29.65170 | 49.13070 |
| H | 33.70510 | 29.79300 | 51.83450 |
| H | 32.70600 | 29.90200 | 49.10510 |
| H | 31.32700 | 29.41340 | 51.20380 |
| H | 31.43330 | 31.15650 | 51.60610 |
| H | 29.11010 | 29.80360 | 48.52270 |
| N | 33.63800 | 32.82200 | 50.35700 |
| H | 33.88040 | 32.60920 | 51.32000 |
| C | 33.90300 | 32.02700 | 44.81800 |
| O | 33.71700 | 32.08700 | 43.59100 |
| N | 33.39200 | 32.93300 | 45.68900 |
| C | 32.51000 | 34.06700 | 45.26000 |
| C | 33.32800 | 35.05900 | 44.40200 |
| O | 32.77200 | 35.51900 | 43.37500 |
| C | 31.81460 | 34.70490 | 46.48320 |

| | | | |
|---|---|---|---|
| C | 31.01210 | 35.93680 | 46.09350 |
| O | 30.89410 | 33.77280 | 47.08350 |
| H | 33.47720 | 32.73370 | 46.68890 |
| H | 31.73810 | 33.67730 | 44.58280 |
| H | 32.58990 | 34.98030 | 47.22370 |
| H | 31.42080 | 33.10900 | 47.58270 |
| H | 30.50810 | 36.33890 | 46.98250 |
| H | 30.25550 | 35.67450 | 45.34090 |
| H | 31.65760 | 36.71680 | 45.67040 |
| N | 34.61300 | 35.31200 | 44.78500 |
| H | 35.05100 | 34.70450 | 45.47030 |
| C | 23.94400 | 27.94100 | 45.50200 |
| O | 23.57900 | 28.09500 | 44.33100 |
| N | 25.20400 | 28.15800 | 45.87400 |
| C | 26.22300 | 28.52000 | 44.89200 |
| C | 26.47000 | 27.38700 | 43.85800 |
| O | 26.67800 | 27.71100 | 42.68900 |
| C | 27.54980 | 28.84190 | 45.60150 |
| C | 27.49540 | 30.05140 | 46.54790 |
| O | 27.10520 | 31.15750 | 46.07750 |
| O | 27.85650 | 29.87670 | 47.76240 |
| H | 25.47090 | 28.04520 | 46.84990 |
| H | 25.87900 | 29.37990 | 44.30480 |
| H | 27.90260 | 27.96220 | 46.15860 |
| H | 28.29210 | 29.07170 | 44.82210 |
| N | 26.43600 | 26.10900 | 44.22600 |
| H | 26.15890 | 25.82260 | 45.15910 |
| C | 22.02800 | 35.96100 | 51.62100 |
| O | 21.52500 | 35.67200 | 52.73400 |
| N | 23.04900 | 35.23800 | 51.14800 |
| C | 23.74700 | 34.28100 | 52.02300 |
| C | 24.10700 | 34.76800 | 53.38500 |
| O | 23.80400 | 34.05500 | 54.34000 |
| C | 25.01930 | 33.77630 | 51.29390 |
| C | 24.74640 | 32.99640 | 49.99820 |
| C | 26.00970 | 32.45520 | 49.30410 |
| C | 26.86610 | 33.54030 | 48.64480 |
| N | 28.00840 | 32.91980 | 47.90510 |
| H | 23.50230 | 35.53090 | 50.28460 |
| H | 23.07210 | 33.43250 | 52.20390 |
| H | 25.67410 | 34.64330 | 51.09970 |
| H | 25.56520 | 33.12700 | 51.99600 |
| H | 24.08940 | 32.14590 | 50.23840 |
| H | 24.18420 | 33.61690 | 49.28000 |
| H | 26.62640 | 31.89530 | 50.02610 |
| H | 25.71410 | 31.73470 | 48.52680 |
| H | 26.27320 | 34.11650 | 47.92040 |
| H | 27.28190 | 34.23880 | 49.38370 |
| H | 28.58760 | 32.34250 | 48.56140 |
| H | 28.65170 | 33.61120 | 47.49220 |
| H | 27.64530 | 32.25450 | 47.13010 |
| N | 24.74100 | 35.92700 | 53.53200 |
| H | 24.85880 | 36.57340 | 52.75860 |
| H | 35.81620 | 29.27050 | 51.07330 |
| H | 33.84920 | 33.74890 | 50.00060 |
| H | 34.49780 | 31.23040 | 45.31340 |
| H | 35.21010 | 35.82330 | 44.14210 |
| H | 23.26530 | 27.63480 | 46.32760 |
| H | 26.53540 | 25.39880 | 43.50820 |
| H | 21.65640 | 36.76500 | 50.95420 |
| H | 24.88930 | 36.27960 | 54.47220 |

**Table S32.** Optimized transition state for proton transfer from complex 2 to complex 3 without explicit waters.

| | | | |
|---|---|---|---|
| C | 35.07600 | 29.61800 | 50.32400 |
| O | 35.29000 | 29.74100 | 49.12000 |
| N | 33.82600 | 29.84000 | 50.82400 |
| C | 32.80700 | 30.46300 | 50.04700 |
| C | 33.11400 | 31.92700 | 49.53200 |
| O | 32.91300 | 32.28500 | 48.36700 |
| C | 31.47330 | 30.47550 | 50.81740 |
| C | 30.28040 | 30.84350 | 49.93000 |
| O | 29.55910 | 31.81980 | 50.23890 |
| O | 30.11780 | 30.07280 | 48.91190 |
| H | 33.70340 | 29.79240 | 51.83420 |
| H | 32.67030 | 29.89860 | 49.11350 |
| H | 31.29110 | 29.45780 | 51.19650 |
| H | 31.51770 | 31.16710 | 51.67000 |
| H | 29.00660 | 29.94880 | 48.35060 |
| N | 33.63800 | 32.82200 | 50.35700 |
| H | 33.87760 | 32.60880 | 51.32060 |
| C | 33.90300 | 32.02700 | 44.81800 |
| O | 33.71700 | 32.08700 | 43.59100 |
| N | 33.39200 | 32.93300 | 45.68900 |
| C | 32.51000 | 34.06700 | 45.26000 |
| C | 33.32800 | 35.05900 | 44.40200 |
| O | 32.77200 | 35.51900 | 43.37500 |
| C | 31.82030 | 34.70510 | 46.48620 |
| C | 30.99450 | 35.92160 | 46.09860 |
| O | 30.92460 | 33.76280 | 47.10850 |

| | | | |
|---|---|---|---|
| H | 33.47900 | 32.73260 | 46.68860 |
| H | 31.73840 | 33.67660 | 44.58290 |
| H | 32.60140 | 34.99890 | 47.21350 |
| H | 31.47510 | 33.09570 | 47.58030 |
| H | 30.50330 | 36.32810 | 46.99290 |
| H | 30.22600 | 35.64180 | 45.36450 |
| H | 31.62210 | 36.70380 | 45.65320 |
| N | 34.61300 | 35.31200 | 44.78500 |
| H | 35.05120 | 34.70470 | 45.47030 |
| C | 23.94400 | 27.94100 | 45.50200 |
| O | 23.57900 | 28.09500 | 44.33100 |
| N | 25.20400 | 28.15800 | 45.87400 |
| C | 26.22300 | 28.52000 | 44.89200 |
| C | 26.47000 | 27.38700 | 43.85800 |
| O | 26.67800 | 27.71100 | 42.68900 |
| C | 27.58280 | 28.73750 | 45.57950 |
| C | 27.71020 | 29.85710 | 46.60950 |
| O | 27.50970 | 31.04410 | 46.28550 |
| O | 28.08940 | 29.45000 | 47.78810 |
| H | 25.47230 | 28.03040 | 46.84770 |
| H | 25.90310 | 29.39550 | 44.31570 |
| H | 27.90370 | 27.79910 | 46.05170 |
| H | 28.30910 | 28.97540 | 44.78680 |
| N | 26.43600 | 26.10900 | 44.22600 |
| H | 26.16840 | 25.81840 | 45.16050 |
| C | 22.02800 | 35.96100 | 51.62100 |
| O | 21.52500 | 35.67200 | 52.73400 |
| N | 23.04900 | 35.23800 | 51.14800 |
| C | 23.74700 | 34.28100 | 52.02300 |
| C | 24.10700 | 34.76800 | 53.38500 |
| O | 23.80400 | 34.05500 | 54.34000 |
| C | 25.02980 | 33.78660 | 51.30650 |
| C | 24.79160 | 33.02010 | 49.99690 |
| C | 26.08040 | 32.47170 | 49.35750 |
| C | 26.97620 | 33.54860 | 48.73740 |
| N | 28.20150 | 32.91540 | 48.15800 |
| H | 23.50240 | 35.53150 | 50.28490 |
| H | 23.07560 | 33.42920 | 52.20030 |
| H | 25.68590 | 34.65710 | 51.13460 |
| H | 25.56520 | 33.13220 | 52.01190 |
| H | 24.11870 | 32.17370 | 50.20510 |
| H | 24.26480 | 33.65190 | 49.26190 |
| H | 26.66010 | 31.91260 | 50.11040 |
| H | 25.81390 | 31.75130 | 48.56830 |
| H | 26.45070 | 34.08090 | 47.93240 |
| H | 27.30510 | 34.28500 | 49.48300 |
| H | 28.73540 | 32.42340 | 48.93940 |
| H | 28.85740 | 33.58390 | 47.72200 |
| H | 27.93770 | 32.18710 | 47.42360 |
| N | 24.74100 | 35.92700 | 53.53200 |
| H | 24.86240 | 36.57300 | 52.75900 |
| H | 35.81760 | 29.27260 | 51.07330 |
| H | 33.84430 | 33.75100 | 50.00320 |
| H | 34.49740 | 31.23050 | 45.31370 |
| H | 35.20980 | 35.82340 | 44.14190 |
| H | 23.26650 | 27.63220 | 46.32730 |
| H | 26.55440 | 25.39990 | 43.51020 |
| H | 21.65650 | 36.76520 | 50.95430 |
| H | 24.89390 | 36.27710 | 54.47240 |

**Table S33.** Step 8 of the nudged elastic band for proton transfer from complex 2 to complex 3 without explicit waters.

| | | | |
|---|---|---|---|
| C | 35.07600 | 29.61800 | 50.32400 |
| O | 35.29000 | 29.74100 | 49.12000 |
| N | 33.82600 | 29.84000 | 50.82400 |
| C | 32.80700 | 30.46300 | 50.04700 |
| C | 33.11400 | 31.92700 | 49.53200 |
| O | 32.91300 | 32.28500 | 48.36700 |
| C | 31.48550 | 30.51210 | 50.83560 |
| C | 30.32150 | 31.02180 | 49.97500 |
| O | 29.56850 | 31.92100 | 50.44330 |
| O | 30.21010 | 30.48410 | 48.82700 |
| H | 33.70250 | 29.79240 | 51.83390 |
| H | 32.64830 | 29.89930 | 49.11790 |
| H | 31.23230 | 29.48790 | 51.15330 |
| H | 31.58660 | 31.13920 | 51.73300 |
| H | 29.01870 | 29.94660 | 48.21560 |
| N | 33.63800 | 32.82200 | 50.35700 |
| H | 33.86820 | 32.61150 | 51.32340 |
| C | 33.90300 | 32.02700 | 44.81800 |
| O | 33.71700 | 32.08700 | 43.59100 |
| N | 33.39200 | 32.93300 | 45.68900 |
| C | 32.51000 | 34.06700 | 45.26000 |
| C | 33.32800 | 35.05900 | 44.40200 |
| O | 32.77200 | 35.51900 | 43.37500 |
| C | 31.81220 | 34.69150 | 46.48910 |
| C | 30.98370 | 35.90830 | 46.10860 |
| O | 30.91770 | 33.73560 | 47.09350 |
| H | 33.47180 | 32.72920 | 46.68900 |
| H | 31.74050 | 33.67620 | 44.58050 |

| | | | |
|---|---|---|---|
| H | 32.58730 | 34.98000 | 47.22450 |
| H | 31.46320 | 33.08240 | 47.59200 |
| H | 30.48950 | 36.30810 | 47.00420 |
| H | 30.21740 | 35.63110 | 45.37120 |
| H | 31.61030 | 36.69490 | 45.66930 |
| N | 34.61300 | 35.31200 | 44.78500 |
| H | 35.05010 | 34.70500 | 45.47150 |
| C | 23.94400 | 27.94100 | 45.50200 |
| O | 23.57900 | 28.09500 | 44.33100 |
| N | 25.20400 | 28.15800 | 45.87400 |
| C | 26.22300 | 28.52000 | 44.89200 |
| C | 26.47000 | 27.38700 | 43.85800 |
| O | 26.67800 | 27.71100 | 42.68900 |
| C | 27.59360 | 28.68800 | 45.56490 |
| C | 27.78990 | 29.75100 | 46.62780 |
| O | 27.46440 | 30.93420 | 46.47920 |
| O | 28.41780 | 29.25840 | 47.67940 |
| H | 25.47010 | 28.02710 | 46.84750 |
| H | 25.91410 | 29.40290 | 44.32200 |
| H | 27.91100 | 27.72550 | 45.98780 |
| H | 28.31650 | 28.93980 | 44.77150 |
| N | 26.43600 | 26.10900 | 44.22600 |
| H | 26.15410 | 25.81660 | 45.15570 |
| C | 22.02800 | 35.96100 | 51.62100 |
| O | 21.52500 | 35.67200 | 52.73400 |
| N | 23.04900 | 35.23800 | 51.14800 |
| C | 23.74700 | 34.28100 | 52.02300 |
| C | 24.10700 | 34.76800 | 53.38500 |
| O | 23.80400 | 34.05500 | 54.34000 |
| C | 25.03180 | 33.79120 | 51.31160 |
| C | 24.80320 | 33.03660 | 49.99490 |
| C | 26.10050 | 32.48760 | 49.37730 |
| C | 27.01220 | 33.56200 | 48.77600 |
| N | 28.27570 | 32.92370 | 48.29460 |
| H | 23.50210 | 35.53110 | 50.28450 |
| H | 23.07670 | 33.42820 | 52.19930 |
| H | 25.69060 | 34.66190 | 51.15150 |
| H | 25.56140 | 33.13040 | 52.01550 |
| H | 24.12240 | 32.19280 | 50.18730 |
| H | 24.29200 | 33.67680 | 49.25640 |
| H | 26.66490 | 31.92990 | 50.14320 |
| H | 25.85010 | 31.76800 | 48.58210 |
| H | 26.52590 | 34.06330 | 47.92770 |
| H | 27.29350 | 34.32410 | 49.51430 |
| H | 28.78150 | 32.48510 | 49.14060 |
| H | 28.94270 | 33.57130 | 47.84120 |
| H | 28.05310 | 32.15900 | 47.60690 |
| N | 24.74100 | 35.92700 | 53.53200 |
| H | 24.86330 | 36.57300 | 52.75910 |
| H | 35.81810 | 29.27230 | 51.07290 |
| H | 33.83790 | 33.75310 | 50.00550 |
| H | 34.49640 | 31.22980 | 45.31370 |
| H | 35.21070 | 35.82170 | 44.14130 |
| H | 23.26700 | 27.63180 | 46.32750 |
| H | 26.55100 | 25.40130 | 43.50810 |
| H | 21.65670 | 36.76540 | 50.95440 |
| H | 24.89590 | 36.27620 | 54.47240 |

**Table S34.** Step 7 of the nudged elastic band for proton transfer from complex 2 to complex 3 without explicit waters.

| | | | |
|---|---|---|---|
| C | 35.07600 | 29.61800 | 50.32400 |
| O | 35.29000 | 29.74100 | 49.12000 |
| N | 33.82600 | 29.84000 | 50.82400 |
| C | 32.80700 | 30.46300 | 50.04700 |
| C | 33.11400 | 31.92700 | 49.53200 |
| O | 32.91300 | 32.28500 | 48.36700 |
| C | 31.48310 | 30.50080 | 50.83220 |
| C | 30.31280 | 30.94830 | 49.94730 |
| O | 29.59010 | 31.90970 | 50.32440 |
| O | 30.17220 | 30.28460 | 48.86720 |
| H | 33.70280 | 29.79410 | 51.83400 |
| H | 32.65300 | 29.89790 | 49.11760 |
| H | 31.26270 | 29.47900 | 51.17960 |
| H | 31.56300 | 31.15990 | 51.70850 |
| H | 28.96540 | 29.96260 | 48.26550 |
| N | 33.63800 | 32.82200 | 50.35700 |
| H | 33.87540 | 32.60930 | 51.32110 |
| C | 33.90300 | 32.02700 | 44.81800 |
| O | 33.71700 | 32.08700 | 43.59100 |
| N | 33.39200 | 32.93300 | 45.68900 |
| C | 32.51000 | 34.06700 | 45.26000 |
| C | 33.32800 | 35.05900 | 44.40200 |
| O | 32.77200 | 35.51900 | 43.37500 |
| C | 31.81970 | 34.69630 | 46.49030 |
| C | 30.99820 | 35.91860 | 46.11280 |
| O | 30.92160 | 33.74850 | 47.10030 |
| H | 33.47770 | 32.73300 | 46.68910 |
| H | 31.73810 | 33.67740 | 44.58280 |
| H | 32.60030 | 34.98070 | 47.22200 |
| H | 31.46870 | 33.08090 | 47.57680 |

| | | | |
|---|---|---|---|
| H | 30.50370 | 36.31630 | 47.00910 |
| H | 30.23280 | 35.64880 | 45.37180 |
| H | 31.63040 | 36.70390 | 45.67940 |
| N | 34.61300 | 35.31200 | 44.78500 |
| H | 35.05000 | 34.70810 | 45.47390 |
| C | 23.94400 | 27.94100 | 45.50200 |
| O | 23.57900 | 28.09500 | 44.33100 |
| N | 25.20400 | 28.15800 | 45.87400 |
| C | 26.22300 | 28.52000 | 44.89200 |
| C | 26.47000 | 27.38700 | 43.85800 |
| O | 26.67800 | 27.71100 | 42.68900 |
| C | 27.58390 | 28.71540 | 45.57210 |
| C | 27.73230 | 29.80290 | 46.62030 |
| O | 27.44990 | 30.98720 | 46.39410 |
| O | 28.25050 | 29.33860 | 47.73610 |
| H | 25.47070 | 28.02500 | 46.84720 |
| H | 25.90640 | 29.39800 | 44.31810 |
| H | 27.90900 | 27.76580 | 46.01730 |
| H | 28.31000 | 28.97200 | 44.78410 |
| N | 26.43600 | 26.10900 | 44.22600 |
| H | 26.16650 | 25.81680 | 45.15940 |
| C | 22.02800 | 35.96100 | 51.62100 |
| O | 21.52500 | 35.67200 | 52.73400 |
| N | 23.04900 | 35.23800 | 51.14800 |
| C | 23.74700 | 34.28100 | 52.02300 |
| C | 24.10700 | 34.76800 | 53.38500 |
| O | 23.80400 | 34.05500 | 54.34000 |
| C | 25.03180 | 33.78850 | 51.31270 |
| C | 24.79860 | 33.02220 | 50.00360 |
| C | 26.09160 | 32.47670 | 49.37220 |
| C | 26.98980 | 33.55520 | 48.75800 |
| N | 28.23250 | 32.92410 | 48.21440 |
| H | 23.50430 | 35.53450 | 50.28680 |
| H | 23.07570 | 33.42890 | 52.19940 |
| H | 25.68830 | 34.65940 | 51.14450 |
| H | 25.56390 | 33.13460 | 52.02120 |
| H | 24.12630 | 32.17500 | 50.21030 |
| H | 24.27340 | 33.65340 | 49.26700 |
| H | 26.66750 | 31.92090 | 50.13060 |
| H | 25.83110 | 31.75580 | 48.58140 |
| H | 26.47540 | 34.07350 | 47.93670 |
| H | 27.29800 | 34.30290 | 49.50080 |
| H | 28.77350 | 32.46440 | 49.02260 |
| H | 28.88240 | 33.58530 | 47.75830 |
| H | 27.97900 | 32.17840 | 47.50830 |
| N | 24.74100 | 35.92700 | 53.53200 |
| H | 24.86830 | 36.57130 | 52.75850 |
| H | 35.81790 | 29.27210 | 51.07310 |
| H | 33.84330 | 33.75120 | 50.00360 |
| H | 34.49750 | 31.23040 | 45.31340 |
| H | 35.21000 | 35.82460 | 44.14320 |
| H | 23.26640 | 27.63190 | 46.32710 |
| H | 26.55780 | 25.40030 | 43.51020 |
| H | 21.65740 | 36.76610 | 50.95500 |
| H | 24.90090 | 36.27420 | 54.47210 |

**Table S35.** Step 6 of the nudged elastic band for proton transfer from complex 2 to complex 3 without explicit waters.

| | | | |
|---|---|---|---|
| C | 35.07600 | 29.61800 | 50.32400 |
| O | 35.29000 | 29.74100 | 49.12000 |
| N | 33.82600 | 29.84000 | 50.82400 |
| C | 32.80700 | 30.46300 | 50.04700 |
| C | 33.11400 | 31.92700 | 49.53200 |
| O | 32.91300 | 32.28500 | 48.36700 |
| C | 31.47570 | 30.49440 | 50.82220 |
| C | 30.30450 | 30.91200 | 49.92040 |
| O | 29.60440 | 31.90680 | 50.24530 |
| O | 30.13960 | 30.18610 | 48.88200 |
| H | 33.70010 | 29.79180 | 51.83360 |
| H | 32.65570 | 29.89790 | 49.11660 |
| H | 31.26930 | 29.47490 | 51.18390 |
| H | 31.53980 | 31.16870 | 51.68840 |
| H | 28.92570 | 29.96120 | 48.27380 |
| N | 33.63800 | 32.82200 | 50.35700 |
| H | 33.87130 | 32.61090 | 51.32240 |
| C | 33.90300 | 32.02700 | 44.81800 |
| O | 33.71700 | 32.08700 | 43.59100 |
| N | 33.39200 | 32.93300 | 45.68900 |
| C | 32.51000 | 34.06700 | 45.26000 |
| C | 33.32800 | 35.05900 | 44.40200 |
| O | 32.77200 | 35.51900 | 43.37500 |
| C | 31.81390 | 34.69800 | 46.48580 |
| C | 30.99380 | 35.92060 | 46.10400 |
| O | 30.91190 | 33.75100 | 47.09190 |
| H | 33.47280 | 32.73060 | 46.68920 |
| H | 31.73940 | 33.67750 | 44.58080 |
| H | 32.59070 | 34.98290 | 47.22080 |
| H | 31.45620 | 33.08530 | 47.57310 |
| H | 30.49670 | 36.31880 | 46.99880 |
| H | 30.22990 | 35.65230 | 45.36070 |

| | | | |
|---|---|---|---|
| H | 31.62720 | 36.70600 | 45.67260 |
| N | 34.61300 | 35.31200 | 44.78500 |
| H | 35.05050 | 34.70510 | 45.47110 |
| C | 23.94400 | 27.94100 | 45.50200 |
| O | 23.57900 | 28.09500 | 44.33100 |
| N | 25.20400 | 28.15800 | 45.87400 |
| C | 26.22300 | 28.52000 | 44.89200 |
| C | 26.47000 | 27.38700 | 43.85800 |
| O | 26.67800 | 27.71100 | 42.68900 |
| C | 27.57780 | 28.73030 | 45.56910 |
| C | 27.69490 | 29.83020 | 46.60460 |
| O | 27.42700 | 31.01270 | 46.34480 |
| O | 28.15920 | 29.38190 | 47.74810 |
| H | 25.46950 | 28.02390 | 46.84750 |
| H | 25.89900 | 29.39330 | 44.31450 |
| H | 27.90570 | 27.78990 | 46.03070 |
| H | 28.30450 | 28.98740 | 44.78260 |
| N | 26.43600 | 26.10900 | 44.22600 |
| H | 26.16040 | 25.81710 | 45.15780 |
| C | 22.02800 | 35.96100 | 51.62100 |
| O | 21.52500 | 35.67200 | 52.73400 |
| N | 23.04900 | 35.23800 | 51.14800 |
| C | 23.74700 | 34.28100 | 52.02300 |
| C | 24.10700 | 34.76800 | 53.38500 |
| O | 23.80400 | 34.05500 | 54.34000 |
| C | 25.02630 | 33.78560 | 51.30620 |
| C | 24.78390 | 33.01670 | 49.99960 |
| C | 26.07310 | 32.47300 | 49.35830 |
| C | 26.96450 | 33.55370 | 48.73820 |
| N | 28.19690 | 32.92620 | 48.16770 |
| H | 23.50020 | 35.53160 | 50.28370 |
| H | 23.07270 | 33.43100 | 52.19930 |
| H | 25.68100 | 34.65630 | 51.13110 |
| H | 25.56200 | 33.13260 | 52.01240 |
| H | 24.11460 | 32.16850 | 50.21210 |
| H | 24.25170 | 33.64510 | 49.26550 |
| H | 26.65570 | 31.91710 | 50.11150 |
| H | 25.80690 | 31.75210 | 48.56940 |
| H | 26.43740 | 34.07890 | 47.92930 |
| H | 27.28580 | 34.29550 | 49.48120 |
| H | 28.75300 | 32.45440 | 48.95780 |
| H | 28.83980 | 33.59320 | 47.71110 |
| H | 27.92940 | 32.19070 | 47.45330 |
| N | 24.74100 | 35.92700 | 53.53200 |
| H | 24.86050 | 36.57230 | 52.75800 |
| H | 35.81710 | 29.27010 | 51.07270 |
| H | 33.83810 | 33.75230 | 50.00320 |
| H | 34.49470 | 31.22830 | 45.31300 |
| H | 35.20860 | 35.82330 | 44.14080 |
| H | 23.26460 | 27.63250 | 46.32580 |
| H | 26.54720 | 25.40120 | 43.50760 |
| H | 21.65410 | 36.76310 | 50.95310 |
| H | 24.89340 | 36.27820 | 54.47200 |

**Table S36.** Step 5 of the nudged elastic band for proton transfer from complex 2 to complex 3 without explicit waters.

| | | | |
|---|---|---|---|
| C | 35.07600 | 29.61800 | 50.32400 |
| O | 35.29000 | 29.74100 | 49.12000 |
| N | 33.82600 | 29.84000 | 50.82400 |
| C | 32.80700 | 30.46300 | 50.04700 |
| C | 33.11400 | 31.92700 | 49.53200 |
| O | 32.91300 | 32.28500 | 48.36700 |
| C | 31.47270 | 30.48730 | 50.81570 |
| C | 30.29670 | 30.89550 | 49.92580 |
| O | 29.58250 | 31.87090 | 50.24290 |
| O | 30.14530 | 30.14680 | 48.88570 |
| H | 33.70240 | 29.78930 | 51.83380 |
| H | 32.66820 | 29.89860 | 49.11410 |
| H | 31.26620 | 29.46670 | 51.17350 |
| H | 31.52550 | 31.16300 | 51.68060 |
| H | 29.06150 | 29.99050 | 48.34310 |
| N | 33.63800 | 32.82200 | 50.35700 |
| H | 33.87200 | 32.61190 | 51.32260 |
| C | 33.90300 | 32.02700 | 44.81800 |
| O | 33.71700 | 32.08700 | 43.59100 |
| N | 33.39200 | 32.93300 | 45.68900 |
| C | 32.51000 | 34.06700 | 45.26000 |
| C | 33.32800 | 35.05900 | 44.40200 |
| O | 32.77200 | 35.51900 | 43.37500 |
| C | 31.81020 | 34.69850 | 46.48310 |
| C | 30.98760 | 35.91770 | 46.09640 |
| O | 30.90920 | 33.75010 | 47.08940 |
| H | 33.47540 | 32.73100 | 46.68870 |
| H | 31.74220 | 33.67550 | 44.57920 |
| H | 32.58350 | 34.98740 | 47.22050 |
| H | 31.45560 | 33.08310 | 47.56530 |
| H | 30.48830 | 36.31770 | 46.98920 |
| H | 30.22580 | 35.64460 | 45.35290 |
| H | 31.61960 | 36.70330 | 45.66310 |
| N | 34.61300 | 35.31200 | 44.78500 |

| | | | |
|---|---|---|---|
| H | 35.05200 | 34.70220 | 45.46760 |
| C | 23.94400 | 27.94100 | 45.50200 |
| O | 23.57900 | 28.09500 | 44.33100 |
| N | 25.20400 | 28.15800 | 45.87400 |
| C | 26.22300 | 28.52000 | 44.89200 |
| C | 26.47000 | 27.38700 | 43.85800 |
| O | 26.67800 | 27.71100 | 42.68900 |
| C | 27.58210 | 28.73970 | 45.57700 |
| C | 27.69980 | 29.84480 | 46.62470 |
| O | 27.42280 | 31.02610 | 46.33490 |
| O | 28.15430 | 29.43490 | 47.77180 |
| H | 25.47230 | 28.02840 | 46.84740 |
| H | 25.90020 | 29.39420 | 44.31560 |
| H | 27.91660 | 27.79670 | 46.03030 |
| H | 28.30280 | 28.99930 | 44.78530 |
| N | 26.43600 | 26.10900 | 44.22600 |
| H | 26.15210 | 25.81870 | 45.15580 |
| C | 22.02800 | 35.96100 | 51.62100 |
| O | 21.52500 | 35.67200 | 52.73400 |
| N | 23.04900 | 35.23800 | 51.14800 |
| C | 23.74700 | 34.28100 | 52.02300 |
| C | 24.10700 | 34.76800 | 53.38500 |
| O | 23.80400 | 34.05500 | 54.34000 |
| C | 25.02510 | 33.78480 | 51.30200 |
| C | 24.77890 | 33.01920 | 49.99350 |
| C | 26.06590 | 32.47380 | 49.34970 |
| C | 26.95670 | 33.55290 | 48.72660 |
| N | 28.18430 | 32.92030 | 48.15130 |
| H | 23.49930 | 35.52750 | 50.28190 |
| H | 23.07430 | 33.43040 | 52.20160 |
| H | 25.68110 | 34.65470 | 51.12650 |
| H | 25.56320 | 33.12930 | 52.00420 |
| H | 24.10780 | 32.17200 | 50.20420 |
| H | 24.24810 | 33.65050 | 49.26100 |
| H | 26.64900 | 31.91770 | 50.10240 |
| H | 25.80080 | 31.75230 | 48.56120 |
| H | 26.43010 | 34.07850 | 47.91790 |
| H | 27.28140 | 34.29510 | 49.46830 |
| H | 28.73160 | 32.45410 | 48.93750 |
| H | 28.82940 | 33.58540 | 47.69550 |
| H | 27.91260 | 32.17320 | 47.43680 |
| N | 24.74100 | 35.92700 | 53.53200 |
| H | 24.85710 | 36.57420 | 52.75900 |
| H | 35.81690 | 29.27040 | 51.07310 |
| H | 33.83800 | 33.75290 | 50.00450 |
| H | 34.49540 | 31.22910 | 45.31400 |
| H | 35.21010 | 35.82010 | 44.13950 |
| H | 23.26640 | 27.63260 | 46.32740 |
| H | 26.54180 | 25.40100 | 43.50700 |
| H | 21.65450 | 36.76310 | 50.95300 |
| H | 24.88890 | 36.28010 | 54.47210 |

**Table S37.** Step 4 of the nudged elastic band for proton transfer from complex 2 to complex 3 without explicit waters.

| | | | |
|---|---|---|---|
| C | 35.07600 | 29.61800 | 50.32400 |
| O | 35.29000 | 29.74100 | 49.12000 |
| N | 33.82600 | 29.84000 | 50.82400 |
| C | 32.80700 | 30.46300 | 50.04700 |
| C | 33.11400 | 31.92700 | 49.53200 |
| O | 32.91300 | 32.28500 | 48.36700 |
| C | 31.46610 | 30.48340 | 50.80690 |
| C | 30.28880 | 30.88460 | 49.91900 |
| O | 29.57020 | 31.85460 | 50.22100 |
| O | 30.13900 | 30.11380 | 48.88390 |
| H | 33.69790 | 29.78570 | 51.83200 |
| H | 32.67000 | 29.89960 | 49.11090 |
| H | 31.25920 | 29.46280 | 51.16440 |
| H | 31.50910 | 31.16130 | 51.67010 |
| H | 29.11880 | 30.01320 | 48.38360 |
| N | 33.63800 | 32.82200 | 50.35700 |
| H | 33.86420 | 32.61370 | 51.32330 |
| C | 33.90300 | 32.02700 | 44.81800 |
| O | 33.71700 | 32.08700 | 43.59100 |
| N | 33.39200 | 32.93300 | 45.68900 |
| C | 32.51000 | 34.06700 | 45.26000 |
| C | 33.32800 | 35.05900 | 44.40200 |
| O | 32.77200 | 35.51900 | 43.37500 |
| C | 31.79900 | 34.69880 | 46.47580 |
| C | 30.97390 | 35.91500 | 46.08270 |
| O | 30.89630 | 33.74890 | 47.07830 |
| H | 33.46910 | 32.72860 | 46.68720 |
| H | 31.74260 | 33.67470 | 44.57490 |
| H | 32.56620 | 34.99130 | 47.21710 |
| H | 31.44180 | 33.08160 | 47.55350 |
| H | 30.47160 | 36.31660 | 46.97320 |
| H | 30.21400 | 35.63840 | 45.33850 |
| H | 31.60440 | 36.70120 | 45.64860 |
| N | 34.61300 | 35.31200 | 44.78500 |
| H | 35.04990 | 34.69710 | 45.46250 |
| C | 23.94400 | 27.94100 | 45.50200 |

| | | | |
|---|---|---|---|
| O | 23.57900 | 28.09500 | 44.33100 |
| N | 25.20400 | 28.15800 | 45.87400 |
| C | 26.22300 | 28.52000 | 44.89200 |
| C | 26.47000 | 27.38700 | 43.85800 |
| O | 26.67800 | 27.71100 | 42.68900 |
| C | 27.57940 | 28.74880 | 45.57150 |
| C | 27.67810 | 29.85610 | 46.62210 |
| O | 27.40300 | 31.03890 | 46.30890 |
| O | 28.10160 | 29.46770 | 47.77840 |
| H | 25.47000 | 28.03070 | 46.84680 |
| H | 25.89360 | 29.39160 | 44.31230 |
| H | 27.91760 | 27.80700 | 46.02510 |
| H | 28.29430 | 29.01020 | 44.77610 |
| N | 26.43600 | 26.10900 | 44.22600 |
| H | 26.13530 | 25.82000 | 45.15020 |
| C | 22.02800 | 35.96100 | 51.62100 |
| O | 21.52500 | 35.67200 | 52.73400 |
| N | 23.04900 | 35.23800 | 51.14800 |
| C | 23.74700 | 34.28100 | 52.02300 |
| C | 24.10700 | 34.76800 | 53.38500 |
| O | 23.80400 | 34.05500 | 54.34000 |
| C | 25.01720 | 33.78250 | 51.29250 |
| C | 24.76260 | 33.01950 | 49.98320 |
| C | 26.04650 | 32.47310 | 49.33420 |
| C | 26.93590 | 33.55160 | 48.70850 |
| N | 28.15880 | 32.91660 | 48.12550 |
| H | 23.49150 | 35.52210 | 50.27570 |
| H | 23.06990 | 33.43220 | 52.20010 |
| H | 25.67310 | 34.65160 | 51.11370 |
| H | 25.55770 | 33.12470 | 51.98980 |
| H | 24.09050 | 32.17320 | 50.19430 |
| H | 24.23060 | 33.65240 | 49.25270 |
| H | 26.63160 | 31.91630 | 50.08490 |
| H | 25.78000 | 31.75150 | 48.54650 |
| H | 26.40720 | 34.07850 | 47.90210 |
| H | 27.26460 | 34.29300 | 49.44930 |
| H | 28.70440 | 32.45060 | 48.90540 |
| H | 28.80380 | 33.58210 | 47.67050 |
| H | 27.88140 | 32.16520 | 47.40770 |
| N | 24.74100 | 35.92700 | 53.53200 |
| H | 24.84530 | 36.57570 | 52.75730 |
| H | 35.81370 | 29.26880 | 51.07130 |
| H | 33.82900 | 33.75440 | 50.00280 |
| H | 34.49020 | 31.22690 | 45.31210 |
| H | 35.20680 | 35.81680 | 44.13380 |
| H | 23.26290 | 27.63320 | 46.32470 |
| H | 26.52610 | 25.40250 | 43.50230 |
| H | 21.64900 | 36.76010 | 50.94930 |
| H | 24.87590 | 36.28480 | 54.47100 |

**Table S38.** Step 3 of the nudged elastic band for proton transfer from complex 2 to complex 3 without explicit waters.

| | | | |
|---|---|---|---|
| C | 35.07600 | 29.61800 | 50.32400 |
| O | 35.29000 | 29.74100 | 49.12000 |
| N | 33.82600 | 29.84000 | 50.82400 |
| C | 32.80700 | 30.46300 | 50.04700 |
| C | 33.11400 | 31.92700 | 49.53200 |
| O | 32.91300 | 32.28500 | 48.36700 |
| C | 31.46590 | 30.47650 | 50.80600 |
| C | 30.29110 | 30.86360 | 49.91480 |
| O | 29.57050 | 31.84170 | 50.19040 |
| O | 30.14010 | 30.06680 | 48.89500 |
| H | 33.69650 | 29.78370 | 51.83190 |
| H | 32.67360 | 29.90010 | 49.10940 |
| H | 31.26580 | 29.45540 | 51.16690 |
| H | 31.50240 | 31.15880 | 51.66560 |
| H | 29.15330 | 30.01770 | 48.41020 |
| N | 33.63800 | 32.82200 | 50.35700 |
| H | 33.86090 | 32.61240 | 51.32430 |
| C | 33.90300 | 32.02700 | 44.81800 |
| O | 33.71700 | 32.08700 | 43.59100 |
| N | 33.39200 | 32.93300 | 45.68900 |
| C | 32.51000 | 34.06700 | 45.26000 |
| C | 33.32800 | 35.05900 | 44.40200 |
| O | 32.77200 | 35.51900 | 43.37500 |
| C | 31.79150 | 34.69830 | 46.47090 |
| C | 30.96540 | 35.91210 | 46.07190 |
| O | 30.88770 | 33.74750 | 47.07070 |
| H | 33.46790 | 32.72820 | 46.68740 |
| H | 31.74430 | 33.67390 | 44.57250 |
| H | 32.55400 | 34.99360 | 47.21500 |
| H | 31.43280 | 33.08080 | 47.54640 |
| H | 30.45950 | 36.31580 | 46.95950 |
| H | 30.20860 | 35.63120 | 45.32610 |
| H | 31.59570 | 36.69810 | 45.63710 |
| N | 34.61300 | 35.31200 | 44.78500 |
| H | 35.04820 | 34.69200 | 45.45980 |
| C | 23.94400 | 27.94100 | 45.50200 |
| O | 23.57900 | 28.09500 | 44.33100 |
| N | 25.20400 | 28.15800 | 45.87400 |

| | | | |
|---|---|---|---|
| C | 26.22300 | 28.52000 | 44.89200 |
| C | 26.47000 | 27.38700 | 43.85800 |
| O | 26.67800 | 27.71100 | 42.68900 |
| C | 27.57650 | 28.76340 | 45.57290 |
| C | 27.65120 | 29.88000 | 46.61810 |
| O | 27.38000 | 31.05950 | 46.27170 |
| O | 28.04130 | 29.52270 | 47.78970 |
| H | 25.46960 | 28.03380 | 46.84790 |
| H | 25.89110 | 29.39000 | 44.31030 |
| H | 27.92430 | 27.82680 | 46.02890 |
| H | 28.28780 | 29.02820 | 44.77620 |
| N | 26.43600 | 26.10900 | 44.22600 |
| H | 26.12440 | 25.82200 | 45.14740 |
| C | 22.02800 | 35.96100 | 51.62100 |
| O | 21.52500 | 35.67200 | 52.73400 |
| N | 23.04900 | 35.23800 | 51.14800 |
| C | 23.74700 | 34.28100 | 52.02300 |
| C | 24.10700 | 34.76800 | 53.38500 |
| O | 23.80400 | 34.05500 | 54.34000 |
| C | 25.01290 | 33.78020 | 51.28560 |
| C | 24.75020 | 33.01870 | 49.97660 |
| C | 26.03010 | 32.47120 | 49.32050 |
| C | 26.91660 | 33.54900 | 48.68970 |
| N | 28.13140 | 32.91070 | 48.09390 |
| H | 23.48780 | 35.51810 | 50.27240 |
| H | 23.06890 | 33.43310 | 52.19980 |
| H | 25.66970 | 34.64760 | 51.10370 |
| H | 25.55560 | 33.12090 | 51.97890 |
| H | 24.07790 | 32.17310 | 50.18970 |
| H | 24.21600 | 33.65370 | 49.24970 |
| H | 26.61900 | 31.91360 | 50.06750 |
| H | 25.76020 | 31.74940 | 48.53430 |
| H | 26.38300 | 34.07870 | 47.88860 |
| H | 27.25340 | 34.28860 | 49.42890 |
| H | 28.67890 | 32.44020 | 48.86340 |
| H | 28.77540 | 33.57650 | 47.63840 |
| H | 27.84390 | 32.15940 | 47.37100 |
| N | 24.74100 | 35.92700 | 53.53200 |
| H | 24.83840 | 36.57660 | 52.75730 |
| H | 35.81320 | 29.26820 | 51.07060 |
| H | 33.82500 | 33.75490 | 50.00170 |
| H | 34.48950 | 31.22650 | 45.31090 |
| H | 35.20550 | 35.81420 | 44.13000 |
| H | 23.26190 | 27.63340 | 46.32330 |
| H | 26.51570 | 25.40350 | 43.50040 |
| H | 21.64810 | 36.75970 | 50.94810 |
| H | 24.86790 | 36.28690 | 54.47190 |

**Table S39.** Step 2 of the nudged elastic band for proton transfer from complex 2 to complex 3 without explicit waters.

| | | | |
|---|---|---|---|
| C | 35.07600 | 29.61800 | 50.32400 |
| O | 35.29000 | 29.74100 | 49.12000 |
| N | 33.82600 | 29.84000 | 50.82400 |
| C | 32.80700 | 30.46300 | 50.04700 |
| C | 33.11400 | 31.92700 | 49.53200 |
| O | 32.91300 | 32.28500 | 48.36700 |
| C | 31.46170 | 30.46750 | 50.79950 |
| C | 30.28850 | 30.83590 | 49.89910 |
| O | 29.59900 | 31.83810 | 50.11230 |
| O | 30.11010 | 29.97150 | 48.93050 |
| H | 33.69900 | 29.78050 | 51.83250 |
| H | 32.68270 | 29.89960 | 49.11010 |
| H | 31.27170 | 29.44790 | 51.16730 |
| H | 31.48020 | 31.16030 | 51.65080 |
| H | 29.15700 | 29.99880 | 48.43420 |
| N | 33.63800 | 32.82200 | 50.35700 |
| H | 33.86080 | 32.61830 | 51.32590 |
| C | 33.90300 | 32.02700 | 44.81800 |
| O | 33.71700 | 32.08700 | 43.59100 |
| N | 33.39200 | 32.93300 | 45.68900 |
| C | 32.51000 | 34.06700 | 45.26000 |
| C | 33.32800 | 35.05900 | 44.40200 |
| O | 32.77200 | 35.51900 | 43.37500 |
| C | 31.78940 | 34.70030 | 46.46950 |
| C | 30.96560 | 35.91520 | 46.06690 |
| O | 30.88090 | 33.75070 | 47.06390 |
| H | 33.46470 | 32.72410 | 46.68740 |
| H | 31.74880 | 33.67230 | 44.57190 |
| H | 32.54790 | 34.99580 | 47.21900 |
| H | 31.42140 | 33.08460 | 47.54440 |
| H | 30.45820 | 36.31850 | 46.95370 |
| H | 30.20910 | 35.63720 | 45.31960 |
| H | 31.59690 | 36.70140 | 45.63400 |
| N | 34.61300 | 35.31200 | 44.78500 |
| H | 35.05390 | 34.69160 | 45.45620 |
| C | 23.94400 | 27.94100 | 45.50200 |
| O | 23.57900 | 28.09500 | 44.33100 |
| N | 25.20400 | 28.15800 | 45.87400 |
| C | 26.22300 | 28.52000 | 44.89200 |
| C | 26.47000 | 27.38700 | 43.85800 |

| | | | |
|---|---|---|---|
| O | 26.67800 | 27.71100 | 42.68900 |
| C | 27.57460 | 28.77900 | 45.57050 |
| C | 27.61270 | 29.91320 | 46.60120 |
| O | 27.32900 | 31.08240 | 46.22570 |
| O | 27.96630 | 29.58580 | 47.79000 |
| H | 25.47210 | 28.03710 | 46.84800 |
| H | 25.88450 | 29.38480 | 44.30890 |
| H | 27.92350 | 27.85210 | 46.04730 |
| H | 28.28550 | 29.03590 | 44.76990 |
| N | 26.43600 | 26.10900 | 44.22600 |
| H | 26.11920 | 25.82190 | 45.14600 |
| C | 22.02800 | 35.96100 | 51.62100 |
| O | 21.52500 | 35.67200 | 52.73400 |
| N | 23.04900 | 35.23800 | 51.14800 |
| C | 23.74700 | 34.28100 | 52.02300 |
| C | 24.10700 | 34.76800 | 53.38500 |
| O | 23.80400 | 34.05500 | 54.34000 |
| C | 25.01050 | 33.77750 | 51.28190 |
| C | 24.74040 | 33.01330 | 49.97470 |
| C | 26.01730 | 32.46840 | 49.31070 |
| C | 26.89710 | 33.54770 | 48.67340 |
| N | 28.09870 | 32.91130 | 48.05000 |
| H | 23.48700 | 35.51320 | 50.27110 |
| H | 23.06940 | 33.43410 | 52.20480 |
| H | 25.66510 | 34.64560 | 51.09290 |
| H | 25.55850 | 33.11850 | 51.97260 |
| H | 24.07100 | 32.16640 | 50.19200 |
| H | 24.20000 | 33.64470 | 49.24890 |
| H | 26.61240 | 31.91210 | 50.05380 |
| H | 25.74520 | 31.74540 | 48.52650 |
| H | 26.35140 | 34.08480 | 47.88510 |
| H | 27.24790 | 34.28140 | 49.41220 |
| H | 28.65700 | 32.42740 | 48.80130 |
| H | 28.73970 | 33.58130 | 47.59790 |
| H | 27.79550 | 32.16990 | 47.31850 |
| N | 24.74100 | 35.92700 | 53.53200 |
| H | 24.83500 | 36.57850 | 52.75870 |
| H | 35.81420 | 29.26590 | 51.07260 |
| H | 33.82450 | 33.75620 | 50.00560 |
| H | 34.48810 | 31.22450 | 45.31450 |
| H | 35.20890 | 35.81150 | 44.13150 |
| H | 23.26490 | 27.63390 | 46.32710 |
| H | 26.50950 | 25.40370 | 43.49950 |
| H | 21.64630 | 36.75550 | 50.94800 |
| H | 24.86390 | 36.29130 | 54.47090 |

85

**Table S40.** Step 1 of the nudged elastic band for proton transfer from complex 2 to complex 3 without explicit waters.

| | | | |
|---|---|---|---|
| C | 35.07600 | 29.61800 | 50.32400 |
| O | 35.29000 | 29.74100 | 49.12000 |
| N | 33.82600 | 29.84000 | 50.82400 |
| C | 32.80700 | 30.46300 | 50.04700 |
| C | 33.11400 | 31.92700 | 49.53200 |
| O | 32.91300 | 32.28500 | 48.36700 |
| C | 31.45380 | 30.41880 | 50.78830 |
| C | 30.29160 | 30.69780 | 49.84450 |
| O | 29.77380 | 31.81690 | 49.76990 |
| O | 29.93900 | 29.63870 | 49.14650 |
| H | 33.70520 | 29.79340 | 51.83460 |
| H | 32.70690 | 29.90220 | 49.10470 |
| H | 31.33020 | 29.40830 | 51.20180 |
| H | 31.43020 | 31.15300 | 51.60470 |
| H | 29.10140 | 29.79500 | 48.53190 |
| N | 33.63800 | 32.82200 | 50.35700 |
| H | 33.88100 | 32.60950 | 51.31990 |
| C | 33.90300 | 32.02700 | 44.81800 |
| O | 33.71700 | 32.08700 | 43.59100 |
| N | 33.39200 | 32.93300 | 45.68900 |
| C | 32.51000 | 34.06700 | 45.26000 |
| C | 33.32800 | 35.05900 | 44.40200 |
| O | 32.77200 | 35.51900 | 43.37500 |
| C | 31.81240 | 34.70260 | 46.48210 |
| C | 31.01250 | 35.93690 | 46.09420 |
| O | 30.89010 | 33.76950 | 47.07700 |
| H | 33.47540 | 32.73340 | 46.68900 |
| H | 31.74080 | 33.67640 | 44.58010 |
| H | 32.58590 | 34.97680 | 47.22500 |
| H | 31.41500 | 33.10580 | 47.57790 |
| H | 30.50650 | 36.33560 | 46.98360 |
| H | 30.25750 | 35.67970 | 45.33820 |
| H | 31.66050 | 36.71740 | 45.67570 |
| N | 34.61300 | 35.31200 | 44.78500 |
| H | 35.05090 | 34.70520 | 45.47100 |
| C | 23.94400 | 27.94100 | 45.50200 |
| O | 23.57900 | 28.09500 | 44.33100 |
| N | 25.20400 | 28.15800 | 45.87400 |
| C | 26.22300 | 28.52000 | 44.89200 |
| C | 26.47000 | 27.38700 | 43.85800 |
| O | 26.67800 | 27.71100 | 42.68900 |
| C | 27.55000 | 28.84110 | 45.60180 |

| | | | |
|---|---|---|---|
| C | 27.49270 | 30.04670 | 46.55400 |
| O | 27.10990 | 31.15610 | 46.08420 |
| O | 27.84180 | 29.86490 | 47.77040 |
| H | 25.47130 | 28.04480 | 46.84980 |
| H | 25.87880 | 29.37920 | 44.30400 |
| H | 27.90310 | 27.96000 | 46.15660 |
| H | 28.29250 | 29.07370 | 44.82370 |
| N | 26.43600 | 26.10900 | 44.22600 |
| H | 26.16120 | 25.82240 | 45.15970 |
| C | 22.02800 | 35.96100 | 51.62100 |
| O | 21.52500 | 35.67200 | 52.73400 |
| N | 23.04900 | 35.23800 | 51.14800 |
| C | 23.74700 | 34.28100 | 52.02300 |
| C | 24.10700 | 34.76800 | 53.38500 |
| O | 23.80400 | 34.05500 | 54.34000 |
| C | 25.02000 | 33.77640 | 51.29550 |
| C | 24.74960 | 32.98580 | 50.00600 |
| C | 26.01590 | 32.45190 | 49.31220 |
| C | 26.86190 | 33.53990 | 48.64520 |
| N | 28.00640 | 32.92530 | 47.90380 |
| H | 23.50280 | 35.53160 | 50.28500 |
| H | 23.07210 | 33.43250 | 52.20360 |
| H | 25.67100 | 34.64470 | 51.09410 |
| H | 25.57010 | 33.13520 | 52.00180 |
| H | 24.10220 | 32.13030 | 50.25420 |
| H | 24.17840 | 33.59690 | 49.28680 |
| H | 26.63840 | 31.90050 | 50.03590 |
| H | 25.72580 | 31.72420 | 48.53960 |
| H | 26.26250 | 34.10810 | 47.91970 |
| H | 27.27570 | 34.24690 | 49.37730 |
| H | 28.60210 | 32.36520 | 48.56040 |
| H | 28.63660 | 33.62030 | 47.47740 |
| H | 27.64430 | 32.24720 | 47.13820 |
| N | 24.74100 | 35.92700 | 53.53200 |
| H | 24.85880 | 36.57370 | 52.75890 |
| H | 35.81630 | 29.27130 | 51.07360 |
| H | 33.84900 | 33.74890 | 50.00040 |
| H | 34.49770 | 31.23040 | 45.31340 |
| H | 35.21020 | 35.82310 | 44.14210 |
| H | 23.26540 | 27.63500 | 46.32770 |
| H | 26.53750 | 25.39870 | 43.50860 |
| H | 21.65640 | 36.76500 | 50.95420 |
| H | 24.88970 | 36.27920 | 54.47230 |

**Table S41.** Optimized structure in the forward direction from the intrinsic reaction coordinate derived from the transition state optimization for proton transfer from complex 2 to complex 3 without explicit waters.

| | | | |
|---|---|---|---|
| C | 35.07590 | 29.61810 | 50.32390 |
| O | 35.29100 | 29.74080 | 49.11940 |
| N | 33.82760 | 29.84070 | 50.82330 |
| C | 32.80470 | 30.46380 | 50.05000 |
| C | 33.11290 | 31.92590 | 49.53220 |
| O | 32.91430 | 32.28590 | 48.36660 |
| C | 31.47110 | 30.48280 | 50.82100 |
| C | 30.28000 | 30.87240 | 49.93250 |
| O | 29.57780 | 31.86430 | 50.26140 |
| O | 30.09440 | 30.12740 | 48.91100 |
| H | 33.70480 | 29.79030 | 51.83330 |
| H | 32.66280 | 29.89870 | 49.11740 |
| H | 31.28050 | 29.46420 | 51.19350 |
| H | 31.52360 | 31.16810 | 51.67840 |
| H | 28.87560 | 29.93220 | 48.31760 |
| N | 33.63780 | 32.82180 | 50.35660 |
| H | 33.87120 | 32.61120 | 51.32230 |
| C | 33.90280 | 32.02720 | 44.81830 |
| O | 33.71690 | 32.08710 | 43.59090 |
| N | 33.39200 | 32.93280 | 45.68930 |
| C | 32.51010 | 34.06680 | 45.26020 |
| C | 33.32800 | 35.05880 | 44.40210 |
| O | 32.77200 | 35.51900 | 43.37490 |
| C | 31.81960 | 34.70380 | 46.48640 |
| C | 30.99440 | 35.92090 | 46.09900 |
| O | 30.92340 | 33.76030 | 47.10580 |
| H | 33.47730 | 32.73210 | 46.68930 |
| H | 31.73870 | 33.67640 | 44.58290 |
| H | 32.59990 | 34.99670 | 47.21500 |
| H | 31.47280 | 33.09620 | 47.58370 |
| H | 30.50290 | 36.32730 | 46.99320 |
| H | 30.22610 | 35.64150 | 45.36450 |
| H | 31.62230 | 36.70310 | 45.65420 |
| N | 34.61290 | 35.31190 | 44.78500 |
| H | 35.05110 | 34.70490 | 45.47070 |
| C | 23.94250 | 27.94050 | 45.50150 |
| O | 23.57960 | 28.09560 | 44.33070 |
| N | 25.20300 | 28.15830 | 45.87450 |
| C | 26.22070 | 28.51900 | 44.89180 |
| C | 26.47030 | 27.38550 | 43.85710 |
| O | 26.67820 | 27.71180 | 42.68940 |

| | | | |
|---|---|---|---|
| C | 27.58230 | 28.72870 | 45.57110 |
| C | 27.71310 | 29.83830 | 46.60110 |
| O | 27.54560 | 31.02990 | 46.30030 |
| O | 28.07110 | 29.39630 | 47.78660 |
| H | 25.47200 | 28.02160 | 46.84680 |
| H | 25.90140 | 29.39430 | 44.31490 |
| H | 27.90380 | 27.78840 | 46.03840 |
| H | 28.30550 | 28.97070 | 44.77690 |
| N | 26.43600 | 26.10860 | 44.22600 |
| H | 26.16510 | 25.81840 | 45.15970 |
| C | 22.02800 | 35.96110 | 51.62110 |
| O | 21.52490 | 35.67200 | 52.73390 |
| N | 23.04890 | 35.23790 | 51.14790 |
| C | 23.74680 | 34.28100 | 52.02300 |
| C | 24.10710 | 34.76800 | 53.38510 |
| O | 23.80390 | 34.05500 | 54.33990 |
| C | 25.02990 | 33.78670 | 51.30700 |
| C | 24.79350 | 33.02030 | 49.99710 |
| C | 26.08410 | 32.47300 | 49.35980 |
| C | 26.98310 | 33.55200 | 48.74770 |
| N | 28.21970 | 32.92490 | 48.18670 |
| H | 23.50280 | 35.53190 | 50.28510 |
| H | 23.07540 | 33.42920 | 52.20050 |
| H | 25.68590 | 34.65730 | 51.13560 |
| H | 25.56520 | 33.13250 | 52.01270 |
| H | 24.12080 | 32.17350 | 50.20410 |
| H | 24.26740 | 33.65210 | 49.26160 |
| H | 26.66090 | 31.91150 | 50.11320 |
| H | 25.81870 | 31.75530 | 48.56750 |
| H | 26.46430 | 34.08140 | 47.93620 |
| H | 27.29980 | 34.29040 | 49.49630 |
| H | 28.75370 | 32.43540 | 48.98080 |
| H | 28.87770 | 33.59630 | 47.75790 |
| H | 27.97150 | 32.20170 | 47.45270 |
| N | 24.74090 | 35.92700 | 53.53200 |
| H | 24.86340 | 36.57270 | 52.75880 |
| H | 35.81700 | 29.27110 | 51.07310 |
| H | 33.84210 | 33.75100 | 50.00270 |
| H | 34.49680 | 31.23030 | 45.31370 |
| H | 35.20990 | 35.82340 | 44.14210 |
| H | 23.26500 | 27.63020 | 46.32600 |
| H | 26.55530 | 25.39860 | 43.51110 |
| H | 21.65690 | 36.76570 | 50.95460 |
| H | 24.89500 | 36.27680 | 54.47230 |

**Table S42.** Optimized structure in the backward direction from the intrinsic reaction coordinate derived from the transition state optimization for proton transfer from complex 2 to complex 3 without explicit waters.

| | | | |
|---|---|---|---|
| C | 35.07610 | 29.61790 | 50.32410 |
| O | 35.28900 | 29.74120 | 49.12060 |
| N | 33.82440 | 29.83930 | 50.82470 |
| C | 32.80930 | 30.46220 | 50.04400 |
| C | 33.11510 | 31.92810 | 49.53180 |
| O | 32.91170 | 32.28410 | 48.36740 |
| C | 31.48030 | 30.45950 | 50.82380 |
| C | 30.27940 | 30.77320 | 49.94610 |
| O | 29.54400 | 31.74260 | 50.17720 |
| O | 30.12620 | 29.91090 | 48.97140 |
| H | 33.70350 | 29.79730 | 51.83530 |
| H | 32.67960 | 29.89750 | 49.10990 |
| H | 31.32750 | 29.44660 | 51.22700 |
| H | 31.50940 | 31.17330 | 51.65770 |
| H | 29.18180 | 29.94510 | 48.46600 |
| N | 33.63820 | 32.82220 | 50.35740 |
| H | 33.88320 | 32.60700 | 51.31920 |
| C | 33.90320 | 32.02680 | 44.81770 |
| O | 33.71710 | 32.08690 | 43.59110 |
| N | 33.39200 | 32.93320 | 45.68870 |
| C | 32.50990 | 34.06720 | 45.25980 |
| C | 33.32800 | 35.05920 | 44.40190 |
| O | 32.77200 | 35.51900 | 43.37510 |
| C | 31.81820 | 34.70530 | 46.48430 |
| C | 30.99320 | 35.92200 | 46.09580 |
| O | 30.92100 | 33.76260 | 47.10420 |
| H | 33.48390 | 32.73700 | 46.68820 |
| H | 31.73910 | 33.67620 | 44.58210 |
| H | 32.59780 | 34.99890 | 47.21360 |
| H | 31.47200 | 33.09600 | 47.57440 |
| H | 30.50080 | 36.32820 | 46.98950 |
| H | 30.22570 | 35.64230 | 45.36070 |
| H | 31.62150 | 36.70430 | 45.65160 |
| N | 34.61310 | 35.31210 | 44.78500 |
| H | 35.05200 | 34.70330 | 45.46850 |
| C | 23.94550 | 27.94150 | 45.50250 |
| O | 23.57840 | 28.09440 | 44.33130 |
| N | 25.20500 | 28.15770 | 45.87350 |
| C | 26.22530 | 28.52100 | 44.89220 |

| | | | |
|---|---|---|---|
| C | 26.46970 | 27.38850 | 43.85890 |
| O | 26.67780 | 27.71020 | 42.68860 |
| C | 27.57490 | 28.77880 | 45.58840 |
| C | 27.63180 | 29.92990 | 46.60400 |
| O | 27.38450 | 31.10090 | 46.20700 |
| O | 27.96920 | 29.61470 | 47.80010 |
| H | 25.47280 | 28.04430 | 46.84920 |
| H | 25.89630 | 29.39100 | 44.31200 |
| H | 27.90860 | 27.85780 | 46.08720 |
| H | 28.30380 | 29.01620 | 44.79850 |
| N | 26.43600 | 26.10940 | 44.22600 |
| H | 26.16780 | 25.81970 | 45.16060 |
| C | 22.02800 | 35.96090 | 51.62090 |
| O | 21.52510 | 35.67200 | 52.73410 |
| N | 23.04910 | 35.23810 | 51.14810 |
| C | 23.74720 | 34.28100 | 52.02300 |
| C | 24.10690 | 34.76800 | 53.38490 |
| O | 23.80410 | 34.05500 | 54.34010 |
| C | 25.02800 | 33.78480 | 51.30360 |
| C | 24.78140 | 33.01710 | 49.99610 |
| C | 26.06210 | 32.46300 | 49.34660 |
| C | 26.95170 | 33.53280 | 48.70630 |
| N | 28.14590 | 32.88480 | 48.07980 |
| H | 23.50080 | 35.52890 | 50.28310 |
| H | 23.07550 | 33.42940 | 52.20060 |
| H | 25.68480 | 34.65430 | 51.12830 |
| H | 25.56450 | 33.13020 | 52.00800 |
| H | 24.10720 | 32.17320 | 50.20990 |
| H | 24.25150 | 33.64960 | 49.26400 |
| H | 26.65040 | 31.90790 | 50.09580 |
| H | 25.78920 | 31.73610 | 48.56620 |
| H | 26.41010 | 34.07590 | 47.91950 |
| H | 27.31240 | 34.26200 | 49.44500 |
| H | 28.67380 | 32.36550 | 48.82700 |
| H | 28.80990 | 33.55240 | 47.65660 |
| H | 27.84020 | 32.16890 | 47.32350 |
| N | 24.74110 | 35.92700 | 53.53200 |
| H | 24.85870 | 36.57420 | 52.75920 |
| H | 35.81720 | 29.27140 | 51.07310 |
| H | 33.84730 | 33.75060 | 50.00320 |
| H | 34.49810 | 31.23070 | 45.31360 |
| H | 35.20990 | 35.82270 | 44.14120 |
| H | 23.26780 | 27.63510 | 46.32890 |
| H | 26.54840 | 25.40080 | 43.50880 |
| H | 21.65610 | 36.76460 | 50.95390 |
| H | 24.89060 | 36.27840 | 54.47250 |

**Table S43.** Relaxed coordinates of the Asparagine mutant.

| | | | |
|---|---|---|---|
| C | 34.16400 | 35.32800 | 47.73900 |
| O | 34.37300 | 35.13800 | 46.49800 |
| N | 33.27400 | 34.56400 | 48.36300 |
| C | 32.37400 | 33.54700 | 47.83100 |
| C | 31.56500 | 34.13100 | 46.69700 |
| O | 31.29200 | 33.51100 | 45.64200 |
| C | 31.48830 | 32.94060 | 48.93320 |
| C | 30.49190 | 31.96140 | 48.30420 |
| N | 30.94050 | 30.73260 | 48.02900 |
| O | 29.33880 | 32.35830 | 48.03610 |
| H | 33.17260 | 34.73570 | 49.36440 |
| H | 32.97350 | 32.74640 | 47.36850 |
| H | 32.13260 | 32.44040 | 49.67070 |
| H | 30.90830 | 33.72650 | 49.43510 |
| H | 30.37270 | 30.02760 | 47.50780 |
| H | 31.89620 | 30.47540 | 48.25280 |
| N | 31.19000 | 35.39400 | 46.84300 |
| H | 31.43430 | 35.94540 | 47.65840 |
| C | 33.13900 | 33.20200 | 42.26000 |
| O | 33.28000 | 32.46600 | 41.26400 |
| N | 31.90000 | 33.53500 | 42.72600 |
| C | 30.62400 | 33.09900 | 41.98400 |
| C | 30.59000 | 33.89900 | 40.66500 |
| O | 30.13700 | 33.42700 | 39.68600 |
| C | 29.38420 | 33.33110 | 42.86680 |
| C | 28.09090 | 33.01540 | 42.12880 |
| O | 29.42330 | 32.47900 | 44.02290 |
| H | 31.81360 | 33.94330 | 43.65870 |
| H | 30.71400 | 32.04070 | 41.71140 |
| H | 29.38150 | 34.39100 | 43.18890 |
| H | 30.12020 | 32.81240 | 44.64240 |
| H | 28.07870 | 31.96050 | 41.82230 |
| H | 27.97550 | 33.63700 | 41.23280 |
| H | 27.23840 | 33.19730 | 42.79740 |
| N | 31.23400 | 35.10100 | 40.60500 |
| H | 31.76250 | 35.47450 | 41.38390 |
| C | 25.45200 | 25.96200 | 45.80200 |
| O | 24.99200 | 25.97400 | 44.64800 |
| N | 26.77100 | 26.02300 | 46.06800 |
| C | 27.87900 | 25.94800 | 45.01400 |
| C | 28.12200 | 24.48400 | 44.51500 |
| O | 28.39800 | 24.25300 | 43.35800 |
| C | 29.14910 | 26.60200 | 45.54590 |

| | | | |
|---|---|---|---|
| C | 29.02610 | 28.09540 | 45.91510 |
| O | 29.89860 | 28.55560 | 46.68770 |
| O | 28.05000 | 28.76300 | 45.41330 |
| H | 27.05590 | 26.09660 | 47.04250 |
| H | 27.50640 | 26.48550 | 44.13420 |
| H | 29.53340 | 26.06440 | 46.42550 |
| H | 29.92620 | 26.52440 | 44.76920 |
| N | 27.88500 | 23.43300 | 45.35200 |
| H | 27.56060 | 23.55220 | 46.30510 |
| C | 20.19600 | 35.11900 | 46.20900 |
| O | 19.76000 | 35.39100 | 47.31500 |
| N | 21.46900 | 34.65800 | 45.96700 |
| C | 22.43600 | 34.38200 | 47.08200 |
| C | 22.62100 | 35.57700 | 48.07800 |
| O | 22.54900 | 35.51400 | 49.30900 |
| C | 23.77690 | 33.95380 | 46.45030 |
| C | 24.89130 | 33.68970 | 47.46970 |
| C | 26.24520 | 33.30250 | 46.84600 |
| C | 26.29400 | 31.87040 | 46.30590 |
| N | 27.69930 | 31.46110 | 45.98400 |
| H | 21.74210 | 34.42460 | 45.01610 |
| H | 22.03280 | 33.56550 | 47.70040 |
| H | 23.59270 | 33.05210 | 45.84360 |
| H | 24.10760 | 34.74540 | 45.75500 |
| H | 25.04100 | 34.59640 | 48.07630 |
| H | 24.57270 | 32.90410 | 48.17440 |
| H | 26.51060 | 34.01830 | 46.04880 |
| H | 27.02020 | 33.39810 | 47.62200 |
| H | 25.93240 | 31.15940 | 47.05990 |
| H | 25.68950 | 31.73120 | 45.39990 |
| H | 28.32960 | 31.68290 | 46.79870 |
| H | 28.10460 | 31.96320 | 45.16160 |
| H | 27.80600 | 30.42250 | 45.78460 |
| N | 22.81500 | 36.74500 | 47.47800 |
| H | 22.72830 | 36.84970 | 46.47210 |
| O | 26.04800 | 28.80300 | 47.33100 |
| H | 26.76110 | 28.68370 | 46.65920 |
| H | 25.25430 | 28.82460 | 46.76480 |
| O | 25.52300 | 28.90400 | 44.13600 |
| H | 26.46270 | 28.76350 | 44.39320 |
| H | 25.17640 | 27.99100 | 44.06690 |
| H | 34.68230 | 36.07640 | 48.36600 |
| H | 30.64990 | 35.83640 | 46.10640 |
| H | 33.97550 | 33.61690 | 42.85860 |
| H | 31.29740 | 35.55080 | 39.69870 |
| H | 24.81870 | 25.94960 | 46.71270 |
| H | 27.98190 | 22.49700 | 44.97410 |
| H | 19.60560 | 35.22520 | 45.27080 |
| H | 22.84430 | 37.58690 | 48.04450 |
